# An Autonomous Molecular Bioluminescent Reporter (AMBER) for voltage imaging in freely moving animals

**DOI:** 10.1101/845198

**Authors:** Prasanna Srinivasan, Nicole M Griffin, Pradeep Joshi, Dhananjay Thakur, Alex Nguyen-Le, Sean McCotter, Akshar Jain, Mitra Saeidi, Prajakta Kulkarni, Jaclyn T. Eisdorfer, Joel Rothman, Craig Montell, Luke Theogarajan

## Abstract

1.

Genetically encoded reporters have greatly increased our understanding of biology, especially in neuroscience. While fluorescent reporters have been widely used, photostability and phototoxicity have hindered their use in long-term experiments. Bioluminescence overcomes some of these challenges but requires the addition of an exogenous luciferin limiting its use. Using a modular approach we have engineered Autonomous Molecular BioluminEscent Reporter (AMBER), an indicator of membrane potential. Unlike other luciferase-luciferin bioluminescent systems, AMBER encodes the genes to express both the luciferase and luciferin. AMBER is a voltage-gated luciferase coupling the functionalities of the *Ciona* voltage-sensing domain (VSD) and bacterial luciferase, luxAB. When AMBER is co-expressed with the luciferin producing genes it reversibly switches the bioluminescent intensity as a function of membrane potential. Using biophysical and biochemical methods we show that AMBER modulates its enzymatic activity as a function of the membrane potential. AMBER shows several-fold increase in the luminescent (ΔL/L) signal upon switching from the off to on state when the cell is depolarized. *In vivo* expression of AMBER in *C. elegans* allowed detecting pharyngeal pumping action and mechanosensory neural activity from multiple worms simultaneously. AMBER reports neural activity of multiple animals at the same time and can be used in social behavior assays to elucidate the role of membrane potential underlying behavior.

**Significance Statement:** There have been many exciting advances in the development of genetically encoded voltage indicators to monitor intracelluar voltage changes. Most sensors employ fluorescence, which requires external light, potentially causing photobleaching or overheating. Consequently, there has been interest in developing luminescence reporters. However, they require addition of an exogenous substrate to produce light intracellularly. Here, we engineered a genetically encoded bioluminescent voltage indicator, AMBER, which unlike other bioluminescent activity indicators, does not require addition of an exogenous substrate. AMBER allows a large differential signal, a high signal-to-noise ratio, and causes minimal metabolic demand on cells. We used AMBER to record voltage activity in freely-moving C. *elegans*, demonstrating that AMBER is a important new tool for monitoring neuronal activity during social behavior.

## 3. Introduction

Genetically encoded optical reporters have gained prominence for reporting protein-protein interactions, gene expression, and cellular signaling. Most reporters provide an optical readout of the signal of interest via the excitation of a fluorescent protein [1]. Application of molecular engineering to detect, record and modulate neural signatures is a major area underpinning several neurotechnology initiatives. Over the last decade, there has been an exponential growth in the development of genetically engineered molecular neural probes owing to their ability to precisely target tissues of interest. A wide range of biomolecular sensors currently exists for detecting changes in voltage [2,3], and the levels of calcium [2,4,5], potassium [6,7] and neurotransmitters [8-10]. While fluorescent reporters have been the mainstay, it suffers from photobleaching and phototoxicity affecting their long-term use [11]. Near-infrared/red indicators [4,12,13] and two-photon excitation [14,15] overcome some of these limitations, but the development of efficient indicators of neural activity remains an issue.

Intracellular calcium (Ca^2+^) concentration and membrane potential are the two well-characterized proxies of neuronal activity. Several Genetically Encoded Ca^2+^ Indicators (GECIs) [2, 4-5, 16] were developed over the last two decades with GCaMP variants [17] being most widely employed in the neuroscience community. However, the relationship between Ca^2+^ signals and neuronal activity can vary among different neuronal types and may become completely uncoupled [18,19]. Despite the great utility of GECIs, they have some potential limitations. For instance, several organelles (endoplasmic reticulum and mitochondria) and regulatory proteins contribute to the spatially varying Ca^2+^ compartmentalization [20, 21] in neurons. Overexpression of GECIs in these neurons can cause these Ca2+-responsive probes to contribute to Ca2+ buffering, thereby affecting neurophysiological conditions [21]. Despite its high signal-to-noise ratio (SNR), GECIs cannot report subthreshold or hyperpolarizing neuronal events [22,23] compared to fluorescent Genetically Encoded Voltage Indicators (GEVIs) [24]. Recently, neural spikes of hundreds of neurons were recorded for several minutes using a fluorescent hybrid GEVI [25], but the necessity for complexing with a synthetic dye can be a limitation for long-term use. Deciphering the information processing in circuits with a large population of neurons necessitates conducting experiments for a long duration demanding molecular reporters with exceptional photostability.

Recently bioluminescence has garnered attention as an alternative to overcome some of the limitations of fluorescent GEVIs and GECIs [27]. Bioluminescent imaging does not require any external illumination, and light emission is achieved as a by-product of a biochemical reaction – oxidation of a substrate (luciferin) catalysed by an enzyme (luciferase). Commonly used light-generating luciferase from fireflies (*Lampyridae*) and marine organisms such as *Aequoria, Renilla, Gaussia, Metridia*, and *Vargula* [28-32] require exogenous addition of luciferin since the biosynthetic pathways for *in situ* luciferin production have not yet been identified. There are two exceptions – the genes coding for the fungal luciferin, 3-hydroxyhispidin [33] and the bacterial luciferin, a molecular complex of fatty aldehyde (Myristyl aldehyde, CH_3_(CH_2_)_12_CHO) and reduced flavin mononucleotide, FMNH_2_ [34]. In a *tour-de-force*, Kotlobay et al. [35] discovered a set of genes required for expressing both the fungal luciferase and luciferin. However, the fungal bioluminescence system requires a total of seven genes assembled from multiple organisms, which can be challenging to express efficiently in eukaryotic systems. In contrast, the bacterial *lux* operon genes are encoded in a single polycistronic mRNA [36] and can enable enhanced bioluminescence [37].

The *lux* operon, unlike other bioluminescent systems, consists of a series of six genes *(luxCDABE* and the flavin oxidoreductase gene, *frp*) synergistically combining to produce both the luciferase and a luciferin-generating secondary protein complex (See Figure S1). The luciferin synthesizing protein complex recycles the products (Myristic acid, CH_3_(CH_2_)_12_COOH and flavin mononucleotide, FMN) and the intermediates of the light reaction endogenously to light generating substrates (CH_3_(CH_2_)_12_CHO and FMNH_2_) using metabolic pathways [36]. Fortuitously, CH_3_(CH_2_)_12_CHO is not freely available in large quantities in eukaryotes [38], ensuring low background activation. There were concerns about apparent cytotoxic effects of aldehyde compounds in eukaryotes [39] but the concentration of CH_3_(CH_2_)_12_CHO synthesized using *lux* operon expression does not seem to attain the toxic dose affecting cell viability [38]. Recent work also confirmed there is no cytotoxic effect when the lux cassette is expressed in mammalian cells [40].

In this paper, we describe a new type of LuVI (Luminescent Voltage Indicator) named AMBER for observing neuronal activity in freely moving animals. AMBER uses the functionalities of the ascidian *Ciona intestinalis* voltage-sensing domain (VSD) [41], bacterial luciferase, luxAB [38] (derived from *P. luminescens*), and a fluorescent protein, YPet [42]. AMBER leverages the molecular architecture of a previously developed GEVI, VSFP2.1 [43]. The rationale behind this choice stems from the efficient coupling between the conformational change of the VSD and the Cerulean/Citrine FRET pair of VSFP2.1 that operates within the physiological voltage range. AMBER provides an unprecedentedly large dynamic range in the optical readout and an increased efficiency of the biophotonic emission. Unlike other voltage probes reported to date, AMBER undergoes an increase in the enzymatic activity upon depolarization. The overall light emission is 10X times brighter than the initial resting signal, which is greater than the signal change observed in the LOTUS-V [44], the first LuVI developed using the deep shrimp luciferase [45]. We expressed the optimized first generation of AMBER in C. *elegans* pharyngeal muscles and mechanosensory neurons and demonstrate that it functions *in vivo*. AMBER allowed recording neural signatures in multiple freely moving animals in different directions simultaneously, which is not possible using fluorescent imaging approaches.

## 4. Results

### 4.1 Optimizing engineered AMBER Constructs

We postulated voltage sensing ability can be conferred to the *lux*-based LuVIs by replacing the fluorescent FRET donor, Cerulean, of VSFP2.1 [43] with the mammalian codon-optimized synthetic luciferase, enhanced luxAB (‘eluxAB’) [46]. This replacement resulted in the protein construct VSD-eluxAB-YPet (*See the Supporting Information, SI under Section 1*). Previous reports on BRET performance show that the acceptor position could influence the brightness of the probe. We, therefore, swapped the position of eluxAB and YPet (designated VSD-YPet-eluxAB) to compare its performance with the VSD-eluxAB-YPet.

FMNH_2_ is the rate-limiting substrate in the bacterial bioluminescent reaction [47]. Fusing FRP to luxAB would enable direct transfer of FMNH_2_ to luxAB from the FRP-FMNH_2_ complex. This direct transfer minimizes FMNH_2_ oxidation via the dark pathway, thereby significantly increasing the bioluminescent quantum yield [48,49] of the bioluminescent reaction. We tested the hypothesis that placing FRP in the vicinity of the membrane-targeted eluxAB would increase bioluminescent light intensity. We therefore fused the FRP domain to the N-termini of both VSD-YPet-eluxAB and VSD-eluxAB-YPet to obtain FRP-VSD-YPet-eluxAB and FRP-VSD-eluxAB-YPet, respectively (*See the SI under Section 1* and Figure 1).

**Figure 1:**
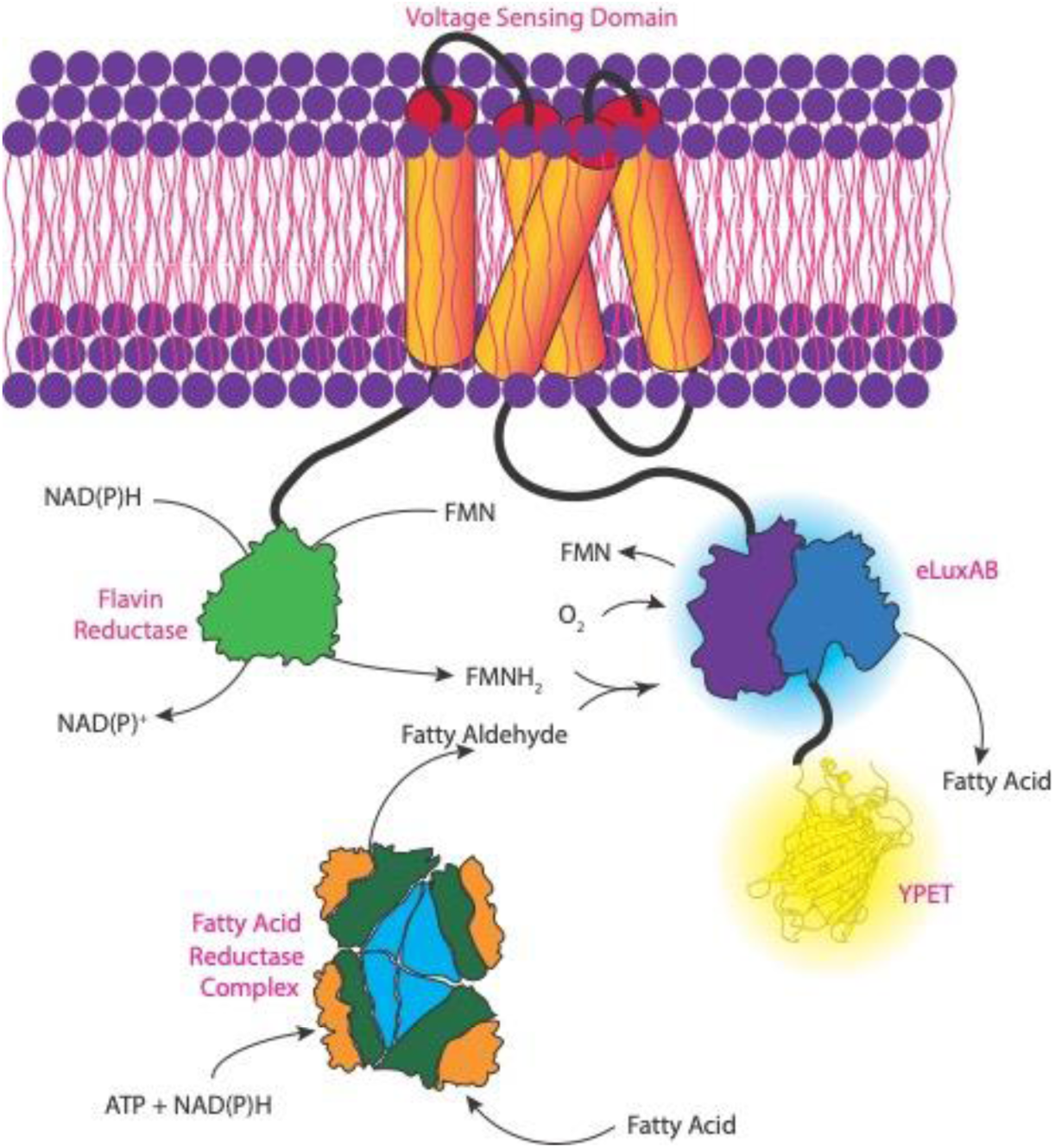
Schematic representation illustrating the molecular architecture of AMBER. AMBER is a plasma membrane resident *Ciona* voltage-sensing domain (Ci-VSD) whose N and C termini are fused to Flavin reductase phosphate (FRP) and a BRET pair comprising bright bacterial luciferase bioluminescent donor (eluxAB) and a yellow fluorescent protein acceptor (YPet). An increase in membrane potential increases the probability of photon emission (λ_max_ ∼ 490nm) by the luciferase domain that catalyses the reaction between a reduced flavin (FMNH_2_), fatty aldehyde (synthesized by the fatty acid reductase complex) and molecular oxygen. The positioning of the Flavin reductase phosphate in the N-terminus reduces the auto oxidation of the FMNH_2_ by enabling direct shuttling between the luciferase and the flavin reductase.

Plasmid DNAs encoding the four principal engineered protein constructs − VSD-eluxAB-YPet, VSD-YPet-eluxAB, FRP-VSD-eluxAB-YPet, and FRP-VSD-YPet-eluxAB were co-expressed with their respective substrate proteins in HEK293 cells (*See the SI under Section 2).* Bioluminescent images were recorded using a custom-built imaging set up (*See the SI under Section 3*) in a dark room. The recorded images were subsequently processed using the ImageJ software (*See the SI under Section 10*).

Efficacy of all the engineered constructs were tested by imaging cells before and after depolarization by KCl addition (50 mM final concentration). VSD-eluxAB-YPet performed better than VSD-YPet-eluxAB in modulating the eluxAB activity (Figure 2a). Surprisingly FRP-VSD-YPet-eluxAB performed the poorest of the constructs tested, indicating the positioning of eluxAB and YPet affects the light modulation more than the proximity to FRP. However, as expected FRP-VSD-eluxAB-YPet performed the best and this result confirms that placing the eluxAB and FRP domains in close apposition increases the propensity of direct FMNH_2_ transfer from FRP to eluxAB. The signal-to-noise ratio (SNR) of the images (Figure 2b) obtained for the various constructs after KCl addition indicates that the FRP-VSD-eluxAB-YPet performed the best among all the engineered constructs tested (Figure S2). Light due to endogenously produced substrates (Figure S3) is very dim – about 10 times smaller than the maximum achieved signal even after integrating three times as long. In contrast, untransfected cells showed no detectable intensity change after KCl challenge (Figure S4).

**Figure 2:**
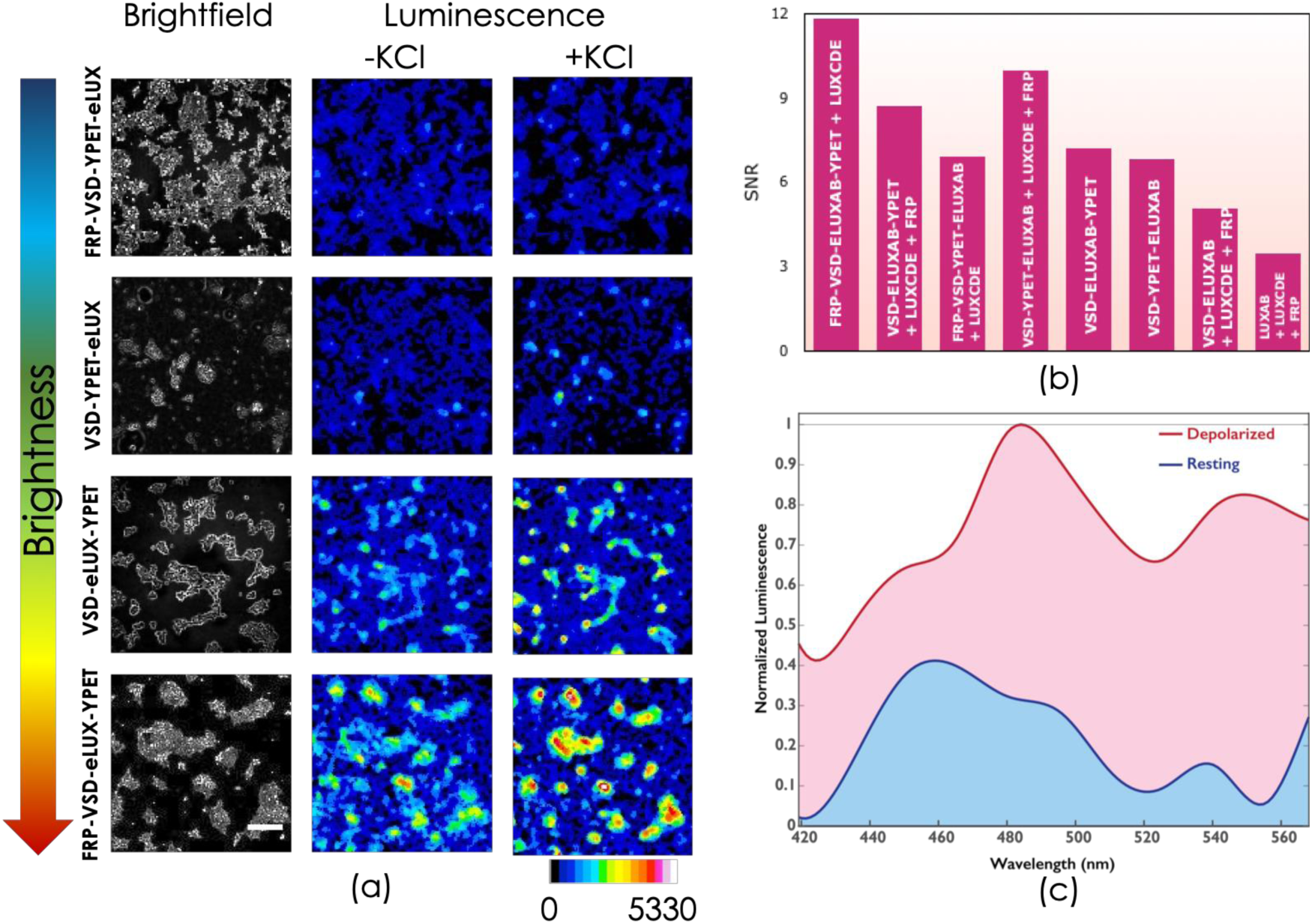
(a) In vitro bioluminescent performance before and after KCl challenge of various engineered protein constructs expressed in HEK293 cells. FRP-VSD-eluxAB-YPet is the best candidate (Maximum fractional luminescence, (ΔL/L)_max_ ∼ 28; Average fractional luminescence, (ΔL/L)_avg_ ∼ 0.72; Information entropy ratio, IR ∼ 85%; n=53 cells; p<0.00001). See supplementary tables S2 and S3 for the information entropy and t-test statistics of all engineered protein constructs; (b) Signal-to-noise ratio of various engineered constructs. Performance of the majority of the constructs meet Rose criterion (SNR > 5) suggesting the signal is sufficiently higher than the background noise. (c) Emission spectra of HEK293 cells expressing the bright construct before and after KCl challenge. Photon counts for each spectral wavelength show the mean counts obtained from n=8 trials. The scale bar of the micrograph corresponds to 200μm

### 4.2 Plate reader assay

An increase in the bioluminescent signal by several-fold after KCl addition precludes that the majority of the emitted energy is due to the resonance energy transfer. We, therefore, performed a more quantitative assay by recording the emission spectra using a plate reader (*See the SI under Section 4*). Luminescence spectra of HEK293 cells expressing the bright construct (FRP-VSD-eluxAB-YPet) were recorded before and after KCl addition. Figure 2c provides direct evidence of enzymatic modulation after KCl addition, as there were increases in both the eluxAB donor peak (∼ 490 nm) and the YPet acceptor peak (∼ 530nm). In comparison, the number of photons detected before KCl addition were smaller, indicating a very low basal activity of eluxAB.

### 4.3 Dark Mutant

Additional evidence to confirm the enzymatic switching of AMBER upon depolarization was obtained by mutating residues G65T and G67A within the YPet chromophore of FRP-VSD-eluxAB-YPet [50] (*See the SI under Section 1*). These mutations (creating the ‘dark mutant’), greatly diminish the resonant energy transfer at the YPet wavelength. Bioluminescent imaging of HEK293 cells expressing the dark mutant showed an increase in the intensity upon depolarization using KCl (Figure 3). Plate reader experiments using the dark mutant showed an increase in the spectral intensity at 490 nm upon depolarization suggesting modulation of the enzymatic activity as the dominant mode of the light emission (*See Section 4* and Figure S5 *in the SI*).

**Figure 3:**
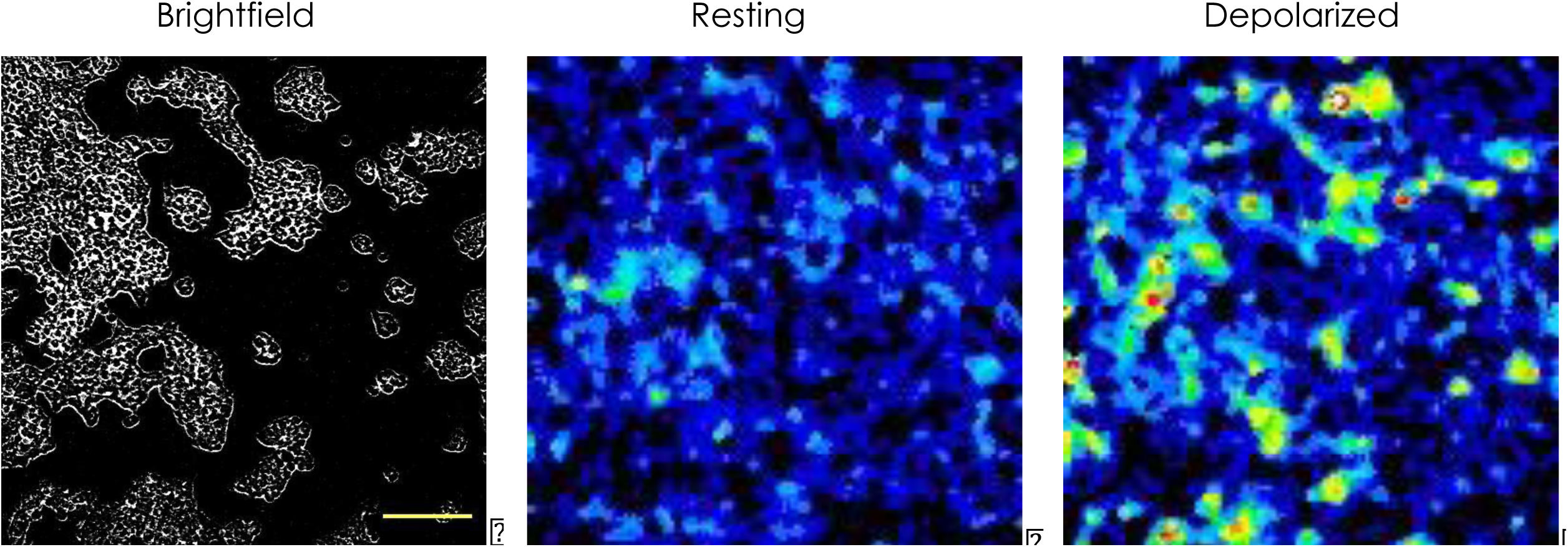
Bioluminescent emission of HEK293 cells expressing the dark mutant probe before and after KCl challenge. The data suggests the structural interaction between YPet and eluxAB domains is the primary driver for the modulation of activity. BRET seems to play only a minor role in the transduction of the membrane potential into photon count. (Maximum fractional luminescence, (ΔL/L)_max_ ∼76; Average fractional luminescence, (ΔL/L)_avg_ ∼1.2; Average information entropy ∼73.2%; n=83 cells; p<0.00001). The scale bar corresponds to 200*μ*m.

### 4.4 NADH Assay

Tracking NADH production provides indirect evidence for eluxAB enzymatic activity since NADH production increases with the drop in cellular O_2_ concentration. NADH is used both for regenerating FMNH_2_ from FMN catalyzed by FRP and for the fatty aldehyde synthesis from the fatty acid via the luxCDE complex. While the reduced form (NADH) is fluorescent, the oxidized form (NAD^+^) is not. We monitored endogenous NADH fluorescence before and after KCl addition, which showed an increase in the NADH fluorescence after KCl addition (*See Sections 3, 10 and* Figure S6 *in the SI*). While the steady-state NADH fluorescence before KCl addition is not large enough to be detected, a step increase after KCl addition invariably points to the drop in O_2_ concentration. Cytosolic NADH is produced at detectable levels within a few seconds after O_2_ consumption during the light reaction (*See* Figure S6 *in the SI*), and similar observations have been reported earlier [51]. Thus, we confirmed the causal link between depolarization and O_2_ consumption by the light reaction evidenced by the increase in the NADH fluorescence, strongly suggesting a voltage induced enzymatic-switching mechanism. This result also suggests that the steady-state aldehyde concentration in cells is sufficient to carry out the light reaction for a long duration (at least for a few tens of minutes) since an increase in fluorescence indicates that the rate of NADH production due to an O_2_ drop is faster than the rate of NADH consumption for the aldehyde synthesis.

### 4.5 Voltage switching characterization

We performed electrophysiology experiments to determine the voltage-dependent characteristics of the probe. We transiently co-expressed FRP-VSD-eluxAB-YPet and luxCDE in HEK293 cells to record single-cell bioluminescence under voltage clamp. Results from the patch-clamp experiments are shown in Figure 4a (*See section 5 in the SI*). The half maximal voltage, V_1/2_ ∼ −30 mV is within the physiological range of neuronal action potentials similar to other GEVIs [52-54]. Increase in the bioluminescence of HEK293 cells expressing two different probe constructs – FRP-VSD-eluxAB-YPet and VSD-YPet-eluxAB (with their respective substrates) was recorded by titrating with incremental amounts of KCl (Figure 4b). The V_1/2_ deduced applying Nernstian relation to the KCl titration data agrees closely with the value obtained via patch-clamp recordings. Interestingly, V_1/2_ is more positive compared to that of VSFP2.1 (V_1/2_ ∼ −70 mV) from which it was derived. Moreover V_1/2_ agrees closely with that of VSFP2.3, which is derived by modifying the linker length between the fluorescent donor and the VSD. This analysis suggests a possible structural coupling between the eluxAB and the VSD domains during activation. Additionally, V_1/2_ shifts towards the positive direction for FRP-VSD-eluxAB-YPet compared to VSD-YPet-eluxAB. We do not know if the BRET pair interacts with the VSD domain and whether its rearrangement caused the voltage shift. Earlier work showed evidence for an interaction between the GFP derived fluorescent protein and the VSD domain, but the mechanism of the fluorescence modulation by the VSD is not fully understood [52]. We speculate that the interaction between the YPet and the VSD domains could account for the observed shift.

**Figure 4:**
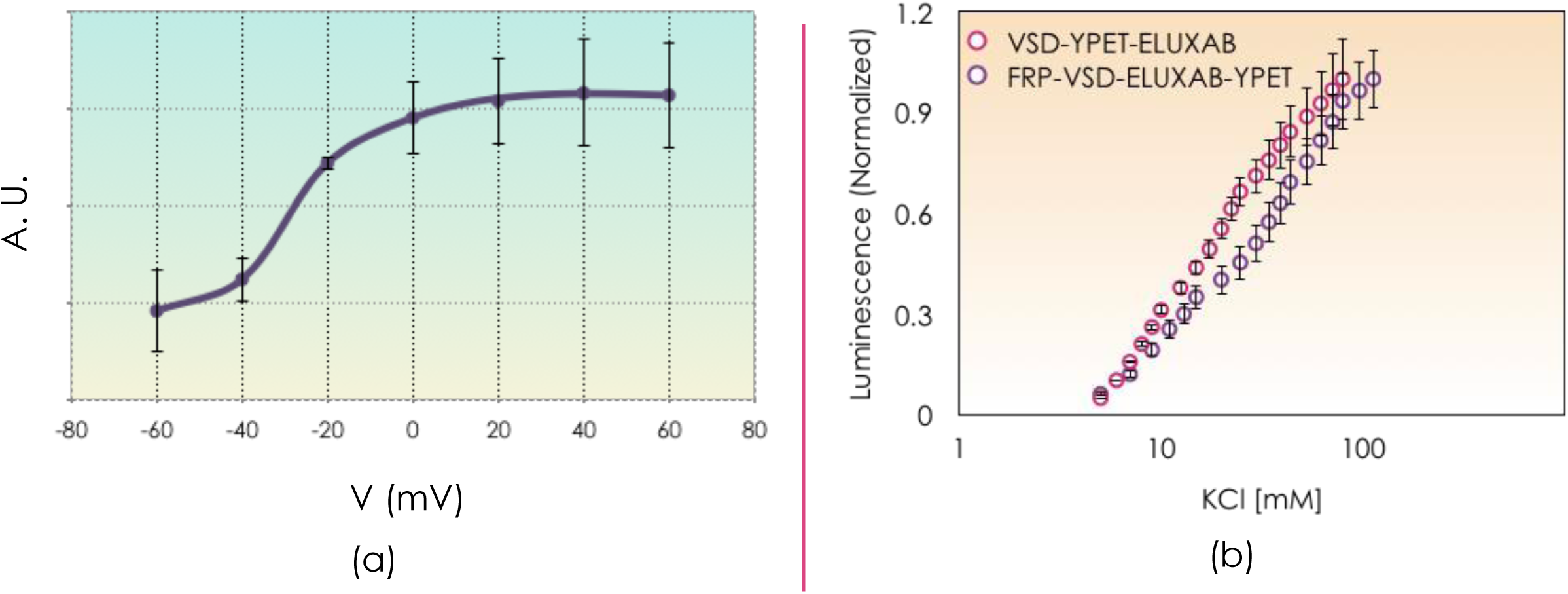
(a) Representative whole cell voltage clamp recording from a HEK293 cell expressing the bright candidate and its substrate. Recordings from six independent repeats varying the membrane potential between −60mV and 60mV at increments of 20mV were shown. The intensity of the light emitted under electrical stimulation follows the classical Boltzmann distribution with V_1/2_ ∼ −30mV. Paired t-test statistic between −60mV (off state) and +20mV (on state) confirms statistically significant difference between their bioluminescent intensities (*α* =0.05, p < 0.02) (b) Normalized bioluminescent response of the top two high performing constructs under KCl titration. The semi-log plot shows an order of magnitude change in the extracellular KCl concentration is necessary to achieve the maximum intensity for the chosen constructs.

### 4.6 Chemogenetic activation

Since an increase in extracellular K^+^ affects many cellular signaling processes, we wanted to independently verify if enzymatic switching could be achieved using a different chemical activator under physiological conditions. To do so, we co-expressed rat TRPV1 (rTRPV1) with the bright probe construct (and its substrate luxCDE) in HEK293 cells and stimulated the cells using capsaicin [55]. rTRPV1 has a large single-channel conductance [56]. Therefore, capsaicin-induced activation of rTRPV1 causes depolarization due to a large inward cation current [56]. Bioluminescent imaging performed under these conditions also exhibited a signal increase when subjected to the chemical stimulus (*See section 3 and* Figure S7 *in the SI*)

### 4.7 In vivo Activity of AMBER

We created *C. elegans* transgenic lines to test AMBER function *in vivo*. We chose *C. elegans* for these analyses due to the optically transparent properties of the tissues, and the slow kinetics of the muscle and neuronal voltage signals (about a few seconds). We subcloned the AMBER (and its substrate) cDNAs in *C. elegans* vectors (*See Section 1 in the SI*) and expressed them in pharyngeal muscles and mechanosensory touch neurons under the control of the *myosin-2 heavy chain* (*myo-2*) promoter [57] and the *β-tubulin* (*mec-7*) promoter [58,59], respectively. We recorded bioluminescent signals using a custom-built imaging set up that allowed tracking the positions of the animal simultaneously (*See Sections 6 and 7 the SI* and Figure S8).

First, we wanted to confirm that AMBER is capable of reporting physiological voltage signals. We expressed AMBER in the pharyngeal muscles of *C. elegans* and recorded (*See Section 8 in the SI*) the voltage activity of the corpus, isthmus and terminal bulb (TB) during natural feeding. We tracked movement of the TB muscles during feeding by simultaneously recording the bright field images (See Figures 5a-b and Supplementary Movie SM1). *C. elegan*s uses pharyngeal pumping to concentrate its food, and starvation drives neuromuscular signals [60] causing rhythmic pumping (in the corpus, isthmus and TB), peristaltic movements (in the isthmus) and grinding actions (in the TB). A few earlier studies reported voltage activity of the serotonin-induced pharyngeal pumping using fluorescent GECIs [61,62] and GEVIs [63]. Here, we report the voltage activity of the three components during natural feeding on bacteria.

**Figure 5:**
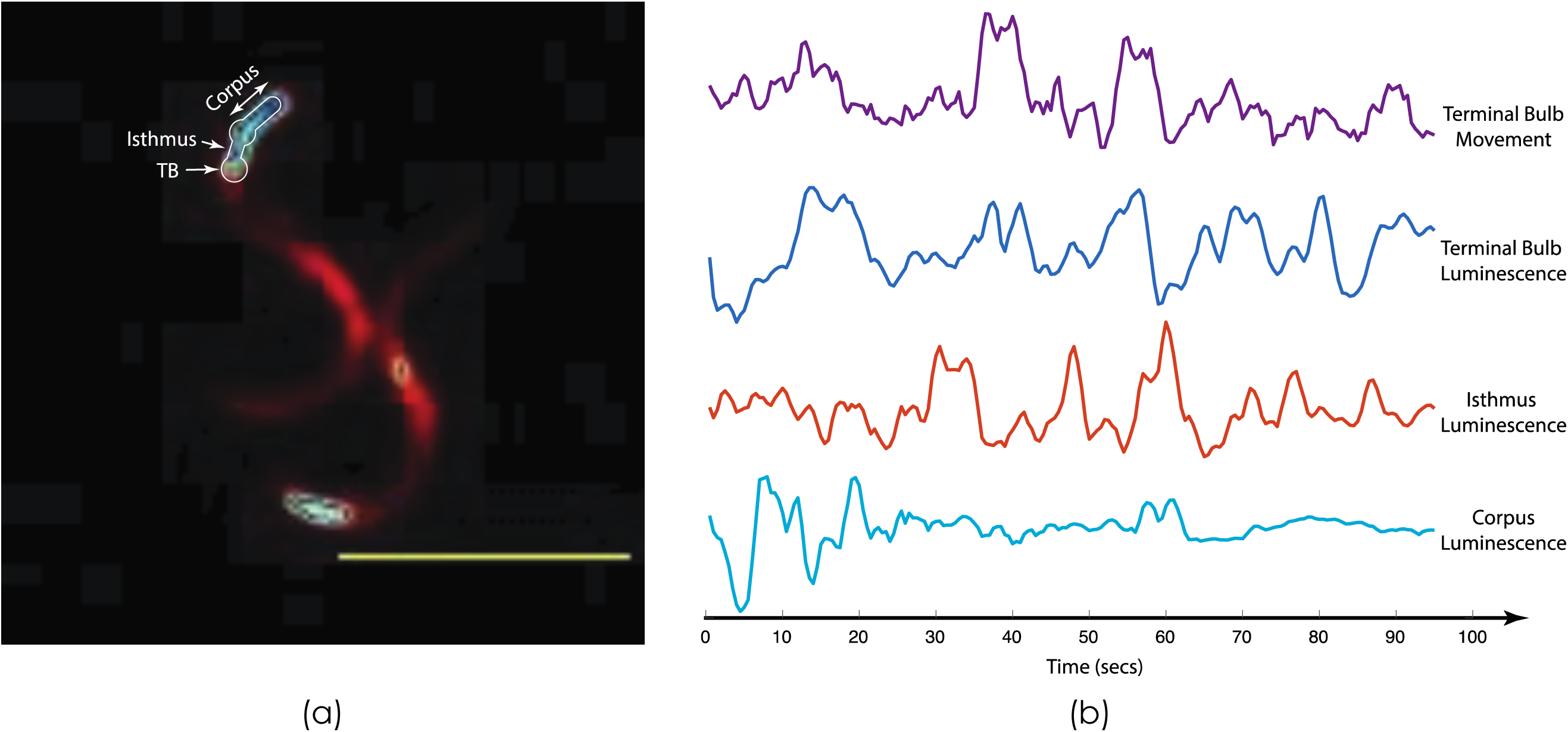
(a) In vivo voltage imaging of adult C.elegans pharyngeal muscle activity during natural feeding. (b) Live tracking of the Terminal Bulb (TB) movement during voltage imaging of TB, Isthmus and Corpus using AMBER. The voltage trace of TB follows closely the shape of TB movement. In contrast, the activity of Isthmus shows a time lead with respect to the TB activity. The Corpus trace is not well resolved due to the inability to map its position precisely. The scale bar corresponds to 500*μ*m.

We found the filtered trace of the voltage activity recorded using AMBER was a direct reflection of the muscle movement. AMBER detects the key features of the muscle kinetics (contracted and relaxed states) as observed previously in serotonin-induced pumping events (between 0.5−1 Hz) using a fluorescent GECI, DRIP [62]. However, unlike the slow DRIP kinetics, AMBER faithfully reproduced the sharp jumps of the muscle motion, as indicated by the voltage transients (See the TB traces corresponding to the voltage activity and muscle movement in Figure 5b and Figure S9). We also observed variability in the voltage traces between the animals reflecting the diversity of their feeding behavior [64,65]. AMBER enables recording voltage activity at multiple time scales (seconds to tens of minutes), which can provide new insights into the animal behavioral outputs. These observations demonstrate the ability of AMBER to sense physiological voltage *in vivo*.

Next, we expressed both the probe and the substrate (*See Sections 1, 6 and 9 in the SI*) in mechanosensory touch neurons (ALMR, ALML, AVM, PLMR, PLML, PVM) and the anterior nerve ring (See Figure S10 for *mec-7* promoter driven eGFP expression). Signals from the touch neurons did not show any activity when the worm was moving unilaterally in the forward direction but showed bursts of activity while making spatially localized movements and frequent reversals (see supplementary movies SM2, SM3 and Figure 6a) or during collisions (Figure 6b). The mechanosensory touch neurons responded to differential force, but not to constant force [66,67]. We observed activities in the touch neurons of animals during collisions, which may be due to transient differential forces caused by momentum transfer. The underlying mechanics in other cases (frequent reversals and localized movements) could not be teased out, although it is known that such motions on the agar bed help the animals to detect bacteria on the plate [68]. A huge advantage of AMBER is the ability to simultaneously record the activity of a neural circuit from multiple worms moving in different directions within the field of view, which is not possible using fluorescence (Figure 6c and Supplementary Movie SM4).

**Figure 6:**
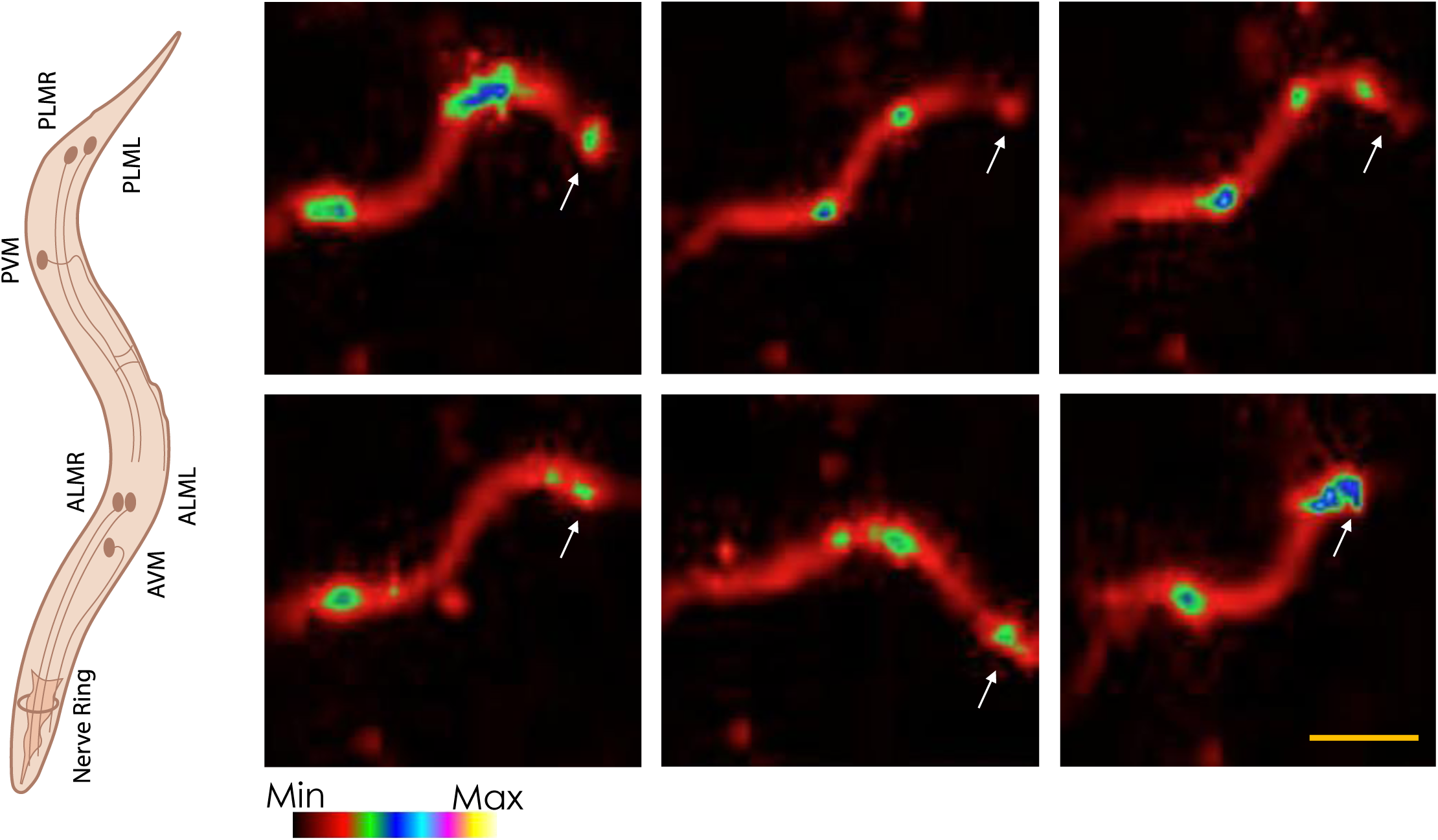
(a) Snapshots of the touch neurons activities in a single worm obtained using the bright probe. The worm takes different shapes as shown while performing frequent reversal movements causing graded activity in the mechanosensory touch neurons (PLM, PVM, ALM, AVM and the anterior nerve ring neurons). Arrow heads indicate the anterior of the worm. The scale bar corresponds to 500*μ*m.

**Figure 6:**
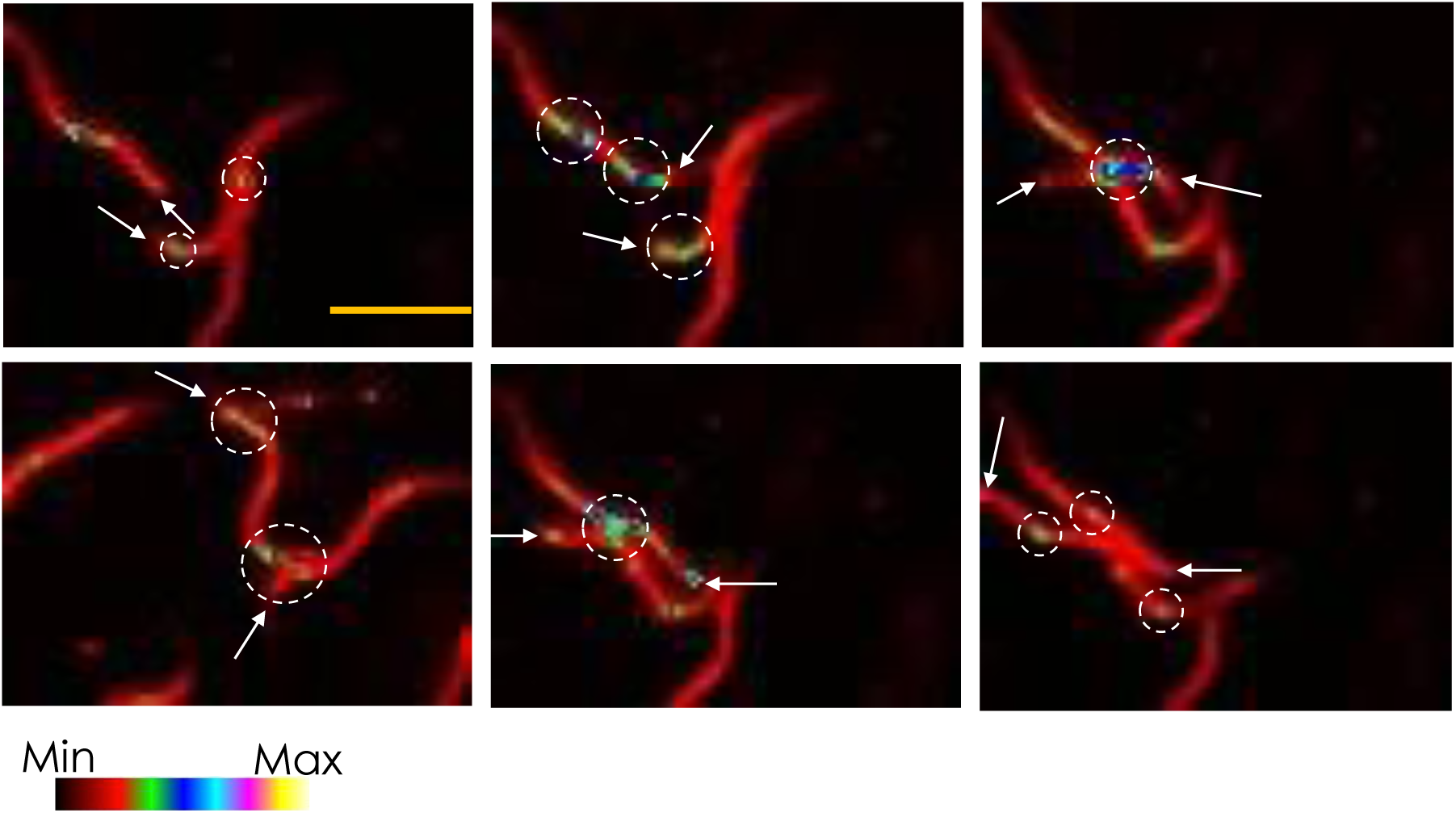
(b) Snapshots of activities of the touch neurons in a population of worms during collision. Touch neurons respond to differential forces exerted on the colliding worms due to momentum transfer. Arrow heads indicate the anterior of the worms. The scale bar corresponds to 500*μ*m.

**Figure 6c:**
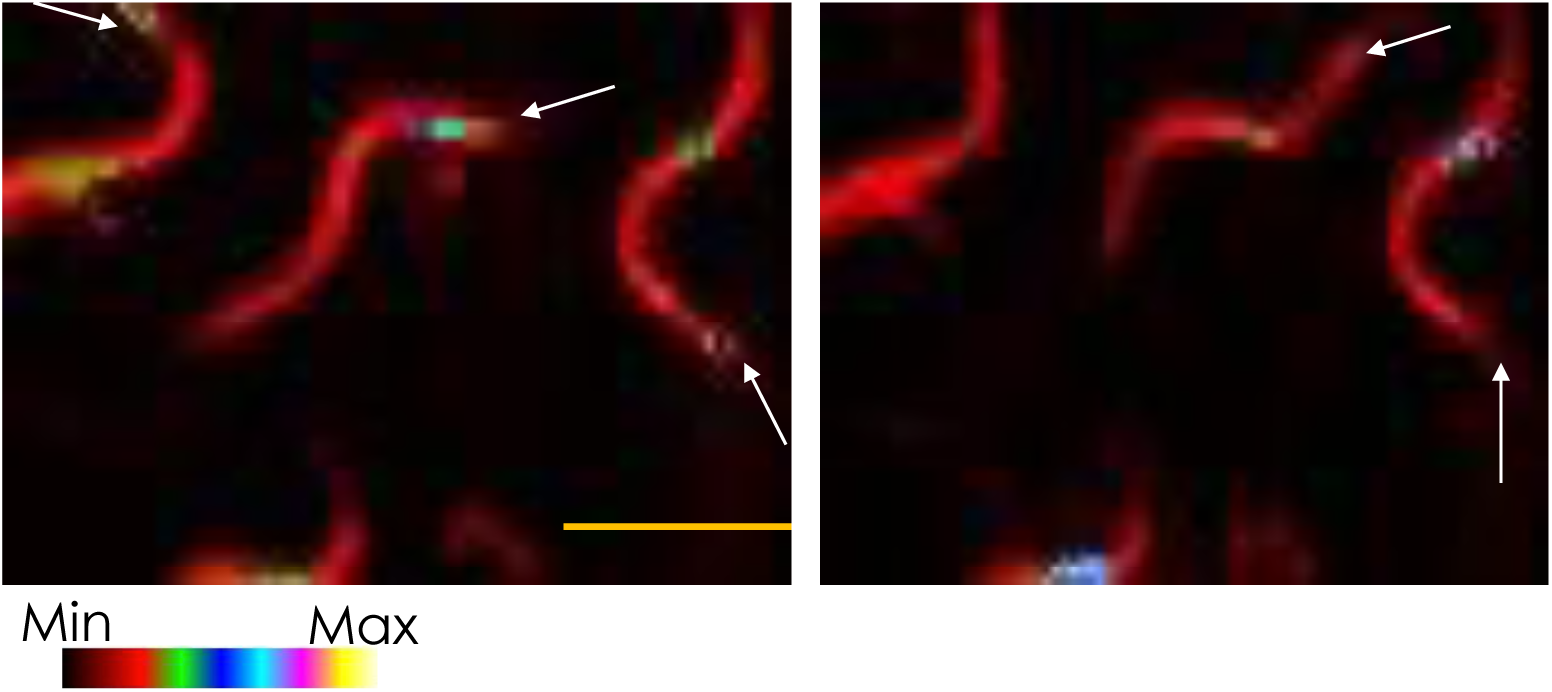
Snapshots of varying levels of activity in touch neurons from a population of worms recorded in a single field of view. Arrow heads indicate the anterior of the worms. The scale bar corresponds to 500*μ*m.

## 5. Discussion

We report the creation of a first-generation voltage-sensitive bioluminescent probe (AMBER), which uses a *Ciona* VSD, a synthetic enhanced bacterial luciferase, eluxAB and a β barrel protein. The system is autonomous and enables expression of both the probe and substrate genetically, circumventing the major drawback of other bioluminescent systems. We engineered the placement of various domains to achieve a high SNR and efficient modulation of the enzymatic activity (See Figures S11-S13). Of particular importance, using *C. elegans*, we demonstrate the effectiveness of AMBER in monitoring voltage changes simultaneously in many free moving animals exhibiting normal behavior.

We performed several experiments to understand the mechanism through which AMBER switches its enzymatic activity reversibly with membrane potential. Direct modulation of fluorescence is known to occur in GEVIs where the fluorescent protein is fused directly to the C-terminus of the Ci-VSD. We ruled out direct bioluminescence modulation by designing a construct with eluxAB fused to the C-terminus of VSD. Though the construct showed some weak signals it did not show a large change in the light emission as observed in the FRP-VSD-eluxAB-YPet. Dimerization of the acceptor fluorescent protein can cause a voltage-dependent FRET in fluorescent GEVIs [69]. We ruled out this mechanism using a dark mutant of YPet that showed voltage-dependent bioluminescent signals but no acceptor resonance energy transfer. Additional evidence comes from observing the activity of FRP-VSD-YPet-eluxAB, which exhibits poor voltage modulation. Given this observation, one possibility is that the position of YPet might have prevented dimerization and therefore exhibits no modulation; implying dimerization is the key to the modulation. However, if the undimerized state is the bright or ‘on’ state, this particular construct should constantly be emitting a large bioluminescent signal, which is not observed in experiments (See Figure 2a).

The presence of the YPet β barrel could structurally modulate the coupling between eluxA and eluxB domains. To rule this out we used the construct that has both the YPet and eluxAB positions switched (FRP-VSD-YPet-eluxAB). As mentioned previously, this construct was the poorest performing among the constructs we tested. If the movement of the VSD S4 transmembrane domain had coupled eluxA and eluxB non-optimally mediated by the YPet, then this construct should show an increase in the effect because YPet is closer to VSD than eluxAB. However, this was not observed.

The availability of FMNH_2_ is necessary for light production, and is therefore a rate limiting substrate. FRP, the enzyme that catalyze the conversion of FMN to FMNH_2_ is directly coupled to the N-terminus of VSD. It is possible that the rate of FMNH_2_ production is voltage-dependent, thereby modulating the luminescence. However, we excluded this mechanism as well by creating constructs with and without FRP, and measuring the luminescence output. Although the brightest construct was the one with FRP on the N-terminus (with eluxAB-YPet in the C-termini), when compared to their counterparts (with YPet-eluxAB in the C-termini), the constructs without FRP showed higher modulation thereby eliminating this proposal.

Lastly, the movement of the VSD could directly modulate the accessibility of the substrate to eluxAB by modifying the substrate-binding pocket. This possibility is excluded by the results from the construct with only eluxAB coupled to VSD. We observed almost no switching as evidenced by the SNR plot in Figure 2b.

We suggest that one plausible mechanism for the voltage dependent activity could be the structural arrangement of FRP-VSD-eluxAB-YPet indirectly resulting in the modulation of the substrate accessibility. We speculate that the presence of the β barrel structure of YPet prevents substrate access to eluxAB before depolarization and adopts an open confirmation when the voltage changes.

Currently, our experimental setup does not allow quantifying the voltage-dependent bioluminescent kinetics. However, there are some general trends that we observed. In most experiments, the integration time was 1 sec, and we have been able to observe *in vivo* signals with a 500 msec integration time in cases where we obtained high expression levels of the probe proteins. This can be seen from the experiments performed on *C. elegans*. High expression in the pharyngeal muscles allowed us to detect voltage changes in 500 msec, thereby enabling to track the muscle movement with voltage signals (Figure 5a-b).

Our creation of AMBER proteins show for the first time the ability to modulate enzymatic activity of eluxAB by varying the membrane potential to realize a voltage-gated luciferase. We propose that voltage imaging using bioluminescence as described here will have a broad applicability.

## 6. Conclusions

We engineered an autonomous voltage-sensitive bioluminescent reporter based on the bacterial bioluminescent system. The ability to genetically encode the luciferase and the luciferin overcomes many challenges inherent with firefly and the deep-sea shrimp bioluminescent systems. Moreover, we optimized the performance of the probe to emit bright signal after depolarization using molecular engineering approaches. AMBER exhibits a voltage-dependent enzymatic switching not observed in other similar systems (e.g. LOTUS-V). We successfully expressed and demonstrated AMBER function both *in vitro* (HEK293 cells) and *in vivo* (C. *elegans*). We reported the voltage activities of the mechanosensory neural circuit and the pharyngeal muscles from multiple animals using AMBER. We believe this will greatly enhance the ability to track the behavior of multiple animals, while simultaneously monitoring the activities of neurons.

## 7. Acknowledgements

This work was partially funded by the Otis Williams Fellowship (PS), the NSF Neuronex (Award #1934288 LT), the National Eye Institute (EY008117 and EY010852; CM), the National Institute of Health (1R01HD082347 and 1R01HD081266; JR), the American Heart Foundation (19POST34430179; DT) and the Institute of Collaborative Biotechnologies at the UCSB. The authors are grateful to Sumita Pennathur, UCSB for generously lending the EMCCD camera and Spencer Smith, UCSB for helping build the patch clamp set up. The authors acknowledge the use of the Biological Nanostructures Laboratory within the California NanoSystems Institute, supported by the University of California, Santa Barbara and the University of California, Office of the President. The authors also thank Dan Close and Gary Sayler at 490 Biotech for providing us the lux plasmid. We acknowledge insightful discussions with Ute Hochgeschwender (Central Michigan University) and Nathan Shaner (UC San Diego).

## Notes

### Competing Interest Statement

The authors have declared no competing interest.

### Summary of Updates

Fig.2C was plotted incorrectly in the earlier version. A new amended plot is included in the revised version. We also included the MATLAB code used for processing the in vivo data in the SI appendix file.

## References

1. Mehta, S. & Zhang, J. Reporting from the Field: Genetically encoded fluorescent reporters uncover signaling dynamics in living biological systems. Annu Rev Biochem. 80, 375–401 (2011)

2. Lin, M. Z., & Schnitzer, M. J. Genetically encoded indicators of neuronal activity. Nat. Neurosci. 19, 1142–1153 (2016)

3. Xu, Y., Zou, P. & Cohen, A. E. Voltage imaging with genetically encoded indicators. Curr. Opin. Chem. Biol. 39, 1–10 (2017).

4. Qian, Y. et al. A genetically encoded near-infrared fluorescent calcium indicator. Nat. Methods, 16, 171–174 (2019).

5. Tian, L., Andrew Hires, S., & Looger, L. L. Imaging neuronal activity with genetically encoded calcium indicators. Cold Spring Harb protoc. doi:10.1101/pdb.top069609 (2012)

6. Bischof, H. et al., Novel genetically encoded fluorescent probes enable real-time detection of potassium in vitro and in vivo. Nat. Comm. 8, 1422 (2017).

7. Shen, Y. et al. Genetically encoded fluorescent indicators for imaging intracellular potassium ion concentration. Commun. Biol. 2, 18 (2019).

8. Liang, R., Broussard, G. J., & Tian, L. Imaging chemical neurotransmission with genetically encoded fluorescent sensors. ACS Chem. Neurosci. 6, 84–93 (2015).

9. Okumoto, S. et al. Detection of glutamate release from neurons by genetically encoded surface-displayed FRET nanosensors. Proc. Natl. Acad. Sci. USA. 102, 8740–8745 (2005)

10. Marvin, J. S. et al. An optimized fluorescent probe for visualizing glutamate neurotransmission. Nat. Methods. 10, 162–170 (2013)

11. Yang, H. H. & St-Pierre, F. Genetically encoded voltage indicators: Opportunities and Challenges. J. Neurosci. 36, 9977–9989 (2016).

12. Abdelfattah, A. S., et al. A bright and fast red fluorescent protein voltage indicators that reports neuronal activity in organotypic brain slices. J. Neurosci. 36, 2458–2472 (2016).

13. Kannan, M. et al. Fast in vivo voltage imaging using a red fluorescent indicator. Nat. Methods. 15, 1108–1116 (2016)

14. Yang, W. & Yuste, R. In vivo imaging of neural activity, Nat. Methods. 14, 349–359 (2017).

15. Stirman, J. N. et al. Wide field-of-view, multi-region two-photon imaging of neuronal activity in the mammalian brain. Nat. Biotechnol. 34, 857–862 (2016)

16. Fosque, B. F. et al. Labeling of active neural circuits in vivo with designed calcium integrators. Science. 347, 755–760 (2015).

17. Zhao, Y. et al. An expanded palette of genetically encoded Ca^2+^ indicators. Science. 333, 1888–1891 (2011).

18. Hang, H. H., St-Pierre, F., Sun, X., Ding, X., Lin, M. Z. & Clandinin, T. R. Subcellular imaging of voltage and calcium signals reveals neural processing in vivo. Cell. 166, 245–257 (2016)

19. Wei, Z., Lin, B., Chen, T., Daie, K., Svoboda, K. & Druckmann, S. A comparison of neuronal population dynamics measured with calcium and electrophysiology. bioRxiv 840686; doi: https://doi.org/10.1101/840686

20. Burgoyne, R. D. Neuronal Calcium sensor proteins: Generating diversity in neuronal Ca^2+^ signaling. Nat. Neurosci. 8, 182–193 (2007).

21. Grienberger, C. & Konnerth, A. Imaging calcium in neurons. Neuron. 73, 862–885 (2012)

22. Knöpfel, T. Genetically encoded optical indicators for the analysis of neuronal circuits. Nat. Rev Neurosci. 13, 687–700 (2012).

23. Antic, S. D., Empson, R. M. & Knöpfel, T. Voltage imaging to understand connections and functions of neuronal circuits. J. Neurophysiol. 116, 135–152 (2016).

24. Piatkevich, K. D. et al. Population imaging of neural activity in awake behaving mice. Nature. 574, 413–417 (2019).

25. Abdelfattah, A. S. et al., Bright and photostable chemigenetic indicators for extended in vivo voltage imaging. Science. 365, 699–704 (2019).

26. Rose, A. Vision: Human and Electronic (Springer, 2013).

27. Bando, Y., Sakamoto, M., Kim, S., Ayzenshtat, I. & Yuste, R. Comparative evaluation of genetically encoded voltage indicators. Cell Reports. 26, 802–813 (2019)

28. Lorenz, W. W., McCann, R. O., Longiaru, M. & Cormier, M. J. Isolation and expression of a cDNA encoding renilla reniformis luciferase. Proc. Natl Acad. Sci. USA 88, 4438–4442 (1991).

29. Markova, S. V., Golz, S., Frank, A. A., Kalthof, B. & Vysotski, E. S. Cloning and expression of cDNA for a luciferase from the marine copepod Metridia *longa*, a novel secreted bioluminescent reporter enzyme. J. Biol. Chem. 279, 3212–3217 (2004).

30. Thompson, E. M., Nagata, S. & Tsuji, F. I. Cloning and expression of cDNA for the luciferase from the marine ostracod Vargula *hilgendorfii*. Proc. Natl Acad. Sci. USA 86, 6567–6571 (1989).

31. Verhaegen, M. & Christopoulos, T. K. Recombinant Gaussia luciferase. Overexpression, purification, and analytical application of a bioluminescent reporter for DNA hybridization. Anal. Chem. 74, 4378–4385 (2002).

32. Wood, K. V., Lam, Y. A. & McElroy, W. D. Introduction to beetle luciferases and their applications. J. Biolumin. Chemilumin. 4, 289–301 (1989).

33. Kaskova, Z. M. et al. Mechanism and color modulation of fungal bioluminescence. Sci. Adv. 3, e1602847 (2017)

34. Brodl, E., Winkler, A. & Macheroux, P. Molecular mechanisms of bacterial bioluminescence. Comput. Struct. Biotechnol. J. 16, 551–564 (2018).

35. Kotlobay, A. A. et al. Genetically encodable bioluminescent system from fungi. Proc. Natl Acad. Sci. USA 115, 12728–12732 (2018).

36. Shieh, Y. W. et al. Operon structure and cotranslational subunit association direct protein assembly in bacteria, Science, 350, 678–80 (2015)

37. Gregor, C. et al. Strongly enhanced bacterial bioluminescence with the ilux operon for single-cell imaging. Proc. Natl Acad. Sci. USA. 115, 962–967 (2018)

38. Close, D. M. et al. Autonomous bioluminescent expression of the bacterial luciferase gene cassette (*lux*) in a mammalian cell line. PLoS ONE, 5, e12441 (2010).

39. Hollis, R. P. et al. Toxicity of the bacterial luciferase substrate,n-decyl aldehyde, to *Saccharomyces cerevisiae* and *Caenorhabditis elegans*. FEBS lett. 506, 140–142 (2001).

40. Gregor, C. et al. Autonomous bioluminescence imaging of single mammalian cells with the bacterial bioluminescence system, Proc. Natl Acad. Sci. USA. 116, 26491–26496 (2019)

41. Li, Q. et al. Structural mechanism of voltage-dependent gating in an isolated voltage-sensing domain. Nat. Struct. Mol. Biol. 21, 244–252 (2014)

42. Nguyen A. W. & Daugherty, P. S. Evolutionary optimization of fluorescent proteins for intracellular FRET. Nat. biotech. 23, 355–360 (2005).

43. Dimitrov, D. et al. Engineering and characterization of an enhanced fluorescent protein voltage sensor, PLoS ONE. 2, e440 (2007).

44. Inagaki, S. et al. Genetically encode bioluminescent voltage indicator, Sci. Rep. 7, 42398 (2017).

45. Hall, M. P., et al. Engineered luciferase reporter from a deep sea shrimp utilizing a novel imidazopyrazinone substrate. ACS Chem. Biol. 7, 1848–1857 (2012).

46. Cui, B. et al. Engineering an enhanced, thermostable, monomeric bacterial luciferase gene as a reporter in plant protoplasts. PLoS ONE. 9, e107885 (2014).

47. Lei, B. & Tu, S. Mechanism of reduced flavin transfer from Vibrio *harveyi* NAD(P)H-FMN oxidoreductase to luciferase. Biochemistry. 37, 14623--14629 (1998).

48. Tu, S. Activity coupling and complex formation between bacterial luciferase and flavin reductases. Photochem. Photobiol. Sci. 7, 183–188 (2008).

49. Jawanda, N., Ahmed, K. & Tu, S. Vibrio *harveyi* Flavin reductase-luciferase fusion protein mimics a single-component bifunctional monooxygenase. Biochemistry. 47, 368–377 (2008).

50. Kutrowska, B. W. et al. Folding and unfolding of a non-fluorescent mutant of green fluorescent protein. J. Phys Condens Matter. 18, 285223 (2007)

51. Schaefer, P. M., Kalinina, S., Rueck, A., von Arnim, C. A. F. & von Einem, B. NADH Autofluorescence – A marker on its way to boost bioenergetic research. Cytometry A. 95A, 34–46 (2019).

52. Sung, U. et al. Developing fast fluorescent protein voltage sensors by optimizing FRET interactions. PLoS ONE. 10, e0141585 (2015)

53. Akemann, W. et al. Effect of voltage sensitive fluorescent proteins on neuronal excitability. Biophys. J. 96, 3959–3976 (2009)

54. Jin, L., Han, Z., Platisa, J., Wooltorton, J. R., Cohen, L. B. & Pieribone, V. A. Single action potentials and subthreshold electrical events imaged in neurons with a novel fluorescent protein voltage probe. Neuron. 75, 779–785 (2012)

55. Caterina, M. J., Schumacher, M. A., Tominaga, M., Rosen, T. A., Levine, J. D. & Julius, D. The capsaicin receptor: a heat-activated ion channel in the pain pathway. Nature, 389, 816–824 (1997)

56. Cao, E., Cordero-Morales, J. F., Liu, B., Qin, F. & Julius, D. TRPV1 channels are intrinsically heat sensitive and negatively regulated by phosphoinositide lipids, Neuron. 77, 667–679 (2013).

57. Okkema, P. G., Harrison, W. W., Plunger, V., Aryana, A. & Fire, A. Sequence requirements for myosin gene expression and regulation in *Caenorhabditis elegans*. Genetics. 135, 385–404 (1993)

58. Hamelin, M., Scott, I. M., Way, J. C. & Culotti J. G. The mec-7 beta-tubulin gene of *Caenorhabditis elegans* is expressed primarily in the touch receptor neurons. EMBO J. 11, 2885–93 (1992)

59. Zhang, Y. et al. Identification of genes expressed in C. elegans touch receptor neurons. Nature. 418, 331–335 (2002)

60. Avery, L. and You, Y.J. *C. elegans* feeding (May 21, 2012), WormBook, ed. The C. elegans Research Community, WormBook, doi/10.1895/wormbook.1.150.1, http://www.wormbook.org.

61. Kerr, R., Lev-Ram, Varda., Baird, G., Vincent, P., Tsien, R. Y. & Schafer, W. R. Optical imaging of calcium transients in neurons and pharyngeal muscle of C.elegans. Neuron. 26, 583–594 (2000)

62. Shimozono, S., Fukano, T., Kimura, K. D., Mori, I., Kirino, Y. & Miyawaki, A. Slow Ca^2+^ dynamics in pharyngeal muscles in Caenorhabditis elegans during fast pumping. EMBO Rep. 5, 521–526 (2004)

63. Hashemi, N. A. et al. Rhodopsin-based voltage imaging tools for use in muscles and neurons of *Caenorhabditis elegans*. Proc. Natl Acad. Sci. USA 116, 17051–60 (2019)

64. Scholz, M., Lynch, D. J., Lee, K. S., Levine, E. & Biron, D. A scalable method for automatically measuring pharyngeal pumping in C. elegans. J. Neurosci. Methods. 274, 172–178 (2016)

65. Rodriguez-Palero, M. J., et al. An automated method for the analysis of food intake behavior in Caenorhabditis elegans. Sci. Rep. 8, 3633 (2018)

66. Goodman, M. B. Deconstructing C.elegans sensory mechanotransduction. Science. 306, 427–428 (2004).

67. Krieg, M., Dunn, A. & Goodman, M. B. Mechanical systems biology of C.elegans touch sensation. Bioessays. 37, 335–344 (2015)

68. Goodman, M.B. Mechanosensation (January 06, 2006), WormBook, ed. The C. elegans Research Community, WormBook, doi/10.1895/wormbook.1.62.1, http://www.wormbook.org.

69. Yi, B., Kang, B. E., Lee, S., Braubach, S. & Baker, B. J. A dimeric fluorescent protein yields a bright, red-shifted GEVI capable of population signals in brain slice. Sci. Rep., 8, 15199 (2018)

