## Supplementary material for "An Autonomous Molecular Bioluminescent Reporter (AMBER) for voltage imaging in freely moving animals": SI appendix

<u>Supplementary Information</u>

This document includes supplementary **Sections 1-13**, supplementary **Tables S1-S3**, supplementary **Figures S1-S14**, and legends for supplementary movies **SM1-SM4**.

### 1. Molecular cloning of Engineered AMBER constructs

We used two plasmids – $pCMV_{lux}$ and pcDNA 3.1-VSFP2.1 to construct a mammalian plasmid vector that expresses engineered protein constructs of first-generation AMBER. $pCMV_{lux}$ that codes for mammalian codon-optimized lux operon was purchased from 490 Biotech. pcDNA 3.1-VSFP2.1 was a gift from Thomas Knopfel laboratory (Addgene plasmid # 1255; http://n2t.net/addgene:16255; RRID: Addgene_16255).

We applied a structure-function relationship to assess the functional equivalence of luxAB-YPeT BRET pair as a replacement for Cerulean/Citrine FRET pair in VSFP2.1. Both YPet and Citrine were derived from YFP (a GFP variant) [1,2] and have similar beta scaffold structure. However, *P. luminescens* luxAB (pluxAB) structure is not yet solved although it shares a comparable amino acid sequence identity (89% *luxA* and 54% *luxB*) with *V. harveyi* luxAB (vluxAB) whose structure is known [3,4]. vluxAB halo enzyme has a characteristic TIM barrel structure [3] showing discrete vluxA and vluxB domains. A fusion of vluxA and vluxB domains with a polypeptide linker catalyzes light reaction albeit with varied activity depending on linker length [5]. We, therefore, hypothesized that replacing FRET pair of VSFP2.1 with the luxAB/YPet BRET pair is a good starting point for subsequent structural optimizations.

We used the structural framework of VSFP2.1 [6] for designing the first generation AMBER. Our cloning strategy involved three principal steps; (i) mutational changes of the base pairs for creating compatible restriction sites, (ii) Synthesis of double-stranded insert fragments either by restriction endonuclease or Polymerase Chain Reaction (PCR using NEB Phusion-HF DNA polymerase; Catalog# M0530L) amplification of template cDNA with suitable primers and (iii) Fusion of vector backbone and the insert by a ligation reaction. We described the molecular biology approaches used to sub-clone the cDNAs of various constructs in the mammalian and *C.elegans* expression vectors. All expression vectors are shuttle vectors that allow genetic manipulation in bacteria and protein expression in the host cells. Restriction digestions were done mostly with high fidelity restriction enzymes (HF versions, New England Biolabs) overnight to ensure complete digestion of the substrates at the specific sites. Ligations were performed either with T4 DNA ligase (Catalog # M0202S; New England Biolabs inc.) or in-fusion enzyme mix (Catalog # 639649; TaKaRa Bio Inc.) following manufacturer's protocol. We used Stellar competent cells (Catalog # 636766; TaKaRa Bio Inc.) for transformation reactions of ligated products and all transformed products were plated on LB Agar plates with suitable antibiotic selection.

We posited that replacing Cerulean of VSFP2.1 [6] with the mammalian codon-optimized synthetic luciferase, enhanced luxAB [7] (henceforth called as 'eluxAB') would provide the necessary structural framework for the optimization of engineered protein constructs. The Citrine domain of VSFP2.1 was modified to YPet, the brightest variant of YFP whose absorption spectrum substantially overlap with a broad luxAB emission spectrum. Fused luxAB retains its enzymatic activity [8]; therefore, we created a fusion linker between eluxA and eluxB in all the engineered AMBER constructs. Implementation of these modifications to the VSFP2.1 plasmid results in a plasmid coding for VSD-eluxAB-YPet protein. The remaining components of lux operon, *luxCDE-FRP* was kept intact in the $pCMV_{lux}$ backbone allowing polycistronic co-expression of luxCDE-FRP with the membrane-targeted VSD-eluxAB-YPet. Table S1 lists all the relevant biochemical reactions performed to construct the plasmids coding for all the engineered AMBER protein constructs.

#### ***VSD-eluxAB-YPet***

We began with replacing the *Cerulean* genetic sequence in pcDNA3.1-VSFP2.1 with the *luxAB* domain of $pCMV_{lux}$. This led to engineering plasmids that code several other AMBER constructs discussed in this work. The *Cerulean* sequence was flanked with NotI (1671) and BamHI (2374) sites. While the NotI site was unique, there were two BamHI sites (2374 and

3120). To create a unique restriction site flanking the *Cerulean* sequence, the BamHI site at 3120 was removed by modifying the bases so that the open reading frame codes for the same amino acid using an alternative codon (a3122c). Unlike the *Cerulean* sequence in pcDNA3.1-VSFP2.1, there were no flanking restriction sites across the *luxAB* domain in $pCMV_{lux}$. These sites were therefore created using site-directed mutagenesis substitution reaction. The requirement for minimal base pair changes (up to 4) allowed a NotI site to be placed 270bp upfront of *luxA* domain thereby including a part of *luxD* domain in the chosen insert fragment (2300 bp). The *Cerulean* sequence was replaced with the insert fragment synthesised from the $pCMV_{lux}$ using restriction endonuclease (BamHI-HF and NotI-HF enzymes) and ligation (T4 DNA ligase) reactions. The additional 270bp representing a part of the *luxD* domain at the 5' end of the insert was later removed by a deletion mutagenesis reaction to obtain *vsd-luxA-luxB-YFP*. Functional mutations within the *luxA, luxB* and *Citrine* sequences were introduced to create enhanced luciferase domains, *eluxA* and *eluxB* [7] and *YPet* [2] respectively. Furthermore, $pCMV_{lux}$ contains self-cleaving viral 2A genetic sequence between various components of *lux* operon in the open reading frame to allow polycistronic expression of the respective proteins. We therefore created functional mutants of viral 2A sequences reported previously [9] to abrogate self-cleavage between eluxA and eluxB proteins. The resulting modified cDNA, *VSD-eluxAB-YPet* encodes for a single chimeric protein fusing Ciona voltage sensor, VSD, enhanced luciferase, *eluxAB* and a bright fluorescent reporter, *YPet*.

***VSD-YPet-eluxAB***

An additional BamHI site was created by substituting base pairs of EcorI at the 3' end of *VSD-eluxAB-YPet* sequence. This allowed cleaving the *YPet* domain using restriction digestion to create small (*YPet* gene) and large fragments with flanking BamHI sites. Self-ligation of the large fragment creates a plasmid that encodes *VSD-eluxAB*. After interchanging the locations of BamHI and NotI sites in the *VSD-eluxAB*, the double stranded *YPet* fragment was introduced at the N-terminus of VSD the using restriction endonuclease and ligation reactions. This creates a plasmid that encodes *VSD-YPet-eluxAB*.

***luxCDE-FRP***

We excised out the double stranded DNA fragments that encodes *luxA* and *luxB* using the $pCMV_{lux}$ plasmid by a deletion mutagenesis reaction. Genetic sequence between the first nucleobase of *luxA* to the last nucleobase of *P2A* including the end base pairs was removed without disrupting the reading frame. This reaction results in a plasmid that encodes *luxCDE-FRP.*

***FRP-VSD-eluxAB-YPet***

We used a single BmtI site at the 5' end of *VSD-eluxAB-YPet* to introduce the *FRP* sequence. *FRP* gene with flanking BmtI sites was cloned in the *luxCDE-FRP* by a substitution mutagenesis reaction. This was followed by restriction endonuclease at BmtI sites to obtain a small fragment (*FRP* sequence*)* and a large fragment (*luxCDE* sequence). A single site ligation of the *FRP* insert at the BmtI site of the plasmid encoding *VSD-eluxAB-YPet* results in a new plasmid that encodes for *FRP-TAA-VSD-eluxAB-YPet.* The *TAA* stop codon at the 3' end of *FRP* gene transferred from the $pCMV_{lux}$ was mutated to a *GGA* (Glycine linker) to obtain the bright construct, *FRP-VSD-eluxAB-YPet*. Self-ligation of the large fragment resulted in the plasmid *luxCDE*, which encodes for the necessary substrate generating protein complexes.

***FRP-VSD-YPet-eluxAB***

We used the unique BmtI site at the 5' end of *VSD-YPet-eluxAB* to introduce the *FRP* gene. The *FRP* gene fragment with flanking BmtI sites created previously was introduced at the N-terminus of *VSD-YPet-eluxAB* using restriction endonuclease and ligation reactions. These reactions were followed by the substitution mutation of the *TAA* stop codon to *GGA* (Glycine linker) to create a plasmid that encodes for *FRP-VSD-YPet-eluxAB.* The *luxCDE* plasmid constructed earlier encodes for the necessary substrate-generating protein complexes.

***luxAB***

We fused the *luxA* and *luxB* genes in the $pCMV_{lux}$ by mutating the *T2A* element between them so that self-cleavage between these domains during protein synthesis is abrogated. This results in a plasmid that expresses soluble fusion construct *luxAB*.

***pCDNA3.1-rTRPV1***

The cDNA encoding for the recombinant fusion protein, *MBP-8xHis-rTRPV1* (fusion of maltose binding protein with polyhistidine tag to rat *TRPV1*) cloned in an insect cell expression vector was a gift from David Julius laboratory, University of California, San Francisco. We sub-cloned the *rTRPV1* domain into the mammalian expression vector pcDNA3.1(+) using in-fusion cloning approach. This involves restriction endonuclease of the vector at KpnI and EcoRI sites, PCR amplification of the *rTRPV1* cDNA using primers with 15bp vector overlap sequence at 3' and 5' ends, followed by a ligation reaction using in-fusion enzyme mix (TaKaRa Bio Inc; Catalog # 639649). We performed temperature gradient PCR (primer melting temperature, $T_m$ varying from 66–72$^oC$) using 50ng template in a 20µL reaction volume. The primer sequences were optimally designed using Snapgene software (version 3.2.1) to achieve maximal amplification. The PCR products were treated with DpnI enzyme (NEB, R0176S) to cut the methylated parent template before agarose gel electrophoretic separation using TAE buffer (50x Tris-acetate-EDTA buffer, Catalog # FERB49; Fisher Scientific). The excised stranded DNA fragments from the gel were purified using a gel extraction kit (Catalog # 740609.50; TaKaRa Bio Inc) for the ligation.

**Dark Mutant**

Plasmid coding for the dark mutant of *YPet* (Gly65Thr, Gly67Ala) in *FRP-VSD-eluxAB-YPet* was constructed using commercially synthesised *YPet* sequence with appropriate base pair mutations. We purchased synthetic double stranded DNA fragment (Genscript gene blocks) of mutated *YPet* with 15bp vector overlap sequence at the 3' and 5' ends flanking BamHI and EcoRI sites. The fluorescent *YPet* domain of the *FRP-VSD-eluxAB-YPet* was replaced with the mutated *YPet* fragment using restriction endonuclease of *FRP-VSD-eluxAB-YPet* and in-fusion reactions. The large fragment was obtained by electrophoretic separation of the digested products followed by extraction/purification using the gel extraction kit.

**Cloning *FRP-VSD-eluxAB-YPet and luxCDE* in the C.elegans expression vectors**

We sub-cloned *FRP-VSD-eluxAB-YPet and luxCDE* in *C. elegans* expression vectors targeting mechanosensory touch neurons and pharyngeal muscles. We used vectors L3691 (Addgene plasmid # 1587) and L3790 (Addgene plasmid # 1596) obtained as a gift from Andy Fire laboratory. KpnI and EcoRI restriction sites of the L3691 vector were used to insert the cDNAs of *FRP-VSD-eluxAB-YPet and luxCDE* genes separately using infusion-cloning protocols. We optimized the primer sequences for sufficient PCR amplification of the cDNAs using the Snapgene software. PCR reactions were carried out using Phusion-HF DNA polymerase in a 20µL reaction volume containing 50ng of a template (either *FRP-VSD-eluxAB-YPet* or *luxCDE*). *FRP-VSD-eluxAB-YPet* required 3% DMSO (Dimethyl Sulfoxide) addition while *luxCDE* required an additional 0.5mM of $MgCl_2$. All PCR products were treated with DpnI enzyme overnight before electrophoretic separation in TAE buffer and gel purification. The double stranded vector fragments were separately prepared by restriction endonuclease overnight followed by extraction/purification of the fragment. The vector and the insert were fused using the infusion enzyme mix following manufacturer's protocol. We denote the plasmids that express *FRP-VSD-eluxAB-YPet and luxCDE* in *C.elegans* mechanosensory neurons as mec7-Btp ('Btp' represents Bright probe) and mec7-Bts ('Bts' represents Bright substrate) respectively

Plasmids that encode for the *FRP-VSD-eluxAB-YPet* and *luxCDE* for protein expression in the pharyngeal muscles were constructed by replacing the mec-7 promoter sequence of mec7-Btp and mec7-Bts plasmids with a myo-2 promoter sequence from L3790. We performed restriction digestion of L3790 and mec7-Btp plasmids at SphI and ClaI sites and purified the vector and inserts after electrophoretic separation. We then fused the vector and insert using T4 DNA ligase enzyme following manufacturer's protocol. The resultant plasmid obtained from this reaction is denoted as myo2-Btp. myo2-Bts was obtained by insertion of a synthetic myo-2 promoter gene (Genscript geneblocks) with flanking AscI and BamHI sites at the corresponding sites of mec7-Bts using infusion enzyme mix. The double stranded vector fragment with flanking AscI and BamHI sites was constructed by restriction digestion with AscI-HF (NEB; R0558S) and BamHI-HF enzymes followed by agarose gel electrophoretic separation and purification.

**<u>Molecular biology methods for sub-cloning cDNAs into expression vectors</u>**

We performed all substitution, deletion, and insertion mutational changes using QuickChange Lightning Site-Directed Mutagenesis kit (Agilent, Catalog #210518). For implementing base pair changes at multiple locations simultaneously, we used QuickChange Multi Site-Directed Mutagenesis kit (Agilent, Catalog# 200514). The mutational products were treated with DpnI enzyme to cut the parental template before transformation using XL10-Gold Ultracompetent cells (Agilent, Catalog# 200314). Five randomly picked colonies from an Ampicillin selectable plate (10mL of polymerized LB-Agar containing 2% of LB Broth (Lennox; Sigma-Aldrich; Catalog# L7568-1KG), 1.5% of Bacto-Agar solidifying agent (BD Diagnostics, Supplier# 214010) and 1mg of Ampicillin sodium salt (Fisher Scientific, Catalog# BP1760-25)) were screened for every construct. The colonies were inoculated into 3mL LB media containing 1µL/mL of 100mg/mL Ampicllin stock and incubated for 12hrs inside a bacterial incubator (10L Benchmrk Incu-shaker$^{TM}$; Benchmark Scientific, Item# H1010*) at 37$^{o}$C shaking at 225 rpm. Plasmid DNAs purified from single colonies using a DNA miniprep kit (GenCatch Plus Plasmid; Epoch Life Science; Catalog# 21-60250) were sequenced at either Genewiz or Berkeley Sequencing facilities to identify and confirm the positive clones. We used either universal primers or custom-made oligonucleotides to fully sequence the required regions of a plasmid. Plasmid DNA samples purified from the positive clones that yielded high-quality long reads (Q>45) were further sequenced to ensure a 100% match for the nucleobases encoding cDNAs.

### 2. In vitro expression of AMBER proteins in HEK293 cells

HEK293 cells (ATCC certified) were chemically transfected with plasmid DNAs for live bioluminescent imaging. The cell culture protocol involves expanding a frozen stock of cells (low passage number, < 6) in a freshly prepared growth media containing 1X Dulbecco's Modified Eagle's Medium (high glucose DMEM containing 4.5g/L D-Glucose and L-Glutamine; ThermoFisher Scientific; Catalog #11965-092), and 10% heat-inactivated fetal bovine serum (Catalog #10438026, ThermoFisher Scientific). Cells were grown inside a tissue culture incubator (Forma Steri-Cult $CO_2$ Incubator; ThermoFisher Scientific; Catalog# 3307TS) containing 10%$CO_2$ maintained at 37°C and 87%RH. The cells were allowed to recover their stable growth at least over two passages by splitting at 1:10 ratio each time after they reach 80% confluent on a 10 cm tissue culture dish (Catalog #25-202; Genesee Scientific). Each split involves dislodging the attached cells by Trypsin treatment (1mL of Trypsin-EDTA; Life Technologies; Catalog # 25200-056) for 1 min at 37°C after a gentle wash using 1X PBS (ThermoFisher Scientific; Catalog# 10010023). The Trypsin activity was stopped by adding 9mL of the growth media into the plate thereby causing the cells to suspend in solution. The cellular suspension was subjected to differential centrifugation (1000g for 5 min using HERMLE Z300) to retrieve as a pellet at the bottom of a 15mL tube (Catalog #28-103; Genesee Scientific). This step was followed by another round of wash using the growth media to reduce the residual Trypsin if any in the collected cells.

We used $1x10^6$ cells for every batch of transfection experiments. The cells were transfected with a total of 1$\mu$g of DNA (500ng of probe and substrate plasmids for co-transfection) using the commercial Lipofectamine 2000 transfection reagent (ThermoFisher Scientific; Catalog # 11668027). The protocol for preparing the transfection mix involves the preparation of two tubes; one with 1µg DNA solubilized in the Opti-MEM to 50µL and the other with 5µL Lipofectamine reagent mixed with 45mL of Opti-MEM. The contents of the tubes were pre-incubated at the room temperature for 2 min separately before mixing them and incubating for 30 minutes. The transfection mix was then mixed with $1x10^6$ cells (quantified using a hemacytometer, iN CYTO, Catalog # DHC-N01) suspended in 2mL of media and plated directly onto a 60mm tissue culture dish (Corning Inc, Catalog # 430166) placed inside the incubator for 7hrs. Meanwhile, a 12-well plate containing 1% of poly-L-Lysine (Sigma Aldrich, Catalog # P8920-100ML) in PBS was incubated at room temperature inside a UV-sterilised hood. After 7hrs post-transfection, the transfected media was aspirated out and the cells were gently washed with PBS, Trypsin treated to dislodge from the attached plate and seeded again onto PLL-treated wells (8 individual wells) of a 12-well plate (Catalog # 25-106; Genesee Scientific) at a seeding density of ~$10^5$ cells. The 12-well plate was then kept inside the incubator for at least 48 hours before using the optical imaging.

For plate reader assays, we replaced the transfection media in a 60mm dish with fresh growth media after a gentle PBS wash 7hrs post-transfection. Before performing bioluminescent spectral recordings, the attached cells were dislodged by Trypsin treatment, re-suspended in a fresh 1mL growth media and distributed equally among 5 wells of a 96-well plate (Catalog # 3912; Costar; Corning Inc.) with 200$\mu$L each.

We confirmed the functionality of the AMBER under physiological depolarisation by co-expressing capsaicin activated *rTRPV1* along with *FRP-VSD-eluxAB-YPet* and *luxCDE*. We used 500ng of *pcdNA3.1-rTRAPV1* plasmid, 250ng of *FRP-VSD-eluxAB-YPet* and 250ng of *luxCDE* for co-expressing *rTRPV1* with the bright probe and substrate proteins. The cells were seeded into a 12-well plate as before. For the patch-clamp electrophysiology experiments, cells co-expressing the bright probe and its substrate were plated onto a PLL treated 13mm coverslip (Catalog # 63780-1; #1; 0.13-0.17mm; Electron Microscopy Sciences) placed inside each well of 12-well plate at a seeding density of $10^4$ cells per well.

### 3. In vitro Bioluminescent Imaging of AMBER Sensitized to cellular depolarization

A custom imaging set up was built to record bioluminescent signals emitted by the engineered AMBER proteins. We reduced the optical path between the sample and the detector by choosing a suitable inverted microscope (Tritech IX-512) that has a fewer number of components in the light path – an objective (Motic 10X 0.3 ∞/0.17) and a removable filter cube. A sensitive EMCCD (Electron Multiplied Charged Coupled Device) camera (back-illuminated Andor iXonEM$^{+}$ 897, Model # DU-897E-CSO-#BV) was directly mounted on the camera port fitted with a C-mount adapter. Acquired raw images were digitized with 512 × 512 pixels and transferred to a computer through a PCI controller card (CCI-23). The whole imaging set up was placed inside an Aluminum Faraday cage to minimize the effect of stray electromagnetic noise on the recordings.

The EMCCD camera was operated using the open-source Micromanager software v1.4 [10]. We cooled the CCD chip to –90$^{o}$C using externally powered Peltier cooler to minimize the effect of thermal noise in the recordings. The camera has high quantum efficiency over a wide spectral range (varies between 80 and 95% between 400-700 nm). A 12-well tissue culture plate containing transfected cells attached to the bottom of each well was fixed on a manual stage that allows precise in plane positioning of the plate. We imaged cells in three different channels – bright field, fluorescence, and bioluminescence. All imaging experiments were performed at room temperature cutting off stray ambient light using Blackout fabric (black Nylon, Polyurethane-coated; Thorlabs; Part # BK5) and Aluminum foils wherever necessary.

Bright-field images were recorded at very low illumination intensity. The pre-amplifier gain was set at 1X for 10MHz readout rate to eliminate the overflow of CCD registers. The exposure time was set at 5ms and the images were recorded at an EM gain of 10. For fluorescent imaging, we used the same settings for pre-amplifier gain and the acquisition rate. However, the exposure time (10-20ms) and the EM gain (10-40) were varied between the samples depending on the expression level of the proteins.

Transient cellular expression of YPet fused probe molecules was characterized by recording its fluorescence using a long pass GFP filter (Tritech Research; MINJ-F-FITC; 480/535nm). We also recorded the endogenous NADH fluorescence in the DAPI channel (Tritech Research; MINJ-F-DAPI; 350/460nm) to characterize the concentration of $O_2$ in cells before and after depolarization [11].

We observed cell-to-cell variability in protein expression besides the batch-to-batch variation. We observed that in less than 1% of transfected cells, the probe molecules were expressed at a very high level of which a majority was internalized. More than 90% of the transfected cells in each batch expressed the probe proteins at moderate levels targeting the plasma membrane and are sensitized to bioluminescent emission under depolarization.

For bioluminescent imaging, the room was completely cut-off from the stray ambient light. We used a pre-amplifier gain of 5X and a readout rate of 3MHz. The exposure time was set at 10s for a maximum EM gain of 1000. Images were recorded before and after depolarization by KCl (Sigma; Catalog # P9541-500G) addition. Drops of KCl were added directly into the well without disturbing the field of view under focus. Bioluminescent images were recorded 10s immediately after the addition of KCl so that the cells imaged are not challenged for a longer duration. The DMEM media used contains 5mM of KCl. We accounted for this amount in our estimates of the final concentration (~ 50mM). The EMCCD camera is prone to exhibit a latency effect immediately after exposure to ambient light. We found a 10s delay allowed after KCl addition was sufficient to maintain a consistent basal noise floor without causing any latency effect in our recordings. We also recorded the endogenous NADH fluorescence before and after the KCl addition.

For modeling physiological depolarization conditions, cells co-expressing rTRPV1, bright probe and its substrate were used for bioluminescent imaging. We activated rTRPV1 by adding Capsaicin (Sigma Aldrich; Catalog # M2028-250MG) at a final concentration of 100μM and allowed a 10s delay before recording the bioluminescent signals. The necessity to achieve a large depolarization within a short duration demanded a greater amount of Capsaicin in our assay (about 10 fold greater) than reported in the literature [12].

### 4. Bioluminescent Spectral Response of AMBER emission

Bioluminescent emission spectra of AMBER were obtained using a plate reader (TECAN Spark v2.2). We recorded all emission spectra at 37$^{o}$C. HEK293 cells co-expressing the bright probe (or Dark mutant) and its substrate after 48 hours post-transfection were used for recording the bioluminescent emission spectra. Adherent cells grown in a 60mm tissue culture dish were dislodged by Trypsin treatment and suspended in 1mL of freshly prepared growth media. Suspended cells obtained from each dish were loaded into 5 wells of a dark-adapted 96 well plate (Corning Inc; Costar; Catalog # 3912) 200μL each. We dark-adapted opaque 96 well plates covering with an Aluminum foil and storing in a light cut-off dark ambiance for at least 12 hours. Dark adaptation minimizes the absorption of stray ambient light if any that are re-emitted during recordings, which affects the signal-to-noise ratio of the spectral traces. Plate containing suspended cells was loaded on the instrument tray and thermally equilibrated for about 2 minutes at 37$^{o}$C. We recorded emission spectra between 398-593nm at an interval of 15nm allowing 10s integration time at each spectral wavelength. Our experimental method aimed to minimize the long-term effects of the KCl challenge (entire spectral emission recording is completed within 2 min) on the bioluminescence recordings. We obtained paired measurements of emission spectra by recording signals before and after the addition of KCl for each well. We sampled n=8 wells to obtain the mean and standard error of photons counts at each spectral wavelength within the chosen bandwidth.

### 5. Characterizing AMBER Performance by Electrical Stimulation

We characterised the electro-optical performance of AMBER by determing the steady-state half-maximal voltage of the AMBER proteins using electrophysiology experiments. Our experimental approach involves recording bioluminescent intensities of a single cell under a whole cell voltage clamp set up. HEK293 cells co-expressing the bright probe and its substrate were used for electrophysiological measurements after 48 hours post transfection.

Isolated single cells adhered on to a 13mm PLL treated coverslip (Catalog # 63780-1; #1; 0.13-0.17mm; Electron Microsccopy Sciences) were immersed into 150μl Ringers' bath solution added into an imaging chamber (Catalog # 64-1944; MODEL # QR-41LP; Quick release chamber for 18mm coverslips; Warner Instruments). The chemical composition of the bath solution (in mM): 140 NaCl, 5 KCl, 2 $CaCl_2$, 1 $MgCl_2$, 10 glucose, and 10 HEPES at pH 7.4 (adjusted with NaOH). Recording pipettes were pulled from a micropipette glass (Catalog # BF150-86-10; Borosilicate with filament OD 1.5 mm, ID 0.86 mm, 10cm length; Sutter Instruments) to 5 - 7 MΩ. Pipettes were filled with solution containing (in mM): 125 K-Gluconate, 8 NaCl, 0.6 $MgCl_2$, 0.1 $CaCl_2$, 1 EGTA, 10 HEPES, 4 Mg-ATP, 0.4 Na-GTP and pH 7.2 (adjusted with KOH). Cells showing sufficiently detectable YPet fluorescence were selected for patching in the whole-cell mode. A stable Giga Ohm seal was formed between a patch pipette and the cell membrane by spatially controlling the position of the patch pipette using micromanipulators (PCS-PS60, 5000 Series, Burleigh). Voltage clamping at different membrane potential was achieved using Axon 200B amplifier and 1440A Digitizer (Axon™ Digidata 1550; Molecular Devices). The holding potential was stepped up from –60 mV to +60 mV in steps of +20 mV and then stepped back to –60 mV in steps of –20 mV. At each holding potential, bioluminescent signals from the patched cell was recorded using an EMCCD camera (Andor iXON$^{EM+}$ Ultra 897) with a CCD chip cooled to sub-zero temperature (–90°C) using a Peltier cooler. The camera was exposed for 10s at a 1fps acquisition rate, 3MHz readout rate and a maximum gain (1000) to record the bioluminescent signals from a single cell. We obtained paired measurements by recording the bioluminescent intensities of a patched cell for 6 independent repeats at various holding potentials within the chosen range.

### 6. In vivo Expression of AMBER in *C. elegans*

We followed the protocols for C.elegans maintenance documented earlier [13]. Transgenic animals co-expressing *FRP-VSD-eluxAB-YPet* and *luxCDE* genes from high copy extra-chromosomal arrays were created using microinjection protocols described earlier [14,15]. Briefly, plasmids encoding the bright probe and its substrate generating proteins at ~50ng/µl each along with ~100ng/µl pRF4 [*rol-6(su104)*] were injected into the distal gonads of well-fed day-1 or day-2 gravid adult hermaphrodites. We used C. *elegans* expression vectors with *mec-7* and *myo-2* promoters that drive the transgene expression in mechanosensory touch neurons and pharyngeal muscles respectively. We screened for Rol phenotype or YPet fluorescence in the transgenic F1 progeny and maintained those worms in separate plates. Transgenic animals from lines with high transmission of the extrachromosomal arrays were subsequently analyzed for expression of the bioluminescent reporters.

We selected those lines exhibiting Rol phenotype that show YPet fluorescent signal for *in vivo* bioluminescent imaging. Surprisingly, Rol phenotype animals with *myo-2* driven expression showed stronger YPet fluorescence as opposed to those with *mec-7* driven expression.

Epifluorescent images of worms expressing *mec7-eGFP* were acquired using a Nikon Eclipse Ti microscope controlled by NIS Elements AR software (Figure S8). Specimens were illuminated with an X-cite light source using a GFP filter (480/40 bandpass excitation filter). All images were obtained with a Hamamatsu CMOS sensor using a 20X objective.

### 7. Experimental Set up For Tracking Freely Moving *C. elegans* During Bioluminescent Voltage Imaging.

Limited photon budget of bioluminescent reaction demands long camera exposure times at high gain to detect the signal. Live tracking of the position and shape changes of an animal is required for spatially mapping the activity in real time. We developed an imaging approach to track the spatial information of a freely moving animal by time-gating the bioluminescent recordings in each exposure cycle. Sacrificing bioluminescent photons temporally at this gated interval (< 10ms) in each exposure cycle practically will not affect the time-integrated bioluminescent intensities.

We performed live functional imaging of *C. elegans* using an inverted microscope (TRITECH IX-512) mounted with an EMCCD camera (Andor iXON$^{EM+}$ 897). The image sensor of the camera was thermoelectrically cooled to –90$^{o}$C. Our *in vivo* imaging assay requires alternating between the bright field (low gain and small exposure time) and the bioluminescent (high gain and long exposure time) recording modes so that the shape changes and the position of the animal can be tracked simultaneously while detecting bioluminescent signal. A similar approach was used for simultaneous tracking of the neural gene expression and behavior of marmosets at a higher frame rate [16]. We accomplished this using a custom-made set up that allows our camera exposure to be synchronized with a light-emitting diode (LED) illumination during the bright field exposure. A micromanager script that instructs the camera to perform a particular exposure sequence alternating between bioluminescent and bright field modes was developed. Execution of that script delivers a trigger voltage to a microcontroller (Arduino Uno Rev.3) that drives an LED flash circuit (a 2N7000 transistor and a 330 Ohm resistor) to illuminate an LED (Adafruit Super bright white 5mm; Product ID#754) selectively during bright field image acquisition. The LED is positioned relative to the sample at a distance so there is no latency while recording the bioluminescent signals.

We exploited orders of difference in the exposure times and gains between the bright field and bioluminescent imaging modes to develop an algorithm for live tracking. Figure S8a shows the connectivity and circuit diagram between different components of the imaging hardware. The flow chart (Figure S8b) explains an algorithm applied for programming the microcontroller hardware. We start the image acquisition by running a micromanager script (see '*imcapture.bsh'* under the section Scripts and Macros) that instructs the camera to follow a programmed exposure pattern (blue for the luminescence and black for the bright field) as shown in Figure S8c. The camera exposure pattern alternates between bioluminescence and bright field modes generating trigger voltage consistent with the exposure times. We programmed the microcontroller using the prior knowledge that a short pulse always follows a long pulse (See '*MicroscopeLight.ino'* under the section Scripts and Macros). This ensures that the lighting patterns driven by the microcontroller follows synchronously with the camera trigger output (set in the '*imcapture.bsh'*). Initially, the microcontroller was in the 'idle state' waiting for a rising edge of a pulse. After encountering the rising edge of a pulse, it evaluates if the pulse width (grey color) was greater than $t_L/2$ (~ 500ms). If so, the microcontroller will switch to 'arm state' waiting for the next rising edge to drive the flash circuit. Once the rising edge of the bright field trace is detected, the LED is turned on instantaneously during the period of bright field exposure ($t_S$ < 10ms). The LED turns off once the falling edge is approached resetting the microcontroller to the 'idle-state' again to encounter the next pulse. We ignored the time for communicating between different hardware during the image acquisition because the bioluminescent photons if any, lost during this time are negligible.

### 8. Live Recording of *C. elegans* Pharyngeal Muscle Activity using AMBER

We demonstrated the in vivo use of AMBER using *C. elegans* animal model. We imaged the activities of the pharyngeal muscles using our custom-built imaging set up. Transgenic animals (1-3 days old) constitutively co-expressing the bright probe and its substrate in the pharyngeal muscles were used for these experiments. We transferred the animals on to a thin LB agar bed created on a glass coverslip (Catalog # 72210-21; Electron Microscopy Sciences) using a flame sterilized platinum wire. The agar bed was sufficiently loaded with OP50 bacteria locally for the worms to feed so that their movements are restricted within the chosen field of view. The animals were allowed to acclimatize the new environment for about 20 minutes before starting the imaging protocol.

Live bioluminescent imaging of animals was performed using our custom-built imaging set up described in the earlier section. We used two different objectives (Plan 4X 0.1 ∞/– and Motic 10X 0.3 ∞/0.17) in our imaging experiments. High expression of probe molecules enabled to use 10X objective for imaging a few animals. The coverslip loaded with the transgenic animals was positioned over the objective using a mechanical stage. The imaging room was completely light cut-off using aluminum foil sheets and black fabric. We captured 200 frames each in bioluminescent and bright field modes alternatively. The programmed exposure trace consists of two alternating exposure times at a 3MHz readout rate – 1s at a maximum gain of 1000 and 5ms at a lower gain of 20. A total of 400 frames (200 frames each containing bioluminescent and bright field images) were recorded using different magnifications. Figure S8c shows the exposure pattern used for one such experiment with $t_L$ = 1s and $t_s$ = 6ms. The raw intensity data obtained were processed further to obtain the optical characterization of the pharyngeal pumping events of animals.

### 9. Recording C. *elegans* touch neurons activities using AMBER

We used AMBER to visualize the activities of mechanosensory touch neurons in freely moving *C. elegans*. We assessed the efficacy of targeting mechanosensory touch neurons by fluorescent imaging of transgenic worms expressing the eGFP protein driven by the *mec-7* promoter (Figure S8). The *mec-7* promoter targets the touch neurons identified earlier [17] – PLM (L/R), PVM, ALM (L/R), AVM and Anterior Nerve Ring (NR). eGFP was expressed at high levels in most of the touch neurons (PLM, PVM, ALM, AVM and NR). Expression in PVD was relatively lower compared to other touch neurons. Transgenic animals (1-3 days old) constitutively expressing the bright probe and its substrate genes under the control of *the mec-7* promoter were used for the bioluminescent imaging experiments.

Our imaging approach involves tracking the spatial locations of the moving animals while recording bioluminescent signals using a custom-built imaging set up described in the earlier section. We used a 4X objective (Plan 4X 0.1 ∞/–) to achieve a greater field of view so that the movements of the worms can be tracked over a large distance. Animals moving randomly over a bacterial lawn created on an agar bed inside a 10cm dish were positioned over the objective of the inverted microscope using a mechanical stage. The imaging room was completely light cut-off and all recordings were taken at room temperature. We captured 200 frames each in bioluminescent and bright field imaging modes alternatively. Bioluminescent images were recorded exposing the image sensor for 1s at a maximum gain (1000) and a minimum readout speed (3MHz). The intervening bright field images were captured exposing the image sensor for 6ms at a very low gain (20) immediately after each bioluminescent frame. Several image stacks containing 400 frames (200 frames each with bioluminescent and bright field images) were captured to detect the activity of the touch neurons as reported under different conditions. Figure S8c shows the exposure pattern used for one such experiment with $t_L$ = 1s and $t_s$ = 6 ms. The raw intensity data obtained from the image stacks using this approach were processed to map the signatures of different touch neurons on freely moving multiple animals.

### 10. Image and Data Processing Methods

We post-processed raw bioluminescent micrographs to determine the true signal intensities. We used the open-source ImageJ software (ImageJ 1.52p) routines to post-process all the recorded images. Raw bioluminescent micrographs (512 × 512 pixels) from the in vitro cell culture experiments were binned spatially (4×4) to increase the signal-to-noise ratio of bioluminescent intensity. We found a 4×4 spatial binning was sufficient to achieve the discernable signal output without substantially losing the resolution of the morphological features spatially. The resulting 128 × 128 pixel images were background subtracted (*Process → Subtract Background*) using a rolling ball radius of 50 pixels. This was followed by the removal of bright signal outliers (*Process → Noise → Remove Outliers*) using a radius of 2 pixels and a threshold intensity of 50. The resulting images were then despeckled (*Process → Noise → Despeckle*) to remove speckle patterns if any in the field of view. The RGB images were finally digitized for 16-bit color code to compute the differential change in the bioluminescent signal intensity, $\Delta L/L$.

We confirmed detecting the spatially graded bioluminescent intensities from the transgenic animals by comparing against the background intensities of the untransfected control animals. Transgenic animals expressing the bright probe (and its substrate proteins) in the mechanosensory touch neurons were used to record 200 frames in the bioluminescent mode allowing 1s exposure for each frame without any time delay. A bright field image was recorded at the end of bioluminescent recordings to capture the traces left on the agar pad by moving animals during image acquisition. We then post-processed the signal by removing the outliers across all images in the bioluminescent stack. A moving window with time-integrated pixel intensities for 10s stepping up at an increment of 1s along the time axis was applied to process the individual frames of the bioluminescent stack. This was done using an AWK programming script (See '*slicegen.awk*' under the section Scripts and Macros), whose output generates an ImageJ macro that performs the time integration of pixel intensities (See '*lum200_11stp.ijm*' created using '*slicegen.awk*' under the section Scripts and Macros). Execution of this ImageJ script generated a modified substack with 189 frames of temporally integrated bioluminescent intensities (10s integration time) at 1s increment. We then applied mean filtering to the pixel intensities keeping the threshold at the maximum background intensity value (varied in each recording but never exceed a value of 3475 arb. units). Spatially graded bioluminescent intensities at different time points co-occur at certain specific locations on the discrete traces left on the agar pad by the animals. These locations presumably agreed with the soma locations of mechanosensory touch neurons mapped previously. Similar recordings from untrasfected control animals did not give any detectable signals for the chosen threshold intensity values. The actual signal intensity recorded from the transgenic worms varied spatially at different time instants.

We developed an image processing protocol to generate a movie that shows live spatial tracking of bioluminescent intensities in a freely moving animal. Image stacks containing bioluminescence and brightfield frames alternatively were post-processed to create animated movies. Image stacks were segregated into brightfield and luminescent substacks using ImageJ built-in function (*Image → Stacks → Tools → Make Substack*). We used different ranges in the substack maker to create these two different substacks – (1-399-2) for bioluminescence and (2-400-2) for brightfield substacks. Execution of an ImageJ script ('*lum200_11stp.ijm*') created a modified substack with 189 frames of temporally integrated bioluminescent intensities (10s integration time) at 1s increment. The modified substack was background subtracted followed by setting an intensity threshold up to the maximum of background intensity to tease out the spatially graded bioluminescent signals at different time points. We then mapped the intensity contour to the spatially binned (4×4) brightfield stack using an ImageJ plugin (Stack Interleaver) developed earlier [18]. This method was applied to spatially map the bioluminescent intensities of freely moving transgenic animals to the corresponding brightfield image recorded at that instant.

Movies showing mechanosensory touch neurons activities of animals were created cropping a window that includes a complete field of view of the motion trajectories. We observed spatial distribution of bioluminescent intensities representing different levels of activity in the moving animals. However, the number of pixels associated with the fractional change of the AMBER intensities varied spatiotemporally at the predetermined locations. This could be either due to signal retention effects of time integration or the spatially restricted movements displayed by the animals.

We characterized the voltage activity of the pharyngeal muscles in freely unrestrained individual animals. We observed a weak background signal from the embryos within young hermaphrodite adults, which was confirmed both from the fluorescent and bioluminescent signals. Live animal tracking protocol discussed before was used to create movies showing pharyngeal muscle activities of young adults for a few minutes. Both bright field and bioluminescent intensities of the terminal bulb were obtained to understand the relation between the voltage activity and the muscle movements. High-frequency signals observed from the bioluminescent traces precisely detected the contraction and relaxation kinetics of the muscle movement.

We used MATLAB programming to process the raw bioluminescent voltage signals obtained from the Corpus, Isthmus and Terminal Blub of individual animals. N2 animals typically exhibit slow pumping frequencies (0.5-1Hz). Low occurrence of fast pumping frequencies (> 1.5Hz) normally requires serotonin neural inputs [19]. Earlier work showed evidences for the maximization of instantaneous pumping rates at about ~1Hz [20] when sufficient food is available. We therefore applied band-pass filtering to the raw data allowing low frequency voltage signals (mostly in the range of 0.08-0.8Hz depending on the SNR of the traces) at a sampling rate of 2Hz (See '*pharyngeal_processing.m*' under the Section 'Scripts and Macors').

### 11. Statistical Methods

All experiments were performed unblinded.

We observed variability in the in vitro expression of engineered AMBER protein constructs among cells in a batch and between the batches. A suitable metric was identified for comparing the performance (fractional luminescence, $R = \Delta L/L$) of various AMBER proteins in cellular depolarization assays. In principle, when cells light up sufficiently bright and can be discerned against their background, then the pixels encoding the differential intensity should describe the actual morphology of the cells as observed under brightfield illumination. We defined a metric to quantitate information transmission, $I_R$ as a ratio of the Shannon information [21] encoded in the pixels representing differential bioluminescent intensity to that of the bright field intensity. Mathematically, $I_R$ is expressed as

$$I_R = 2^{(Hd-Hbf)} \qquad \text{(S1)}$$

where $H_d$ and $H_{bf}$ represents Shannon entropies of the images encoding differential bioluminescence and bright field intensities respectively. We used 8-bit post-processed bioluminescent micrographs to evaluate Shannon entropies using the in-built MATLAB (R2019a, Update 1) function, *entropy* [22]. Given that Shannon entropy quantifies the information density of an image, we estimated the information transmission accompanied by cellular bioluminescence after KCl challenge by computing $I_R$ (See Table S2).

We performed a paired two-tailed Student's t-test to confirm there is a statistically significant difference in the mean intensity of the cells after KCl challenge. We defined our null hypothesis, $H_0$: No statistically significant difference between the mean intensities of the cells before and after KCl addition. The alternate hypothesis, $H_1$ was stated as statistically significant difference exist in the mean intensities of the cells after KCl addition. We ensured the number of cells associated with the chosen pixels were sufficiently high (> at least 40 cells) to provide the necessary statistical power for the hypothesis test. We used pixel-by-pixel intensity difference to compute the test statistic, which was higher than the critical value by a large difference for all the engineered constructs (See Table S3).

For KCl titration assay, 8 different populations of cells were chosen randomly using a $100\mu m \times 100\mu m$ window. We measured an increase in the bioluminescent intensity at those locations after subtracting the background intensities in their vicinity locally for an incremental addition of KCl into the bath. Assuming stable conformational states of the voltage sensor as being described as a two state model [23], the total intensity for any particular KCl concentration was obtained as a cumulative sum of the intensity values of all the previous steps. Mean and standard error of the intensities were computed from the intensity values recorded at the chosen window locations.

Emission spectra obtained from the plate reader experiments were processed from the raw data collected from several samples (n=8). Representative emission spectra at a spectral wavelength resolution of 15 nm were plotted (Figure 2c and Figure S5). We reported the average spectral intensity of 8 independent trials before and after KCl addition. We performed hypothesis test applying unpaired 2-sample two-tailed t-test at 95% confidence ($\alpha$ = 0.5) to assess if the differential spectral intensity after KCl challenge is statistically significant. The computed t-test statistic (~5.52) exceeded the critical value (~2.02) for the associated degrees of freedom (40) confirming the mean spectral intensities of the samples obtained before and after KCl addition are different ($p < 0.0005$).

Representative whole cell voltage clamp recording report the statistical variation of 6 independent measurements between –60mV to 60mV at an increment of 20mV from a stable single cell patch. Mean and standard error of bioluminescent intensities evaluated from the raw data follow the Boltzmann model as observed in the KCl titration assay. We conducted a paired t-test between two states – inactive (–60mV) and the active (20mV) states at 95% confidence level ($\alpha = 0.5$) to determine if the differential single cell intensities between these states are statistically significant. The computed t-test statistic ($t_{st}$~2.9) exceeded the threshold value ($t_{th}$~2.0) suggesting there is likelihood ($p < 0.02$) for the estimated means to be different.

We confirmed the functionality of the bright probe in vivo based on the optical readouts of the AMBER characterising the activities of the pharyngeal muscles and mechanosensory neural circuit. We observed the activities reported by AMBER from several animals (at least n=5) during each recording from both in vivo assays performed.

### 12. Scripts and Macros

**Micromanager script to instruct the EMCCD camera to record bioluminescent and brightfield images alternatively**

**imcapture.bsh**

```
acqName="testacq_01";
rootDirName="C:/Users";
gui.closeAllAcquisitions();
gui.clearMessageWindow();
cameraName=mmc.getCameraDevice();
numFrames=200;
gui.openAcquisition(acqName,rootDirName,numFrames,1,1);
hex=1000;
lex=6;
for (int i=0; i<numFrames; i++)
{
	mmc.setProperty(cameraName, "Gain", "1000");
	mmc.setExposure(hex);
	gui.snapAndAddImage(acqName,(2*i+1),0,0,0);
	mmc.setProperty(cameraName, "Gain", "20");
	mmc.setExposure(lex);
	gui.snapAndAddImage(acqName,(2*i+2),0,0,0);

}
```

**Microcontroller programming for driving flash circuit**

**MicroscopeLight.ino**

---

```
int sensorIn = 5; // For now set as Pin 5 of the arduino
int ledon = 7;    // Output pin to hook to the LED
int sensorRead = 0;
int i;
int time_small = 10;
int time_big = 100;
int time_wait = 20;

void setup() {
// put your setup code here, to run once:
pinMode(sensorIn, INPUT);
pinMode(ledon, OUTPUT);
pinMode(LED_BUILTIN, OUTPUT);

}

void loop() {
// put your main code here, to run repeatedly:
sensorRead = digitalRead(sensorIn);
if(sensorRead == HIGH)
{
for( i=0;i<(time_big/2)+1;i++)
{
delay(1);      // Delay of time_big/2 ms in total
}
sensorRead = digitalRead(sensorIn);
if (sensorRead == HIGH)
{
while( digitalRead(sensorIn) == HIGH)
{
delay(1);
}
for( i=0;i<(time_wait/2);i++)
{
delay(1);      // Delay of time_big/2 ms in total
}
while( digitalRead(sensorIn) == LOW)
{
delayMicroseconds(100);
}
digitalWrite(LED_BUILTIN,HIGH);
digitalWrite(ledon,HIGH);
while( digitalRead(sensorIn) == HIGH)
{
delayMicroseconds(100);
}
digitalWrite(LED_BUILTIN,LOW);
digitalWrite(ledon,LOW);
}
}
}
```

**AWK program for generating ImageJ macro ‘lum200_11stp.ijm’ to process image stacks**

**slicegen.awk**

```
# slicegen.awk for generating ImageJ macro lum200_11stp.ijm

#do while statement
BEGIN {
#initialize a counter
x=0
for (i=1;i<=189; i++)
{
   printf("selectWindow(\"Lum\")\n");
   printf("run(\"Z Project...\", \"start=" i" " "stop="i+10" " "projection =[Sum Slices]\")\n");
}
}
```

**ImageJ script for processing stacks recorded in the luminescent mode**

**lum200_11stp.ijm**

---

```
selectWindow("Lum")
run("Z Project...", "start=1 stop=11 projection =[Sum Slices]")
selectWindow("Lum")
run("Z Project...", "start=2 stop=12 projection =[Sum Slices]")
selectWindow("Lum")
run("Z Project...", "start=3 stop=13 projection =[Sum Slices]")
selectWindow("Lum")
run("Z Project...", "start=4 stop=14 projection =[Sum Slices]")
selectWindow("Lum")
run("Z Project...", "start=5 stop=15 projection =[Sum Slices]")
selectWindow("Lum")
run("Z Project...", "start=6 stop=16 projection =[Sum Slices]")
selectWindow("Lum")
run("Z Project...", "start=7 stop=17 projection =[Sum Slices]")
selectWindow("Lum")
run("Z Project...", "start=8 stop=18 projection =[Sum Slices]")
selectWindow("Lum")
run("Z Project...", "start=9 stop=19 projection =[Sum Slices]")
selectWindow("Lum")
run("Z Project...", "start=10 stop=20 projection =[Sum Slices]")
selectWindow("Lum")
run("Z Project...", "start=11 stop=21 projection =[Sum Slices]")
selectWindow("Lum")
run("Z Project...", "start=12 stop=22 projection =[Sum Slices]")
selectWindow("Lum")
run("Z Project...", "start=13 stop=23 projection =[Sum Slices]")
selectWindow("Lum")
run("Z Project...", "start=14 stop=24 projection =[Sum Slices]")
selectWindow("Lum")
run("Z Project...", "start=15 stop=25 projection =[Sum Slices]")
selectWindow("Lum")
run("Z Project...", "start=16 stop=26 projection =[Sum Slices]")
selectWindow("Lum")
run("Z Project...", "start=17 stop=27 projection =[Sum Slices]")
selectWindow("Lum")
run("Z Project...", "start=18 stop=28 projection =[Sum Slices]")
selectWindow("Lum")
run("Z Project...", "start=19 stop=29 projection =[Sum Slices]")
selectWindow("Lum")
run("Z Project...", "start=20 stop=30 projection =[Sum Slices]")
selectWindow("Lum")
run("Z Project...", "start=21 stop=31 projection =[Sum Slices]")
selectWindow("Lum")
run("Z Project...", "start=22 stop=32 projection =[Sum Slices]")
selectWindow("Lum")
run("Z Project...", "start=23 stop=33 projection =[Sum Slices]")
selectWindow("Lum")
run("Z Project...", "start=24 stop=34 projection =[Sum Slices]")
selectWindow("Lum")
run("Z Project...", "start=25 stop=35 projection =[Sum Slices]")
selectWindow("Lum")
run("Z Project...", "start=26 stop=36 projection =[Sum Slices]")
selectWindow("Lum")
run("Z Project...", "start=27 stop=37 projection =[Sum Slices]")
selectWindow("Lum")
run("Z Project...", "start=28 stop=38 projection =[Sum Slices]")
selectWindow("Lum")
run("Z Project...", "start=29 stop=39 projection =[Sum Slices]")
selectWindow("Lum")
run("Z Project...", "start=30 stop=40 projection =[Sum Slices]")
selectWindow("Lum")
run("Z Project...", "start=31 stop=41 projection =[Sum Slices]")
selectWindow("Lum")
run("Z Project...", "start=32 stop=42 projection =[Sum Slices]")
selectWindow("Lum")
run("Z Project...", "start=33 stop=43 projection =[Sum Slices]")
selectWindow("Lum")
run("Z Project...", "start=34 stop=44 projection =[Sum Slices]")
selectWindow("Lum")
run("Z Project...", "start=35 stop=45 projection =[Sum Slices]")
selectWindow("Lum")
```

```
run("Z Project...", "start=36 stop=46 projection =[Sum Slices]")
selectWindow("Lum")
run("Z Project...", "start=37 stop=47 projection =[Sum Slices]")
selectWindow("Lum")
run("Z Project...", "start=38 stop=48 projection =[Sum Slices]")
selectWindow("Lum")
run("Z Project...", "start=39 stop=49 projection =[Sum Slices]")
selectWindow("Lum")
run("Z Project...", "start=40 stop=50 projection =[Sum Slices]")
selectWindow("Lum")
run("Z Project...", "start=41 stop=51 projection =[Sum Slices]")
selectWindow("Lum")
run("Z Project...", "start=42 stop=52 projection =[Sum Slices]")
selectWindow("Lum")
run("Z Project...", "start=43 stop=53 projection =[Sum Slices]")
selectWindow("Lum")
run("Z Project...", "start=44 stop=54 projection =[Sum Slices]")
selectWindow("Lum")
run("Z Project...", "start=45 stop=55 projection =[Sum Slices]")
selectWindow("Lum")
run("Z Project...", "start=46 stop=56 projection =[Sum Slices]")
selectWindow("Lum")
run("Z Project...", "start=47 stop=57 projection =[Sum Slices]")
selectWindow("Lum")
run("Z Project...", "start=48 stop=58 projection =[Sum Slices]")
selectWindow("Lum")
run("Z Project...", "start=49 stop=59 projection =[Sum Slices]")
selectWindow("Lum")
run("Z Project...", "start=50 stop=60 projection =[Sum Slices]")
selectWindow("Lum")
run("Z Project...", "start=51 stop=61 projection =[Sum Slices]")
selectWindow("Lum")
run("Z Project...", "start=52 stop=62 projection =[Sum Slices]")
selectWindow("Lum")
run("Z Project...", "start=53 stop=63 projection =[Sum Slices]")
selectWindow("Lum")
run("Z Project...", "start=54 stop=64 projection =[Sum Slices]")
selectWindow("Lum")
run("Z Project...", "start=55 stop=65 projection =[Sum Slices]")
selectWindow("Lum")
run("Z Project...", "start=56 stop=66 projection =[Sum Slices]")
selectWindow("Lum")
run("Z Project...", "start=57 stop=67 projection =[Sum Slices]")
selectWindow("Lum")
run("Z Project...", "start=58 stop=68 projection =[Sum Slices]")
selectWindow("Lum")
run("Z Project...", "start=59 stop=69 projection =[Sum Slices]")
selectWindow("Lum")
run("Z Project...", "start=60 stop=70 projection =[Sum Slices]")
selectWindow("Lum")
run("Z Project...", "start=61 stop=71 projection =[Sum Slices]")
selectWindow("Lum")
run("Z Project...", "start=62 stop=72 projection =[Sum Slices]")
selectWindow("Lum")
run("Z Project...", "start=63 stop=73 projection =[Sum Slices]")
selectWindow("Lum")
run("Z Project...", "start=64 stop=74 projection =[Sum Slices]")
selectWindow("Lum")
run("Z Project...", "start=65 stop=75 projection =[Sum Slices]")
selectWindow("Lum")
run("Z Project...", "start=66 stop=76 projection =[Sum Slices]")
selectWindow("Lum")
run("Z Project...", "start=67 stop=77 projection =[Sum Slices]")
selectWindow("Lum")
run("Z Project...", "start=68 stop=78 projection =[Sum Slices]")
selectWindow("Lum")
run("Z Project...", "start=69 stop=79 projection =[Sum Slices]")
selectWindow("Lum")
run("Z Project...", "start=70 stop=80 projection =[Sum Slices]")
selectWindow("Lum")
run("Z Project...", "start=71 stop=81 projection =[Sum Slices]")
selectWindow("Lum")
run("Z Project...", "start=72 stop=82 projection =[Sum Slices]")
selectWindow("Lum")
run("Z Project...", "start=73 stop=83 projection =[Sum Slices]")
```

```
selectWindow("Lum")
run("Z Project...", "start=74 stop=84 projection =[Sum Slices]")
selectWindow("Lum")
run("Z Project...", "start=75 stop=85 projection =[Sum Slices]")
selectWindow("Lum")
run("Z Project...", "start=76 stop=86 projection =[Sum Slices]")
selectWindow("Lum")
run("Z Project...", "start=77 stop=87 projection =[Sum Slices]")
selectWindow("Lum")
run("Z Project...", "start=78 stop=88 projection =[Sum Slices]")
selectWindow("Lum")
run("Z Project...", "start=79 stop=89 projection =[Sum Slices]")
selectWindow("Lum")
run("Z Project...", "start=80 stop=90 projection =[Sum Slices]")
selectWindow("Lum")
run("Z Project...", "start=81 stop=91 projection =[Sum Slices]")
selectWindow("Lum")
run("Z Project...", "start=82 stop=92 projection =[Sum Slices]")
selectWindow("Lum")
run("Z Project...", "start=83 stop=93 projection =[Sum Slices]")
selectWindow("Lum")
run("Z Project...", "start=84 stop=94 projection =[Sum Slices]")
selectWindow("Lum")
run("Z Project...", "start=85 stop=95 projection =[Sum Slices]")
selectWindow("Lum")
run("Z Project...", "start=86 stop=96 projection =[Sum Slices]")
selectWindow("Lum")
run("Z Project...", "start=87 stop=97 projection =[Sum Slices]")
selectWindow("Lum")
run("Z Project...", "start=88 stop=98 projection =[Sum Slices]")
selectWindow("Lum")
run("Z Project...", "start=89 stop=99 projection =[Sum Slices]")
selectWindow("Lum")
run("Z Project...", "start=90 stop=100 projection =[Sum Slices]")
selectWindow("Lum")
run("Z Project...", "start=91 stop=101 projection =[Sum Slices]")
selectWindow("Lum")
run("Z Project...", "start=92 stop=102 projection =[Sum Slices]")
selectWindow("Lum")
run("Z Project...", "start=93 stop=103 projection =[Sum Slices]")
selectWindow("Lum")
run("Z Project...", "start=94 stop=104 projection =[Sum Slices]")
selectWindow("Lum")
run("Z Project...", "start=95 stop=105 projection =[Sum Slices]")
selectWindow("Lum")
run("Z Project...", "start=96 stop=106 projection =[Sum Slices]")
selectWindow("Lum")
run("Z Project...", "start=97 stop=107 projection =[Sum Slices]")
selectWindow("Lum")
run("Z Project...", "start=98 stop=108 projection =[Sum Slices]")
selectWindow("Lum")
run("Z Project...", "start=99 stop=109 projection =[Sum Slices]")
selectWindow("Lum")
run("Z Project...", "start=100 stop=110 projection =[Sum Slices]")
selectWindow("Lum")
run("Z Project...", "start=101 stop=111 projection =[Sum Slices]")
selectWindow("Lum")
run("Z Project...", "start=102 stop=112 projection =[Sum Slices]")
selectWindow("Lum")
run("Z Project...", "start=103 stop=113 projection =[Sum Slices]")
selectWindow("Lum")
run("Z Project...", "start=104 stop=114 projection =[Sum Slices]")
selectWindow("Lum")
run("Z Project...", "start=105 stop=115 projection =[Sum Slices]")
selectWindow("Lum")
run("Z Project...", "start=106 stop=116 projection =[Sum Slices]")
selectWindow("Lum")
run("Z Project...", "start=107 stop=117 projection =[Sum Slices]")
selectWindow("Lum")
run("Z Project...", "start=108 stop=118 projection =[Sum Slices]")
selectWindow("Lum")
run("Z Project...", "start=109 stop=119 projection =[Sum Slices]")
selectWindow("Lum")
run("Z Project...", "start=110 stop=120 projection =[Sum Slices]")
selectWindow("Lum")
```

```
run("Z Project...", "start=111 stop=121 projection =[Sum Slices]")
selectWindow("Lum")
run("Z Project...", "start=112 stop=122 projection =[Sum Slices]")
selectWindow("Lum")
run("Z Project...", "start=113 stop=123 projection =[Sum Slices]")
selectWindow("Lum")
run("Z Project...", "start=114 stop=124 projection =[Sum Slices]")
selectWindow("Lum")
run("Z Project...", "start=115 stop=125 projection =[Sum Slices]")
selectWindow("Lum")
run("Z Project...", "start=116 stop=126 projection =[Sum Slices]")
selectWindow("Lum")
run("Z Project...", "start=117 stop=127 projection =[Sum Slices]")
selectWindow("Lum")
run("Z Project...", "start=118 stop=128 projection =[Sum Slices]")
selectWindow("Lum")
run("Z Project...", "start=119 stop=129 projection =[Sum Slices]")
selectWindow("Lum")
run("Z Project...", "start=120 stop=130 projection =[Sum Slices]")
selectWindow("Lum")
run("Z Project...", "start=121 stop=131 projection =[Sum Slices]")
selectWindow("Lum")
run("Z Project...", "start=122 stop=132 projection =[Sum Slices]")
selectWindow("Lum")
run("Z Project...", "start=123 stop=133 projection =[Sum Slices]")
selectWindow("Lum")
run("Z Project...", "start=124 stop=134 projection =[Sum Slices]")
selectWindow("Lum")
run("Z Project...", "start=125 stop=135 projection =[Sum Slices]")
selectWindow("Lum")
run("Z Project...", "start=126 stop=136 projection =[Sum Slices]")
selectWindow("Lum")
run("Z Project...", "start=127 stop=137 projection =[Sum Slices]")
selectWindow("Lum")
run("Z Project...", "start=128 stop=138 projection =[Sum Slices]")
selectWindow("Lum")
run("Z Project...", "start=129 stop=139 projection =[Sum Slices]")
selectWindow("Lum")
run("Z Project...", "start=130 stop=140 projection =[Sum Slices]")
selectWindow("Lum")
run("Z Project...", "start=131 stop=141 projection =[Sum Slices]")
selectWindow("Lum")
run("Z Project...", "start=132 stop=142 projection =[Sum Slices]")
selectWindow("Lum")
run("Z Project...", "start=133 stop=143 projection =[Sum Slices]")
selectWindow("Lum")
run("Z Project...", "start=134 stop=144 projection =[Sum Slices]")
selectWindow("Lum")
run("Z Project...", "start=135 stop=145 projection =[Sum Slices]")
selectWindow("Lum")
run("Z Project...", "start=136 stop=146 projection =[Sum Slices]")
selectWindow("Lum")
run("Z Project...", "start=137 stop=147 projection =[Sum Slices]")
selectWindow("Lum")
run("Z Project...", "start=138 stop=148 projection =[Sum Slices]")
selectWindow("Lum")
run("Z Project...", "start=139 stop=149 projection =[Sum Slices]")
selectWindow("Lum")
run("Z Project...", "start=140 stop=150 projection =[Sum Slices]")
selectWindow("Lum")
run("Z Project...", "start=141 stop=151 projection =[Sum Slices]")
selectWindow("Lum")
run("Z Project...", "start=142 stop=152 projection =[Sum Slices]")
selectWindow("Lum")
run("Z Project...", "start=143 stop=153 projection =[Sum Slices]")
selectWindow("Lum")
run("Z Project...", "start=144 stop=154 projection =[Sum Slices]")
selectWindow("Lum")
run("Z Project...", "start=145 stop=155 projection =[Sum Slices]")
selectWindow("Lum")
run("Z Project...", "start=146 stop=156 projection =[Sum Slices]")
selectWindow("Lum")
run("Z Project...", "start=147 stop=157 projection =[Sum Slices]")
selectWindow("Lum")
run("Z Project...", "start=148 stop=158 projection =[Sum Slices]")
```

```
selectWindow("Lum")
run("Z Project...", "start=149 stop=159 projection =[Sum Slices]")
selectWindow("Lum")
run("Z Project...", "start=150 stop=160 projection =[Sum Slices]")
selectWindow("Lum")
run("Z Project...", "start=151 stop=161 projection =[Sum Slices]")
selectWindow("Lum")
run("Z Project...", "start=152 stop=162 projection =[Sum Slices]")
selectWindow("Lum")
run("Z Project...", "start=153 stop=163 projection =[Sum Slices]")
selectWindow("Lum")
run("Z Project...", "start=154 stop=164 projection =[Sum Slices]")
selectWindow("Lum")
run("Z Project...", "start=155 stop=165 projection =[Sum Slices]")
selectWindow("Lum")
run("Z Project...", "start=156 stop=166 projection =[Sum Slices]")
selectWindow("Lum")
run("Z Project...", "start=157 stop=167 projection =[Sum Slices]")
selectWindow("Lum")
run("Z Project...", "start=158 stop=168 projection =[Sum Slices]")
selectWindow("Lum")
run("Z Project...", "start=159 stop=169 projection =[Sum Slices]")
selectWindow("Lum")
run("Z Project...", "start=160 stop=170 projection =[Sum Slices]")
selectWindow("Lum")
run("Z Project...", "start=161 stop=171 projection =[Sum Slices]")
selectWindow("Lum")
run("Z Project...", "start=162 stop=172 projection =[Sum Slices]")
selectWindow("Lum")
run("Z Project...", "start=163 stop=173 projection =[Sum Slices]")
selectWindow("Lum")
run("Z Project...", "start=164 stop=174 projection =[Sum Slices]")
selectWindow("Lum")
run("Z Project...", "start=165 stop=175 projection =[Sum Slices]")
selectWindow("Lum")
run("Z Project...", "start=166 stop=176 projection =[Sum Slices]")
selectWindow("Lum")
run("Z Project...", "start=167 stop=177 projection =[Sum Slices]")
selectWindow("Lum")
run("Z Project...", "start=168 stop=178 projection =[Sum Slices]")
selectWindow("Lum")
run("Z Project...", "start=169 stop=179 projection =[Sum Slices]")
selectWindow("Lum")
run("Z Project...", "start=170 stop=180 projection =[Sum Slices]")
selectWindow("Lum")
run("Z Project...", "start=171 stop=181 projection =[Sum Slices]")
selectWindow("Lum")
run("Z Project...", "start=172 stop=182 projection =[Sum Slices]")
selectWindow("Lum")
run("Z Project...", "start=173 stop=183 projection =[Sum Slices]")
selectWindow("Lum")
run("Z Project...", "start=174 stop=184 projection =[Sum Slices]")
selectWindow("Lum")
run("Z Project...", "start=175 stop=185 projection =[Sum Slices]")
selectWindow("Lum")
run("Z Project...", "start=176 stop=186 projection =[Sum Slices]")
selectWindow("Lum")
run("Z Project...", "start=177 stop=187 projection =[Sum Slices]")
selectWindow("Lum")
run("Z Project...", "start=178 stop=188 projection =[Sum Slices]")
selectWindow("Lum")
run("Z Project...", "start=179 stop=189 projection =[Sum Slices]")
selectWindow("Lum")
run("Z Project...", "start=180 stop=190 projection =[Sum Slices]")
selectWindow("Lum")
run("Z Project...", "start=181 stop=191 projection =[Sum Slices]")
selectWindow("Lum")
run("Z Project...", "start=182 stop=192 projection =[Sum Slices]")
selectWindow("Lum")
run("Z Project...", "start=183 stop=193 projection =[Sum Slices]")
selectWindow("Lum")
run("Z Project...", "start=184 stop=194 projection =[Sum Slices]")
selectWindow("Lum")
run("Z Project...", "start=185 stop=195 projection =[Sum Slices]")
selectWindow("Lum")
```

```
run("Z Project...", "start=186 stop=196 projection =[Sum Slices]")
selectWindow("Lum")
run("Z Project...", "start=187 stop=197 projection =[Sum Slices]")
selectWindow("Lum")
run("Z Project...", "start=188 stop=198 projection =[Sum Slices]")
selectWindow("Lum")
run("Z Project...", "start=189 stop=199 projection =[Sum Slices]")
```

**MATLAB code for processing the in vivo voltage recording of the Pharyngeal muscles**

**Pharyngeal_processing.m**

---

```
close all; %Close all variables
clear all; %Clear all data
Pharynx_data = readtable('Worm2_data2.txt'); %Read Data
Muscle_data = readtable('Pharyngeal_muscle_data.txt'); %Read Data

TimeX =0.5.*Pharynx_data.Frame; %Define Time Step as 500ms
Time=TimeX(20:190);

TB=Pharynx_data.TB8x8(20:190); % Terminal Bulb

Isthmus=Pharynx_data.Isthmus5x5(20:190); % Isthmus

Corpus=Pharynx_data.Corpus8x8(20:190); % Corpus

Muscle =Pharynx_data.Muscle_mov(20:190); % Muscle Data Brightfield

FpassTB=[0.0845,0.69]; % Define band-pass filter using built-in Matlab Function Y =
bandpass(X,Fpass,Fs) specifies Fs as a positive numeric scalar
                % corresponding to the sample rate of X in Units of Hertz. Fpass = [FpassLower,
FpassUpper], is a two element vector that defines the
                % bandwidth frequency range of the filter in Units of Hertz.
FpassCorp=[0.0845,0.69];
FpassIsth=[0.0845,0.69];
FpassMus =[0.1,0.76];

slope =0.75;% Define the sharpness of bandpass filter
TB_bp=bandpass(TB,FpassTB,2,'Steepness',slope); %Filter the data
Isthmus_bp=bandpass(Isthmus,FpassIsth,2,'Steepness',slope);
Corpus_bp=bandpass(Corpus,FpassCorp,2,'Steepness',slope);
Mus_bp=bandpass(Muscle,FpassMus,2,'Steepness',slope);

TB_bp_norm=normalize(TB_bp,'range');%Normalize the result
Isthmus_bp_norm=normalize(Isthmus_bp,'range');
Corpus_bp_norm=normalize(Corpus_bp,'range');
Mus_bp_norm=normalize(Mus_bp,'range');
figure(1);
plot(Time,TB_bp_norm+2.2,Time,Isthmus_bp_norm+1.1,Time,Corpus_bp_norm+0.15,Time,Mus_bp_
norm+3.2,'Linewidth',2);% Plot the data with an offset for easy visualization
```

---

### 13. Western Blot

We estimated the molecular weights of engineered AMBER protein constructs by Western blot. We transfected $1\times10^6$ HEK293 cells with plasmid DNAs expressing engineered AMBER proteins and plated them into a 60mm tissue culture dish. After 48 hours post-transfection, the culture media was aspirated out and the cells were gently washed with 1mL of 1X PBS. The washed cells were then lysed by gently agitating in lysis buffer (150mM NaCl, 0.1% Triton X-100, 0.5% Sodium deoxycholate, 0.1% Sodium dodecyl sulfate, 50mM Tris-HCl, pH 8.0 with 1 Roche protease inhibitor tablet) for 30 minutes at 4°C. The crude lysate was centrifuged at 10000g for 15 min at 4°C to separate the soluble and insoluble fractions. Protein samples for gel electrophoresis were prepared by mixing 25$\mu$L of the soluble fraction with 25$\mu$l of Laemmli 2X sample buffer (Catalog# 161-0737; BIO-RAD) followed by denaturation by heating up to 70°C for 5 minutes. All heated samples were loaded into individual wells of a precast protein gel (4-15% BIO-RAD Mini-PROTEAN TGX Gels; Catalog # 456-1084) and protein molecules of different molecular weights were electrophoretically separated (120V for 1 hour not exceeding 86mA) in a bath of running buffer (MOPS SDS, 25mM Tris base, 190mM Glycine, 0.1% SDS, pH 8.5). The protein impregnated gel was carefully removed from the cassette and placed on a 5cm x 5cm nitrocellulose membrane (Catalog#162-0112; 0.2$\mu$m; BIO-RAD) with a filter paper (Catalog# 1703965, BIO-RAD) backing submerged inside a bath of transfer buffer (25mM Tris base, 190mM Glycine, 20% methanol, pH 8.3). The gel-membrane pair sandwiched by the filter paper on either side was placed in contact with an ice pack inside a Mini Trans-Blot cell (Serial # 153 BR 76868; BIO-RAD). Electrically driven (~ 90V) wet protein transfer onto the sandwiched membrane was accomplished in a temperature regulated transfer bath. The membrane containing transferred proteins was then blocked using a blocking buffer (LI-COR Odyssey blocking buffer (PBS); Part # 927-40000) overnight at 4°C and washed with TBST buffer (20mM Tris, 150mM NaCl, 0.1% Tween 20, pH 7.5) for 15 minutes thrice. We stained for the YPet using a primary antibody that specifically targets the epitope of most GFP variants (Monoclonal anti-GFP IgG2a raised in the mouse; Invitrogen; Catalog # A-11120). The primary antibody at 1:1000 was added directly into the treatment bath (a mixture of 7mL of TBST and 3mL of blocking buffer) containing the membrane and was incubated at room temperature on a rocking shaker (BR2000 2D rocker; Benchmark Scientific Inc.) for 1hr. The non-specific binding of the primary antibody was removed by washing the membrane thrice for 15 minutes with 10mL of TBST at room temperature on the rocking shaker. This was immediately followed by secondary antibody treatment (IRDye® 680LT Goat anti-Mouse IgG; *LI-COR*; Catalog # 926-68020) at 1:10000 for 1hr at room temperature and subsequently washing with TBST thrice as before. The secondary antibody is sensitized to a fluorescent emission at 700 nm, which is detected using a membrane-imaging system (Odyssey CLx, *LI-COR*). Protein bands emitting fluorescent signals were then processed using Image Studio software (version 5.2).

### References


1. Griesbeck, O., Baird, G. S., Campbell, R. E., Zacharias, D. A., and Tsien, R. Y. Reducing the environmental sensitivity of yellow fluorescent protein: Mechanism and Applications, *J. Biol. Chem.* **276**, 29188-94 (2001).
2. Nguyen A. W. & Daugherty, P. S. Evolutionary optimization of fluorescent proteins for intracellular FRET. *Nat. biotech*. **23**, 355–360 (2005).
3. Fisher, A. J. et al. Three-dimensional structure of bacterial luciferase from Vibrio *harveyi* at 2.4A$^{o}$ resolution. *Biochemistry*. **34**, 6581-6586 (1995)
4. Baldwin, T.O. et al. Structure of bacterial luciferase. *Curr. Opin. Struct. Biol.* **5**, 798-809 (1995).
5. Olsson, O. et al. Engineering of monomeric bacterial luciferases by fusion of *luxA* and *luxB* genes in Vibrio *harveyi*. *Gene*. **81**, 335-347 (1989).
6. Dimitrov, D. et al. Engineering and characterization of an enhanced fluorescent protein voltage sensor, *PLoS ONE*. **2**, e440 (2007).
7. Cui, B. et al. Engineering an enhanced, thermostable, monomeric bacterial luciferase gene as a reporter in plant protoplasts. *PLoS ONE*. **9**, e107885 (2014).
8. Olsson, O. et al. Engineering of monomeric bacterial luciferases by fusion of *luxA* and *luxB* genes in Vibrio *harveyi*. *Gene*. **81**, 335-347 (1989).
9. Szymczak, A. L. et al. Correction of multi-gene deficiency invivo using a single 'self-cleaving' 2A peptide-based retroviral vector. *Nat. Biotech*. **22**, 589–594 (2004)
10. Edelstein, A.D., Tsuchida, M. A., Amodaj, N., Pinkard, H., Vale, R. D. & Stuurman, N. Advanced methods of microscope control using μManager software. *J. Biol. Methods*. **1,** e10 **(**2014)
11. Schaefer, P. M., Kalinina, S., Rueck, A., von Arnim, C. A. F. & von Einem, B. NADH Autoflluorescence – A marker on its way to boost bioenergetic research. *Cytometry A*. **95A**, 34-46 (2019).
12. Cao, E., Cordero-Morales, J. F., Liu, B., Qin, F. & Julius, D. TRPV1 channels are intrinsically heat sensitive and negatively regulated by phosphoinositide lipids, *Neuron*. **77**, 667-679 (2013).
13. Brenner, S. The genetics of Caenorhabditis elegans. *Genetics* **77**, 71–94 (1974)
14. Evans, T. Transformation and microinjection. *WormBook* (2006). doi:10.1895/wormbook.1.108.1
15. Mello, C. & Fire, A. DNA transformation. *Methods Cell Biol.* **48**, 451–482 (1995).
16. Iwano, S., et al. Single-cell bioluminescence imaging of deep tissue in freely moving animals. *Science*, **359**, 935-39, (2018)
17. Savage, C., et al. *mec-7* is a β-tubulin gene required for the production of 15-protofilament microtubules in Caenorhabditis elegans. *Genes and Dev*. **3**, 870-881 (1989)
18. Collins, T. J. ImageJ for microscopy. *Biotechniques*, **43**, 25-30, 2007
19. Kerr, R., Lev-Ram, V., Baird, G., Vincent, P., Tsien, R. Y., & Schafer, W. R. Optical imaging of calcium transients in neurons and pharayngeal muscle of C. *elegans*. *Neuron*. **26**, 583-594 (2000).
20. Scholz, M., Lynch, D. J., Lee, K. S., Levine, E & Biron, D. A scalable method for automatically measuring pharyngeal pumping in C. elegans. *J. Neurosci. Meth.* **274**, 172-178 (2016)
21. Shannon, C. E. A Mathematical theory of communication. *The Bell Systems Technical J,* 27, 379-423 (1948)
22. Gonzalez, R. C., Woods, R. E. & Eddins, S. L. *Digital Image Processing Using MATLAB*. New Jersey, Prentice Hall, Chapter 11 (2003).
23. Li, Q. et al. Structural mechanism of voltage-dependent gating in an isolated voltage-sensing domain. *Nat. Struct. Mol. Biol.* **21**, 244-252 (2014)

**Table S1: Biochemical reactions performed to construct plasmids that express engineered AMBER protein constructs**

| Template | Reagents | Forward (F) and Reverse (R) primers | Reactions | Descriptive Notes | Modified DNA plasmids/fragments |
|---|---|---|---|---|---|
| VSFP2.1 | Quick change site-directed mutagenesis (QCSDM) (Agilent; Catalog # 210518) | F: 5'-gtaccgagctcggagccactagtccagtg-3'<br>R: 5'-cactggactagtggctccgagctcggtac-3 | Substitution mutation | BamHI site flanking *Cerulean* domain. | NI-VSFP-BI |
| pCMV$_{lux}$ | | F: 5'-ccccctgatcccccaccggatcccatatggtatttcttgatgttgtcgt-3'<br>R: 5'-acgacaacatcaagaaataccatatgggatccggtgggggatcaggggg-3' | Substitution mutation | BamHI site flanking *luxB* | luxB-BI |
| luxB-BI | | F: 5'-ctggtagaagttccgcagcacgcggccgctctcgctcaggtcgtgggagc-3'<br>R: 5'-gctcccacgacctgagcgagagcggccgcgtgctgcggaacttctaccag-3' | Substitution mutation | Creating NotI site 270bp upfront of *luxA* inside *luxD.* | 270nt_luxA-luxB-BI |
| NI-VSFP-BI & 270nt_luxA-luxB-BI | BamHI-HF (NEB, R3136S), NotI-HF(NEB, R3189S), 10X CutSmart Buffer | None | Double digestion at BamHI and NotI sites | Double stranded vector and insert fragments | VSFP vector BB and 270nt_luxA-luxB insert |
| VSFP vector backbone and 270nt_luxA-luxB insert | T4 DNA ligase, 10X ligation buffer (NEB, M0202S) | None | Ligation | Ligation of VSFP vector backbone and 270nt_*luxA-luxB* insert | VSD-270nt_luxA-luxB-YFP |
| VSD-270nt_luxA-luxB-YFP | QCSDM (Agilent; Catalog # 210518) | F: 5'-cgtcgatgcggccgcatgaagttcggcaac-3'<br>R: 5’-gttgccgaacttcatgcggccgcatcgacg-3’ | Deletion mutation | Delete the additional 270bp flanking the 5’ of *luxAB* | VSD-luxA-luxB-YFP |
| VSD-luxA-luxB-YFP | Quick change Multi site-directed mutagenesis (QCMSDM), (Agilent; Catalog # 200514) | 5'-tctgcttgtcggcagtgatatagacgttgtggctgttgta-3'<br>5'-cctcgtgaccaccttaggctacggcct-3'<br>5'-aggccgtagcctaaggtggtcacgagg-3'<br>5'-tacaacagccacaacgtctatatcactgccgacaagcaga-3' | Substitution at multiple sites | Modification of *Citrine* to *YFP3* | VSD-luxA-luxB-YFP3 |
| VSD-luxA-luxB-YFP3 | | 5'-tgtggcggatcttgaagttagccttgatgccgttcttctg-3'<br>5'-cggcgagctgcacaccgccgtcctcgatg-3'<br>5'-ctaccagtccgccctgttcaaagaccccaacgag-3' | Substitution at multiple sites | Modification of YFP3 to *YPet* | VSD-luxA-luxB-YPet |

| | | | | | |
|---|---|---|---|---|---|
| VSD-luxA-luxB-YPet | | 5'-ccggcatcagcaacgtctgctgcggcttt-3'<br>5'-aaagccgcagcagacgttgctgatgccgg-3'<br>5'-ccttgatgctgtcgtagtccacgctggtgat-3'<br>5'-atgctggcgacaatctcgtccacggtgccg-3'<br>5'-ctcgttgtcggcttctatgtagccctcggtc-3' | Substitution at multiple sites | Modification of *luxA-luxB* to *eluxA-eluxB* | VSD-eluxA-eluxB-YPet |
| VSD-eluxA-eluxB-YPet | QCSDM (Agilent; Catalog # 210518) | F: 5'-atagcgctgggtgctgcgttttcttcgacatctcc-3'<br>R: 5'-ggagatgtcgaagaaaacgcagcacccagcgctat-3' | Substitution mutation | Mutating Prolines of the T2A to Alanines for creating fusion linker between *luxA* and *luxB* | <u>***VSD-eluxAB-YPet***</u> |
| VSD-eluxAB-YPet | | F: 5'-ctagtccagtgtggtgggatcccttgtacagctcgtcc-3'<br>R: 5'-ggacgagctgtacaagggatcccaccacactggactag-3' | Substitution mutation | Introducing BamHI site at 3' end of YPet | VSD-eluxAB-YPet-Bmod |
| VSD-eluxAB-YPet-Bmod | BamHI-HF (NEB, R3136S), 10X CutSmart Buffer | None | Double digestion at two BamHI sites | Double vector (BI-BB-VSD-eluxAB-BI) and insert (Bi-YPet-BI) fragments | BI-BB-VSD-eluxAB-BI and Bi-YPet-BI with flanking BamHI sites |
| Large fragment (VSD-eluxAB) | T4 DNA ligase, 10X ligation buffer (NEB, M0202S) | None | Ligation | Self ligation of the large fragment | <u>***VSD-eluxAB***</u> |
| VSD-eluxAB | QCSDM (Agilent; Catalog # 210518) | F: 5'-ctcggagccactagtccagtgtggcggccgcacatatggtatttcttgatgttgtc-3'<br>R: 5'-gacaacatcaagaaataccatatgtgcggccgccacactggactagtggctccgag-3' | Substitution mutation | Converting BamHI to the NotI site | VSD-eluxAB-NI |
| VSD-eluxAB-NI | | F: 5'-gcaggaagttgccgaacttcatgtgggatcctcgacgcttgttctgtgatattg-3'<br>R: 5'-caatatcacagaacaagcgtcgaggatcccacatgaagttcggcaacttcctgc-3' | Substitution mutation | Converting 5' NotI flanking eluxAB to BamHI site | VSD-BI-eluxAB-NI |

| | | | | | |
|---|---|---|---|---|---|
| VSD-BI-eluxAB-NI | BamHI-HF (NEB, R3136S), 10X CutSmart Buffer | None | Digestion at the BamHI site | Creating double stranded vector backbone (5’-BI-eluxAB-BB-VSD-BI) | BI-eluxAB-BB-VSD-BI fragment with flanking BamHI sites |
| BI-eluxAB-BB-VSD-BI and BI-YPet-BI fragments | T4 DNA ligase, 10X ligation buffer (NEB, M0202S) | None | Ligation | Ligation of small and large double stranded fragments | ***VSD-YPet-eluxAB*** |
| pCMV$_{lux}$ | QCSDM (Agilent; Catalog # 210518 | F: 5'-gtaaccccggtcctaatttaaatcgcgggtggatg-3'<br>R: 5'-catccacccgcgatttaaattaggaccggggttac-3' | Deletion mutation | Delete the luxA and luxB domains | ***luxCDE-FRP*** |
| luxCDE-FRP | QCSDM (Agilent; Catalog # 210518 | F: 5'-gggcctggccaagagataagctagcagctcgctga-3'<br>R: 5'-ggaggagaatcccggccctgctagcatgaacaacaccatcgaga-3' | Substitution mutation | FRP domain with flanking BmtI sites | luxCDE-BmI-FRP-BmI |
| luxCDE-BmI-FRP-BmI | BmtI-HF (NEB, R3658S), 10X CutSmart Buffer | None | Digestion at BmtI sites | Double stranded fragment of FRP and luxCDE with flanking BmtI sites | BmI-FRP-BmI; BmI-luxCDE-BmI |
| BmI-luxCDE-BmI | T4 DNA ligase, 10X ligation buffer (NEB, M0202S) | None | Ligation | Self ligation of the luxCDE fragment with vector backbone | ***luxCDE*** |
| ***VSD-YPet-eluxAB*** | BmtI-HF (NEB, R3658S), 10X CutSmart Buffer | None | Digestion at BmtI sites | Double stranded fragment of VSD-YPet-eluxAB with flanking BmtI sites | BmI-VSD-YPet-eluxAB-BmI |
| ***VSD-eluxAB-YPet*** | BmtI-HF (NEB, R3658S), 10X CutSmart Buffer | None | Digestion at BmtI sites | Double stranded fragment of VSD-eluxAB-YPet with flanking BmtI sites | BmI-VSD-eluxAB-YPet-BmI |
| BmI-VSD-YPet-eluxAB-BmI and BmI-FRP-BmI | T4 DNA ligase, 10X ligation buffer (NEB, M0202S) | None | Ligation | Ligation of vector and insert | ***FRP-VSD-YPet-eluxAB*** |

| | | | | | |
|---|---|---|---|---|---|
| BmI-VSD-eluxAB-YPet-BmI and BmI-FRP-BmI | T4 DNA ligase, 10X ligation buffer (NEB, M0202S) | None | Ligation | Ligation of vector and insert | ***FRP-VSD-eluxAB-YPet*** |
| $pCMV_{lux}$ | QCSDM (Agilent; Catalog # 210518) | F: 5'-atagcgctgggtgctgcgttttcttcgacatctcc-3'<br>R: 5'-ggagatgtcgaagaaaacgcagcacccagcgctat-3' | Substitution mutation | Converting T2A between *luxA* and *luxB* into a fusion linker | ***luxAB*** |
| MBP-rTRPV1 | Phusion HF DNA polymerase (NEB; M0530L) and DpnI (NEB, R0176S) | F: 5'-ttaaacttaagcttggtaccgaacaacgggctagct-3'<br>R: 5'-gatatctgcagaattcttatttctcccctgggaccatggaa-3' | PCR amplification of rTRPV1 and DpnI treatment | Temperature gradient PCR with $T_m$ ranging 66-72C | Amplified gene product of *rTRPV1* |
| pCDNA3.1(+) | EcoRI-HF (NEB; R3101S), KpnI-HF (NEB; R3142L), 10X CutSmart Buffer | None | Double Digestion | pCDNA3.1(+) vector BB with flanking restriction sites | KpnI-pCDNA3.1-EcoRI |
| Amplified gene product of *rTRPV1* and KpnI-pCDNA3.1-EcoRI | T4 DNA ligase, 10X ligation buffer (NEB, M0202S) | None | Ligation | Ligation of vector and insert | ***pCDNA3.1-rTRPV1*** |
| *FRP-VSD-eluxAB-YPet* and synthetic *YPet* DNA coding *DK* | In-Fusion enzyme mix (TaKaRa Bio Inc; Catalog # 639649) | None | Ligation | Ligation of vector and insert | ***DK*** Mutant |
| L3691 | EcoRI-HF(NEB; R3101S) and KpnI-HF(NEB; R3142L), 10X CutSmart Buffer | None | Ligation | L3691 vector BB with flanking restriction sites | KpnI-L3691-EcoRI |
| *FRP-VSD-eluxAB-YPet* | Phusion HF DNA polymerase (NEB; M0530L) and DpnI (NEB, R0176S) | F: 5'-tagtgagtcgtattggtaccatgaacaacaccatcgagac-3'<br>R: 5'-gccggctagcgaattccttgtacagctcgtccat-3' | PCR amplification and DpnI treatment | Temperature gradient PCR with $T_m$ ranging 66-72C; 3%DMSO | Amplified gene product of *FRP-VSD-eluxAB-YPet* |

| | | | | | |
|---|---|---|---|---|---|
| Amplified *FRP-VSD-eluxAB-YPet* and KpnI-L3691-EcoRI | In-Fusion enzyme mix (TaKaRa Bio Inc; Catalog # 639649) | None | Ligation | Ligation of vector and insert | ***mec7-Btp*** |
| *luxCDE* | Phusion HF DNA polymerase (NEB; M0530L) and DpnI (NEB, R0176S) | F: 5'-tagtgagtcgtattggtaccatgggcaccaagaaga-3'<br>R: 5'-ccggctagcgaattctcagcgagcttctagagggc-3' | PCR amplification of cDNA and DpnI treatment | Temperature gradient PCR with $T_m$ ranging 66-72C; +0.5mM $MgCl_2$ | Amplified gene product of *luxCDE* |
| Amplified gene product of *luxCDE* and KpnI-L3691-EcoRI | In-Fusion enzyme mix (TaKaRa Bio Inc; Catalog # 639649) | None | Ligation | Ligation of vector and insert | ***mec7-Bts*** |
| *mec7-Btp and L3790* | SphI-HF (NEB, R3182S) and ClaI(NEB; R0197S), 10X CutSmart Buffer | None | Restriction digestion | Digestion and purification of vector and insert | mec7-Btp BB and SphI-myo-2-ClaI fragments |
| mec7-Btp BB and SphI-myo-2-ClaI fragments | T4 DNA ligase, 10X ligation buffer (NEB, M0202S) | None | Ligation | Ligation of vector and insert | ***myo2-Btp*** |
| *mec7-Bts* | AscI (NEB, R0558S) and BamHi-HF(NEB, R3658S), 10X CutSmart Buffer | None | Restriction digestion | Digestion and purification of vector | mec7-Bts BB |
| mec7-Bts BB and Synthetic myo-2 promoter | In-Fusion enzyme mix (TaKaRa Bio Inc; Catalog # 639649) | None | Ligation | Ligation of vector and insert | ***myo2-Bts*** |

**Table S2: Information entropy ratio, IR of different engineered AMBER protein constructs**

| No. | Constructs | Substrate expressed | % of Information transmitted |
|---|---|---|---|
| 1 | FRP-VSD-eluxAB-YPet | Yes | 85.4 |
| 2 | Dark Mutant | Yes | 73.2 |
| 3 | VSD-eluxAB-YPet | Yes | 40.25 |
| 4 | VSD-YPet-eluxAB | Yes | 35.15 |
| 5 | FRP-VSD-YPet-eluxAB | Yes | 19.25 |
| 6 | VSD-eluxAB | Yes | - |
| 7 | FRP-VSD-eluxAB-YPet | No | - |
| 8 | VSD-eluxAB-YPet | No | 28.2 |
| 9 | VSD-YPet-eluxAB | No | 28.73 |
| 10 | FRP-VSD-YPet-eluxAB | No | - |
| 11 | VSD-eluxAB | No | 9.7 |
| 12 | Untransfected | | 0.56 |

**Table S3: Paired single tailed Student's t-test statistic of bioluminescent intensities of various AMBER protein constructs expressed in HEK293 cells.**

| No. | Constructs | Substrate expressed | t-test score | No. of cells, n | *p*-value |
|---|---|---|---|---|---|
| 1 | FRP-VSD-eluxAB-YPet | Yes | 57.38 | 53 | < 0.00001 |
| 2 | VSD-eluxAB-YPet | Yes | 50.60 | 47 | |
| 3 | FRP-VSD-YPet-eluxAB | Yes | 67.82 | 77 | |
| 4 | VSD-YPet-eluxAB | Yes | 49.57 | 53 | |
| 5 | VSD-eluxAB-YPet | No | 89.85 | 160 | |
| 6 | VSD-YPet-eluxAB | No | 57.72 | 69 | |
| 7 | VSD-eLuxAB | No | 56.27 | 60 | |
| 8 | Dark mutant | Yes | 65.27 | 83 | |

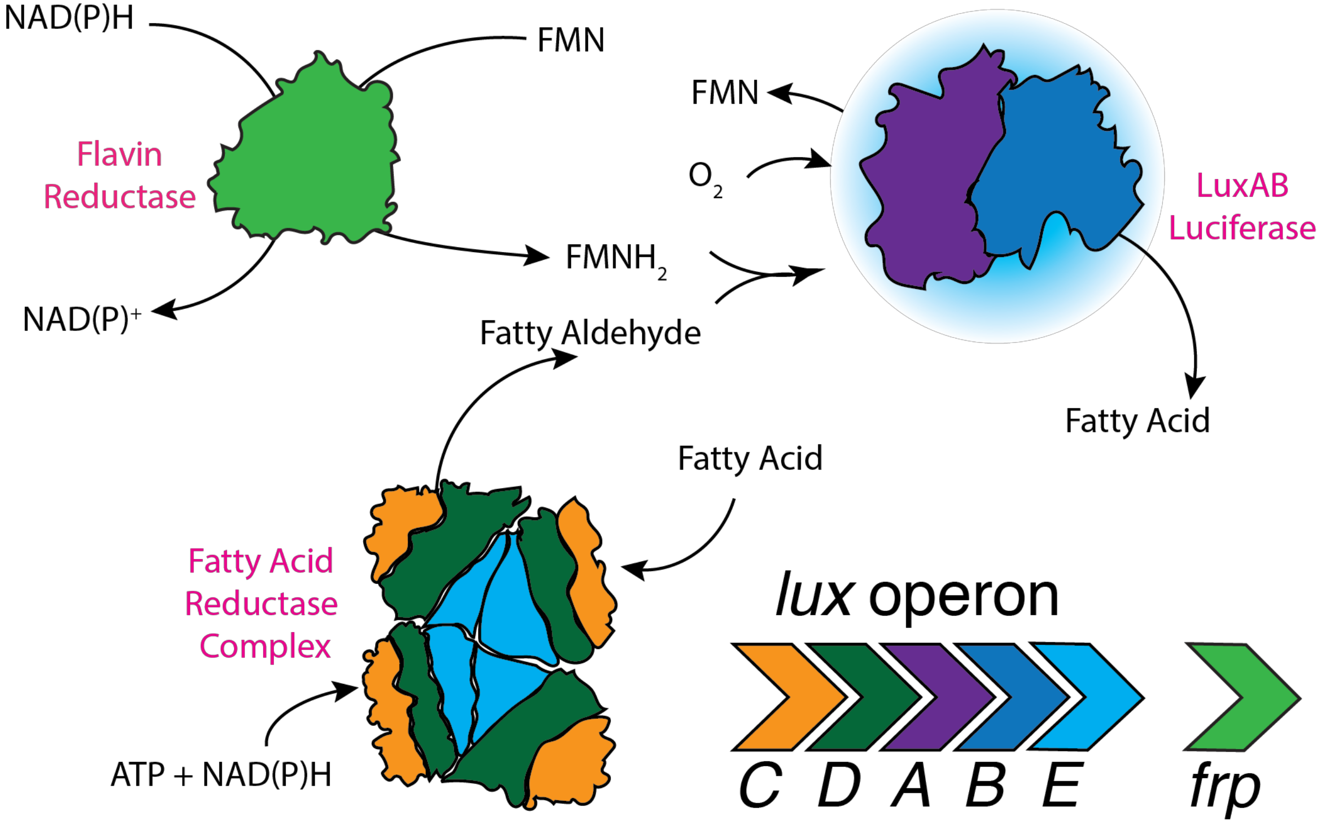


**Figure S1**:Molecular mechanisms of bacterial bioluminescence

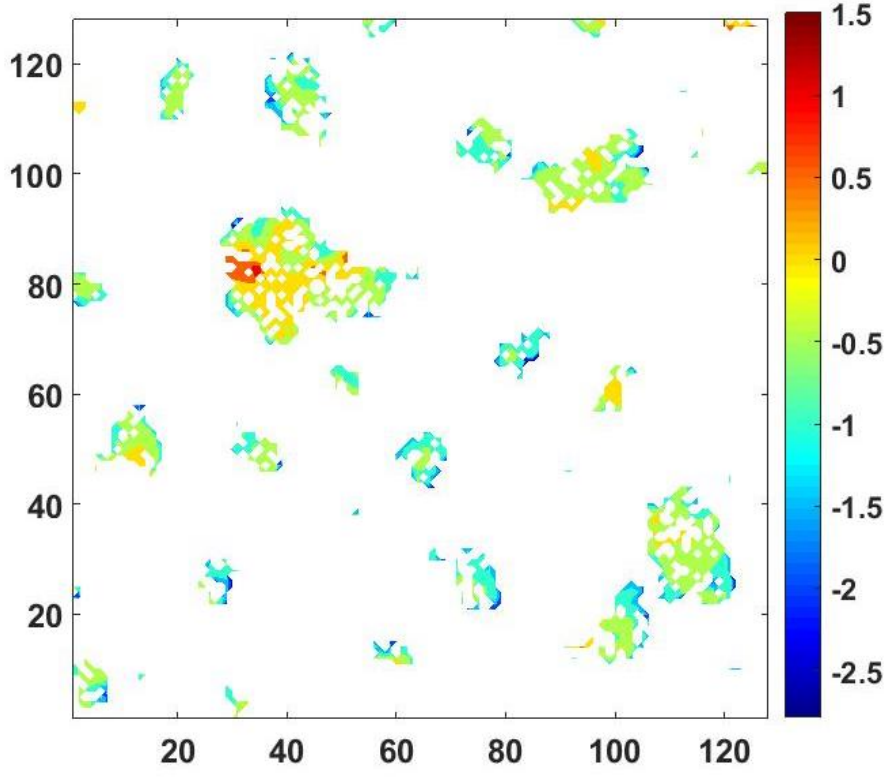


**Figure S2**: Fractional bioluminescent intensity change computed using MATLAB programming in a population of HEK293 cells expressing the bright probe and its substrate. The abscissa and ordinate values correspond to the pixel count (128 x 128). Color-coded contour values represent $\log_{10}(\Delta L/L)$ of individual pixels from a population of cells within the chosen field of view. The maximum and average values of the fractional bioluminescence of pixels are $(\Delta L/L)_{max}$~28 and $(\Delta L/L)_{avg}$ ~ 0.76 respectively. About 85% of information coded in the bright field image was retrieved from the emitted biophotons after KCl addition. Paired single tailed Student's t-test hypothesis confirmed a significant difference in the mean bioluminescent intensity of the chosen cells (t-test statisic ~ 57.8, n= 53 cells) after KCl addition indicating that alternate hypothesis is met at $\alpha$ > 95% confidence ($p < 0.00001$).

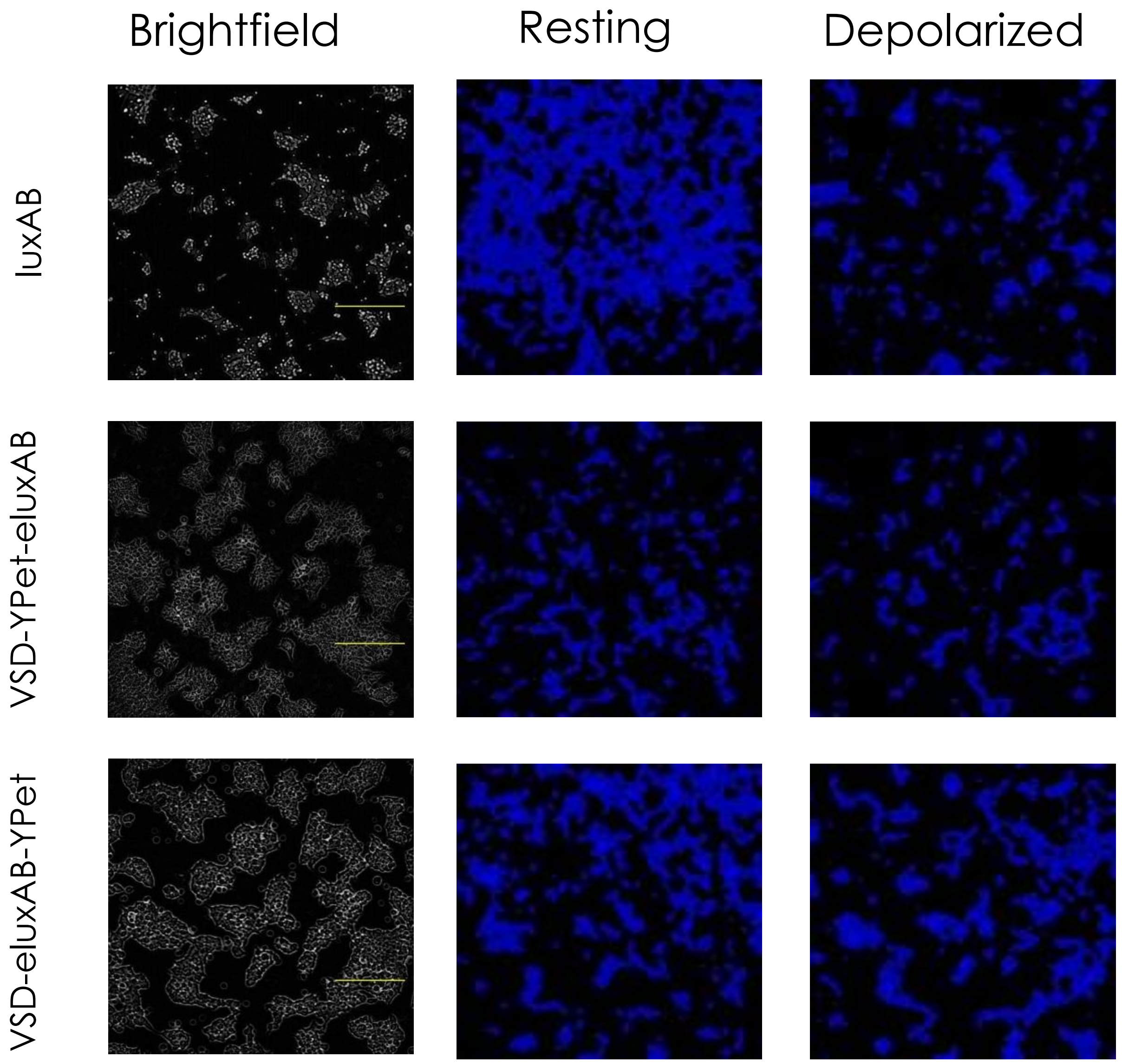


**Fig S3**: Endogenous substrates produce weak bioluminescent signal. Bioluminescent signals were 10 times smaller than the signals obtained when the substrate producing proteins were overexpressed. (Top) In particular, cytosolic expression of lux operon (with a fused luxAB) produced a weak bioluminescence requiring 30s integration for a comparable brightness to other membrane targeted constructs (Middle and Bottom) imaged with 10s integration. See Table S3 for the statistical analysis. The length of the scale bar is 250μm.

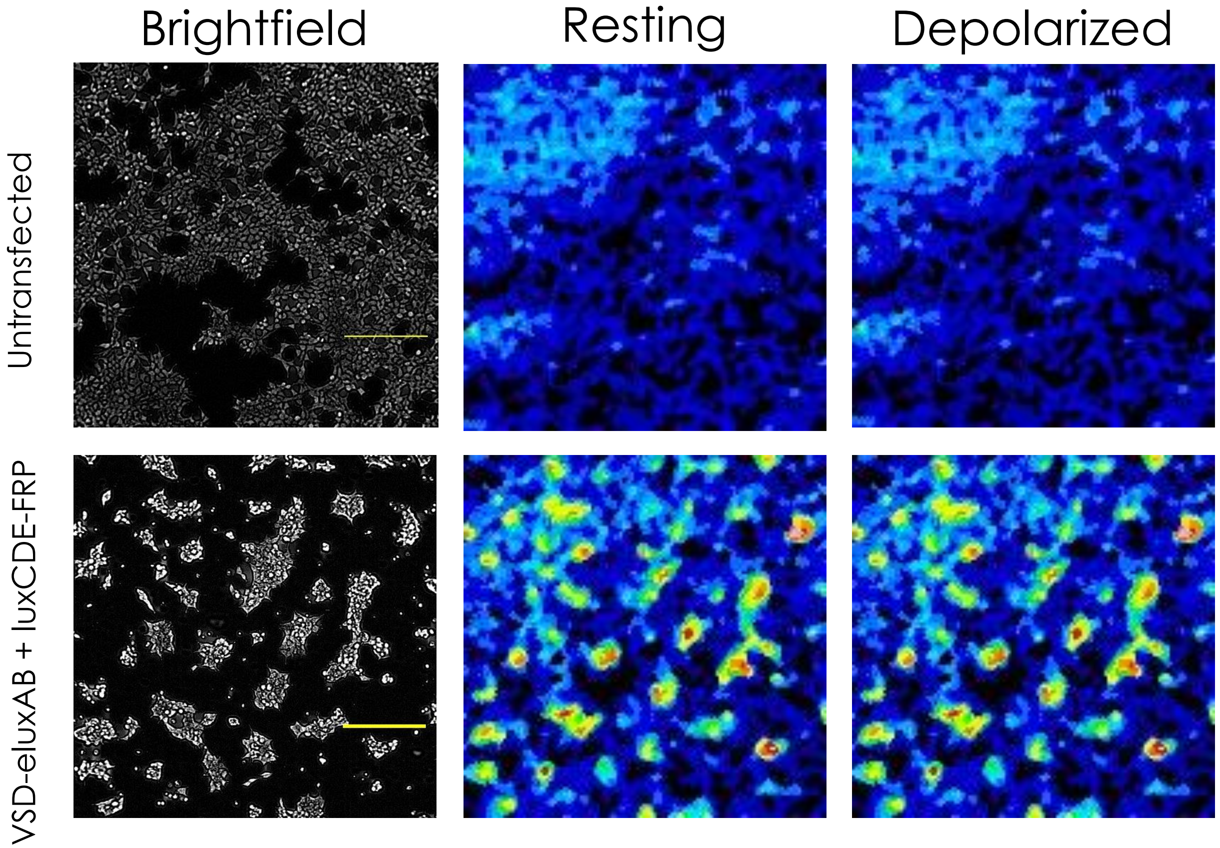


**Figure S4**: (a) Background light emission from the untransfected cells which, does not change after KCl challenge. (b) KCl induced depolarization did not show appreciable intensity change for VSD-eluxAB co-expressed with its substrate producing protein complex. The length of the scale bar is 250μm.

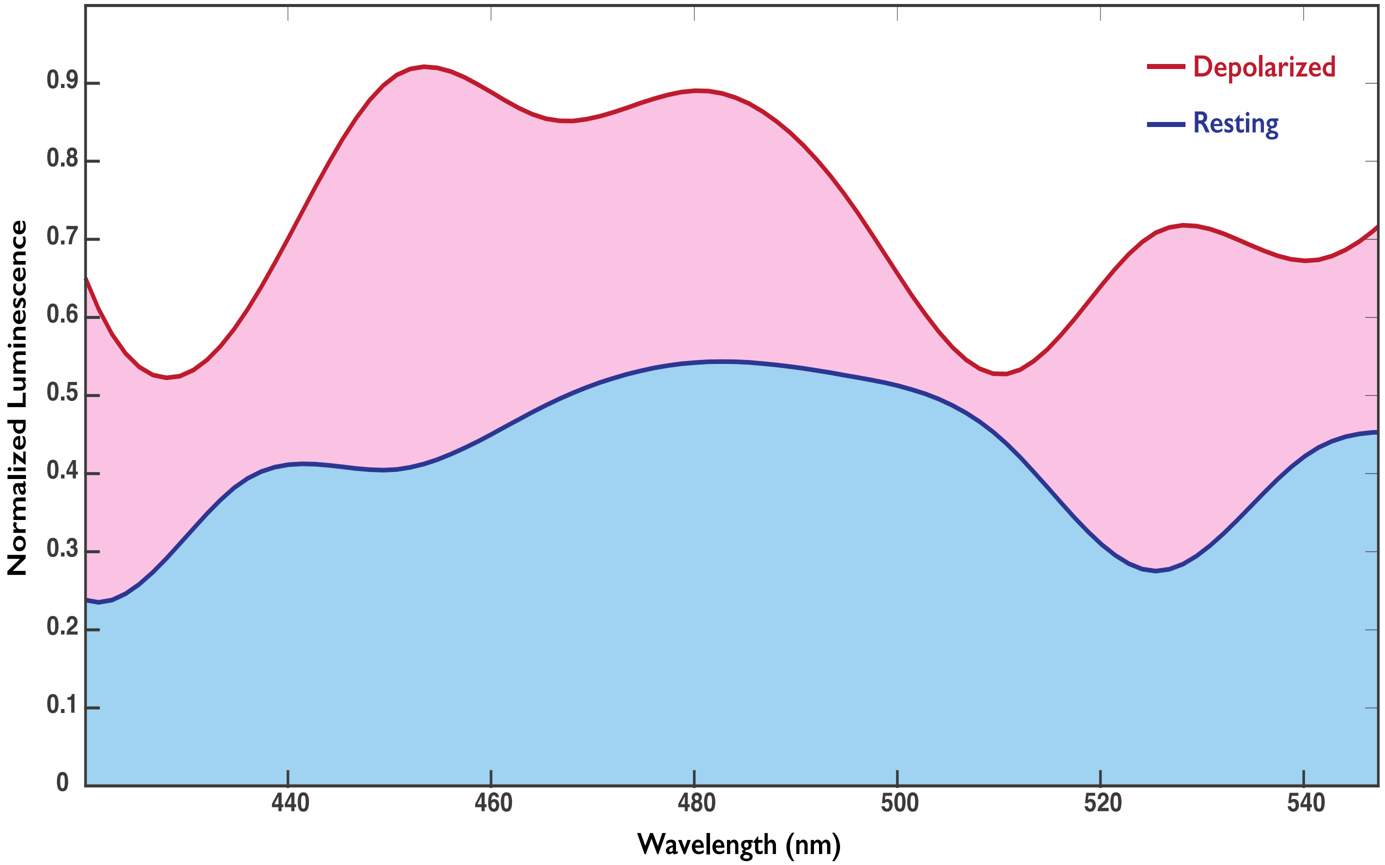


**Figure S5**: Bioluminescent emission spectra of HEK293 cells expressing the dark mutant probe and its substrate before and KCl challenge obtained from plate reader experiments. An increase in the intensity at λ= 490nm suggest that YPet barrel structure (with mutated chromophore) is sufficient to modulate the enzymatic activity of the membrane targeted bright probe. Spectral intensity at different wavelengths are the mean photon counts from n=10 samples. Representative emission spectra was obtained by curve fitting using 2-point moving average method.

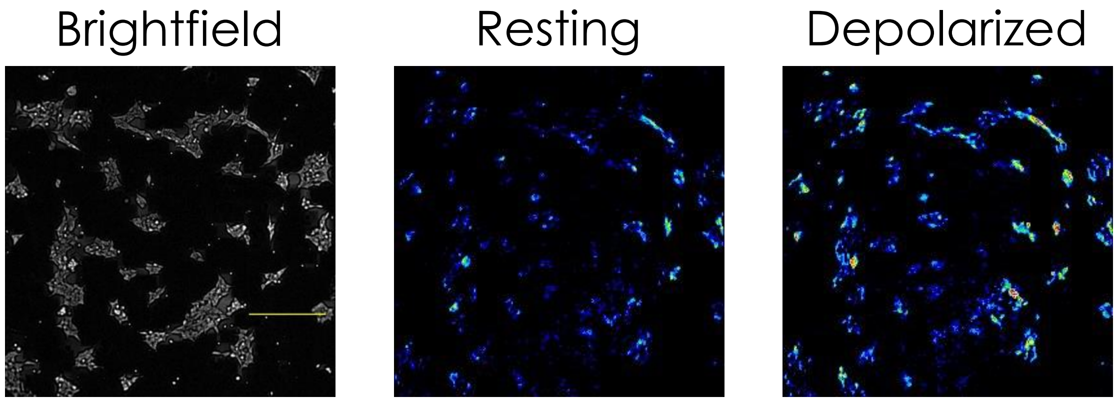


**Figure S6**: Increase in the endogenous NADH fluorescence of HEK293 cells co-expressing the bright probe and its substrate after KCl challenge. The length of the scale bar is 250μm.

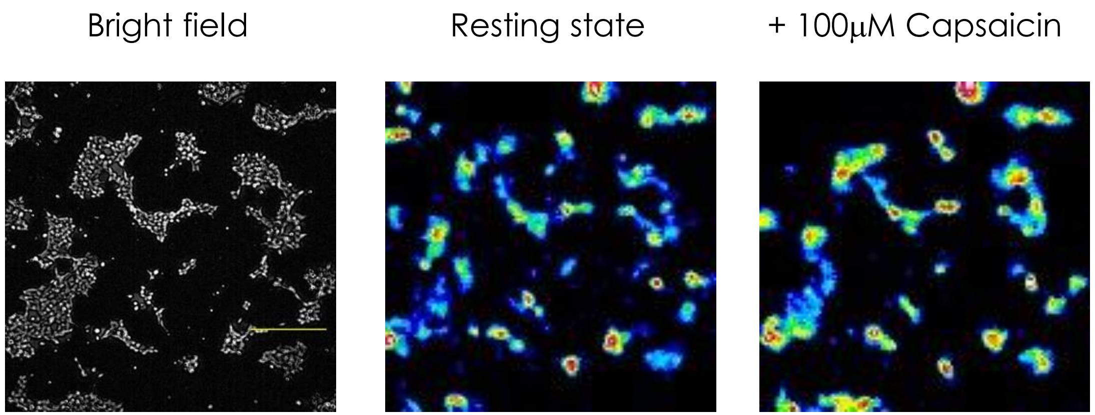


**Figure S7**: Bioluminescence produced by the HEK293 cells co-expressing the bright probe with luxCDE and rTRPV1 after addition of Capsaicin. The hydrophobic property of the Capsaicin necessitate a small assay volume (about 400$\mu$ l) to achieve equilibration at the room temperature when added at 100$\mu$M final concentration. An increase in the bioluminescence after activation of rTRPV1 with Capsaicin is very evident although the fractional change is comparatively smaller than the KCl induced depolarization. The length of the scale bar is 250μm.

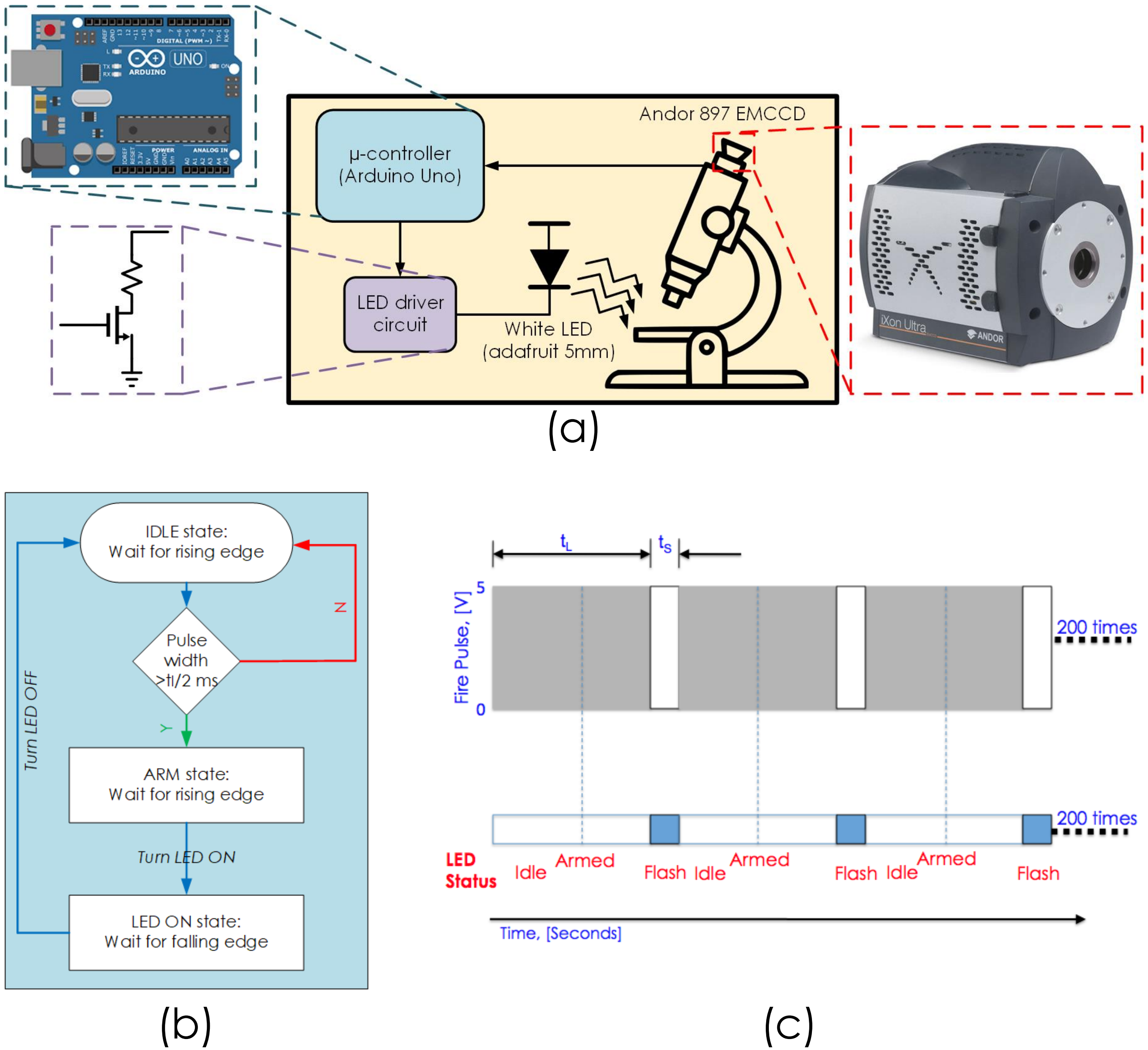


**Figure S8**: Custom-built imaging set up to track the worm position during bioluminescent recording (a) Schematic of the experimental hardware used for tracking the worm position; (b) Algorithm used for programming the Arduino Uno microcontroller that drives the flash circuit based on the trigger output of the EMCCD camera; (c) Representative LED lighting and camera exposure patterns that allows live tracking during bioluminescent recording.

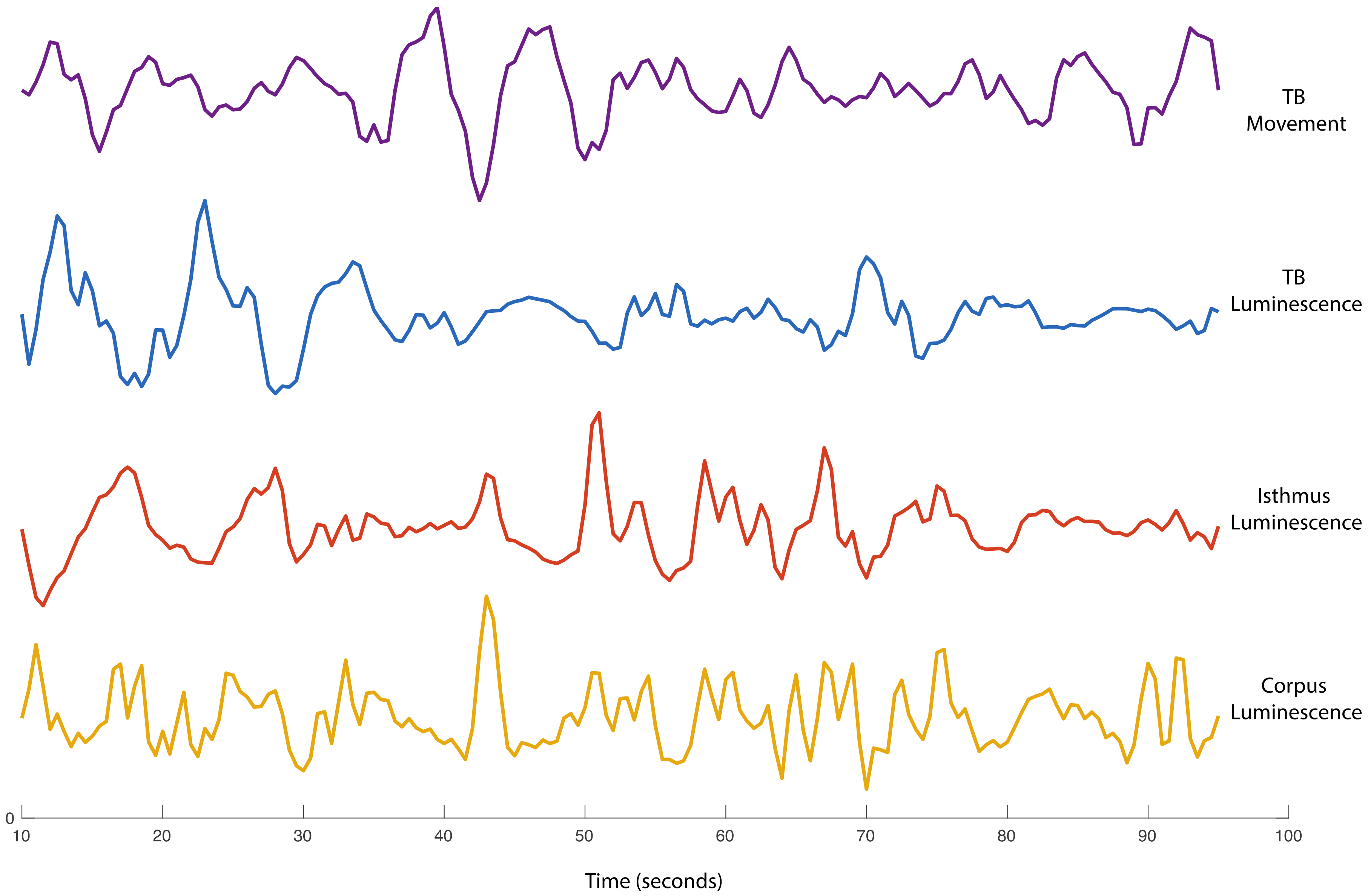


**Figure S9:** Brightfield (top trace) and luminescence data (bottom traces) collected from the pharyngeal muscle in a second worm. The agreement between TB movement and its luminescence is clearly noticeable between 30-60 second window.

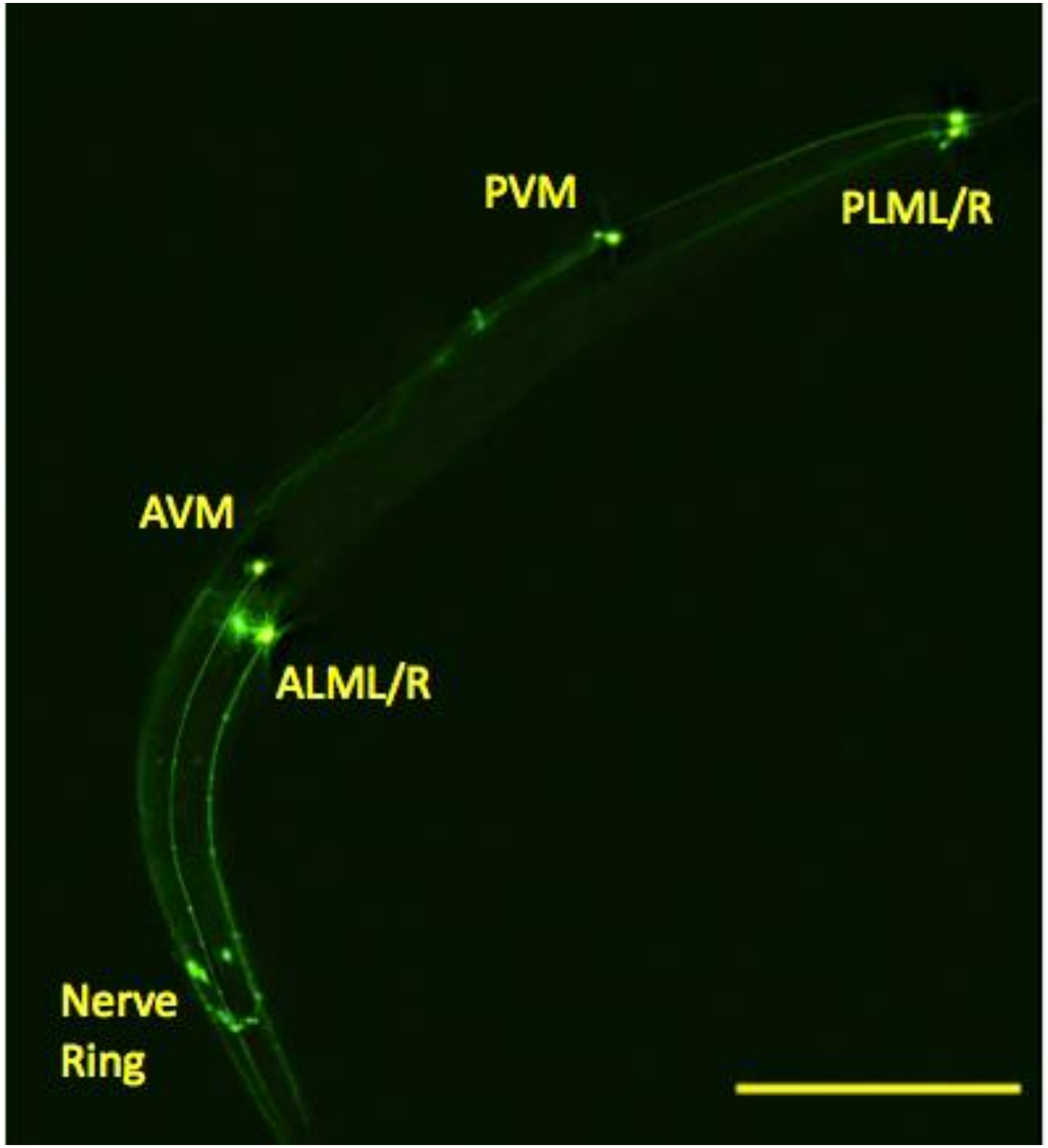


**Figure S10**: Mapping of mechanosensory neuronal circuit using eGFP fluorescence. The scale bar length is 250$\mu$m.

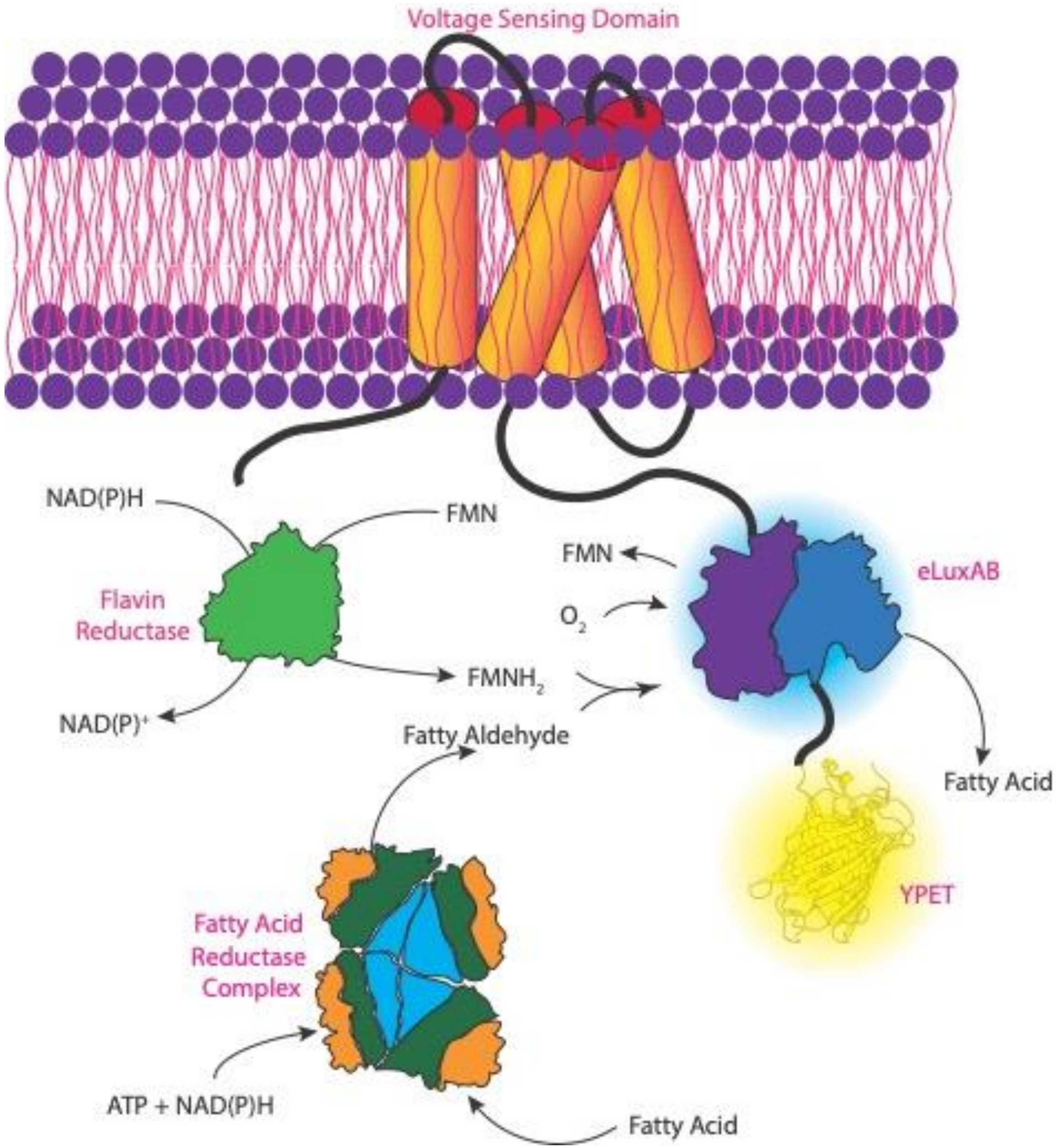


**Figure S11**: Schematic illustration of the light reaction using VSD-eLuxAB-YPet co-expressed with the LuxCDE-FRP

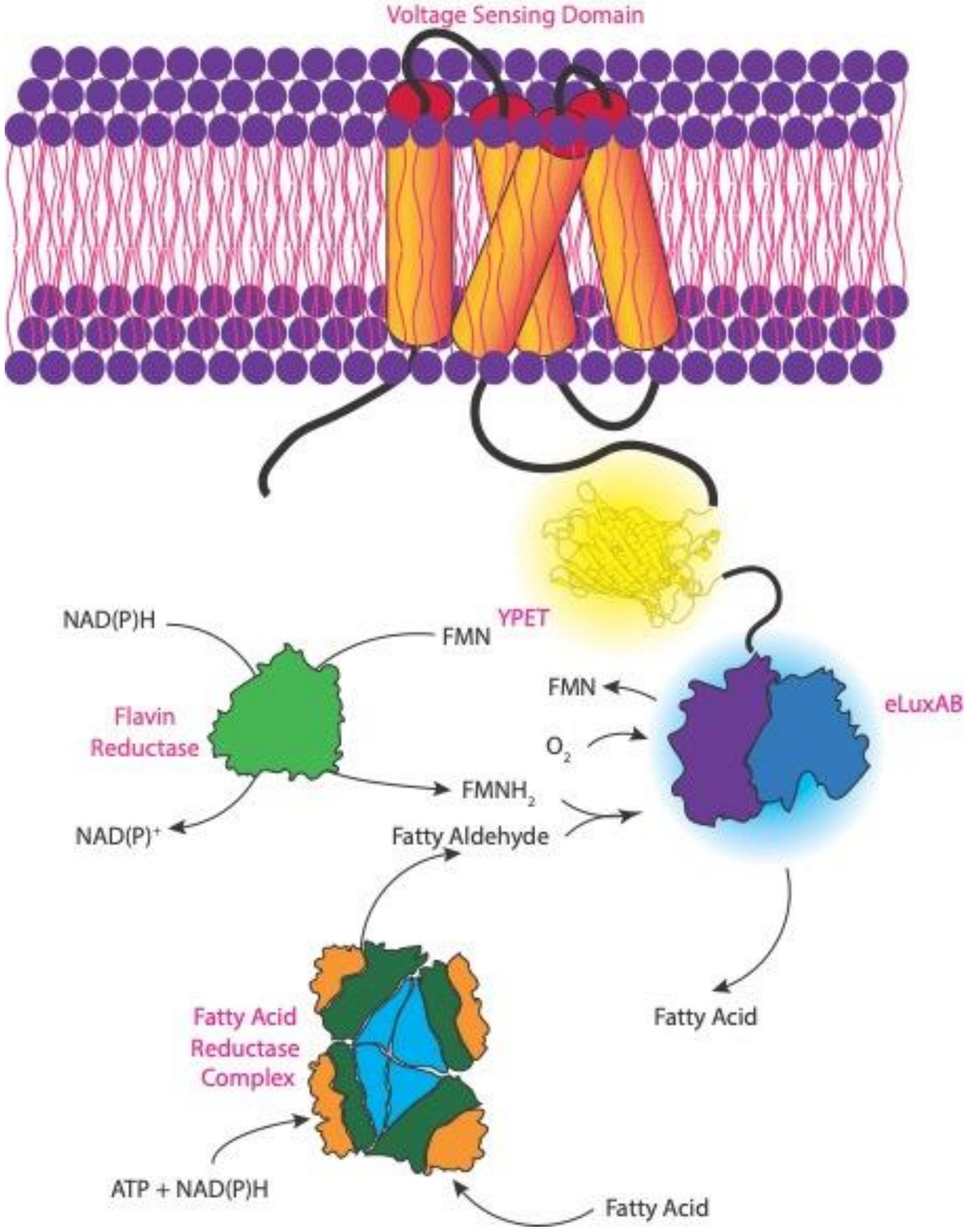


**Figure S12**: Schematic illustration of the light reaction using VSD-YPet-eluxAB co-expressed with LuxCDE-FRP

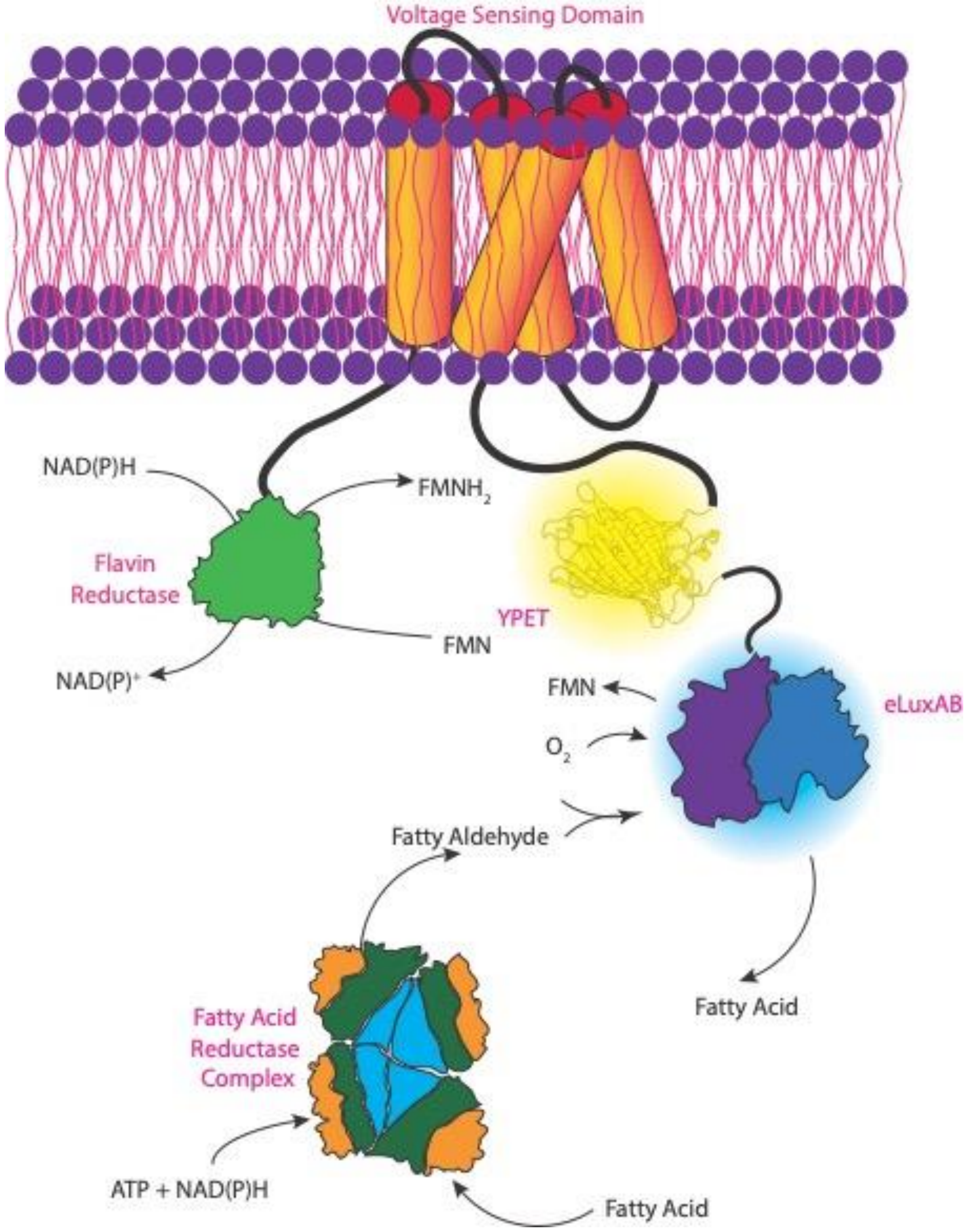


**Figure S13**: Schematic illustration of the light reaction using FRP-VSD-YPet-eluxAB co-expressed with LuxCDE

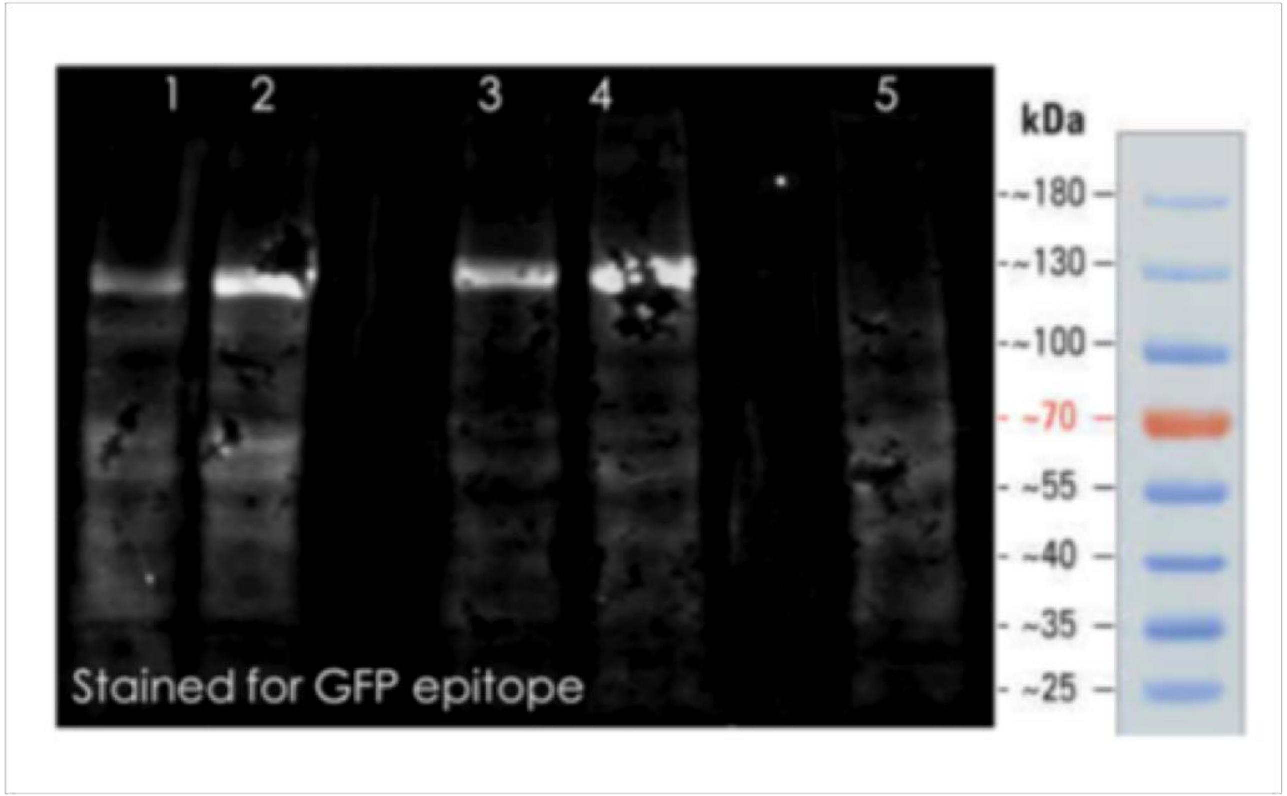


**Figure S14:** Estimation of the molecular weight of engineered AMBER constructs using Western blot. An epitope conserved across all the GFP variants was used to stain the YPet. (Lane 1) VSD-YPet-eluxAB, (Lane 2) VSD-YPet-eluxAB + luxCDE, (Lane 3) VSD-eluxAB-YPet, (Lane 4) VSD-eluxAB-YPet + luxCDE and (Lane 5) HEK293 cells. The estimated molecular weight of the probes are ~ 130 KDa. This estimation agrees closely with a predicted value of ~ 133 KDa assuming an average molecular weight of an amino acid residue of 110g/mole for a total of 1213 residues.

**Legends for Supplementary Movies**

Movie **SM1**: Voltage signals of *C. elegans* pharyngeal muscles recorded using AMBER during natural feeding behavior.

Movie **SM2**: Bursts of mechanosensitive neuronal activities of *C. elegans* recorded using AMBER during frequent reversals.

Movie **SM3**: A cartoon movie showing activities of *C. elegans* mechanosensory neural circuit during frequent reversals.

Movie **SM4**: Mechanosensory neuronal activities of multiple animals during collision recorded using AMBER.
